# Standing genetic variation and chromosomal rearrangements facilitate local adaptation in a marine fish

**DOI:** 10.1101/782201

**Authors:** Hugo Cayuela, Quentin Rougemont, Martin Laporte, Claire Mérot, Eric Normandeau, Yann Dorant, Ole K. Tørresen, Siv Nam Khang Hoff, Sissel Jentoft, Pascal Sirois, Martin Castonguay, Teunis Jansen, Kim Praebel, Marie Clément, Louis Bernatchez

## Abstract

Population genetic theory states that adaptation most frequently occurs from standing genetic variation, which results from the interplay between different evolutionary processes including mutation, chromosomal rearrangements, drift, gene flow and selection. To date, empirical work focusing on the contribution of standing genetic variation to local adaptation in the presence of high gene flow has been limited to a restricted number of study systems. Marine organisms are excellent biological models to address this issue since many species have to cope with variable environmental conditions acting as selective agents despite high dispersal abilities. In this study, we examined how, demographic history, standing genetic variation linked to chromosomal rearrangements and shared polymorphism among glacial lineages contribute to local adaptation to environmental conditions in the marine fish, the capelin (*Mallotus villosus*). We used a comprehensive dataset of genome-wide single nucleotide polymorphisms (25,904 filtered SNPs) genotyped in 1,359 individuals collected from 31 spawning sites in the northwest Atlantic (North America and Greenland waters). First, we reconstructed the history of divergence among three glacial lineages and showed that they diverged from 3.8 to 1.8 MyA. Depending on the pair of lineages considered, historical demographic modelling provided evidence for divergence with gene flow and secondary contacts, shaped by barriers to gene flow and linked selection. We next identified candidate loci associated with reproductive isolation of these lineages. Given the absence of physical or geographic barriers, we thus propose that these lineages may represent three cryptic species of capelin. Within each of these, our analyses provided evidence for large *N_e_* and high gene flow at both historical and contemporary time scales among spawning sites. Furthermore, we detected a polymorphic chromosomal rearrangement leading to the coexistence of three haplogroups within the Northwest Atlantic lineage, but absent in the other two clades. Genotype-environment associations revealed molecular signatures of local adaptation to environmental conditions prevailing at spawning sites. Altogether, our study shows that standing genetic variation associated with both chromosomal rearrangements and ancestral polymorphism contribute to local adaptation in the presence of high gene flow.

## Introduction

Environmental conditions experienced by organisms are highly variable over space and time, making adaptation a central mechanism for their persistence. Population genetic theory states that adaptation can occur both from new mutations and from standing genetic variation (Barrett & Schluter 2008, Tigano & Friesen 2016, Schlötterer et al. 2016). In the first case, new mutations appear at very low frequency in the population and can increase and become fixed if they are beneficial under the given environmental conditions. In the second case, alleles that were previously neutral or deleterious can become beneficial following an environmental change and thus, increase in frequencies (Hermisson & Pennings 2005, Prezeworski et al. 2005). Standing genetic variation is maintained in the population through a balance of recurrent mutation, drift, gene flow, and selection (Hermisson & Pennings 2005, Messer & Petrov 2013). Understanding the complex interplay between those mechanisms and their consequences for the evolutionary potential of organisms facing global changes and other anthropogenic pressures (e.g., climate change, ocean pollution, fisheries) is thus a critical challenge for evolutionists and conservation biologists (Davis & Shaw 2001, Hoffmann & Sgro 2011, Franks & Hoffmann 2012).

Theory also states that adaptation from standing genetic variation appears more frequently than adaptation from *de novo* mutations. Indeed, the probability of allele fixation increases with the magnitude of the beneficial effect and the effective population size (*N_e_*), and this probability is higher when the allele has a high initial frequency (*i.e.*, results from the standing genetic variation; Barrett & Schluter 2008, Hedrick 2013). Hence, standing genetic variation allows for faster adaptation than *de novo* mutations, as beneficial alleles are already present in the population (Barrett & Schluter 2008, Hedrick 2013). Accordingly, adaptation from standing genetic variation should dominate in most natural cases, especially at short time scale in species with relatively high genetic diversity (Bernatchez 2016).

Standing genetic variation is preserved by the interplay of recurrent mutation and genetic drift that maintain the neutral and (slightly) deleterious alleles at a frequency higher than 1/2Ne (Barrett & Schluter 2008). Nevertheless, other evolutionary mechanisms can also play a role on the conservation of genetic polymorphism. In particular, gene flow between populations experiencing contrasting environmental conditions can contribute to maintain a high level of standing genetic variation (Hedrick 2013; Tigano & Friesen 2016). In addition, interbreeding and introgression (adaptive or not) among genetically divergent lineages or related species can also preserve or increase standing genetic variation (Hedrick 2013, Tigano & Friesen 2016). Introgressed alleles may have lower initial frequency than standing genetic variants, which reduces their chance of fixation if the hybridization frequency is low and/or if F1 hybrids and backcrosses have reduced fitness (Hedrick 2013). However, favourable alleles can quickly cross species boundaries and increase in frequency (Barton 1979), resulting in adaptive introgression, as increasingly documented in a wide range of species (Hedrick 2013, Suarez-Gonzalez et al. 2018).

Chromosomal rearrangements, which involve change in the structure of the chromosome (e.g., inversion, fusion, fission, and translocation), may also contribute to preserving standing genetic variation within a population or a species (Wellenreuther & Bernatchez 2018, Faria et al. 2019, Wellenreuther et al. 2019). Indeed, low recombination within chromosomal rearrangements may lead to an independent evolution of the genomic regions affected (Faria & Navarro 2010, Wellenreuther et al. 2019). Beyond their contribution to standing genetic variation, rearrangements play a central role in adaptation, especially in the presence of high gene flow (Tigano & Friesen 2016). For instance, chromosomal rearrangements as inversions are increasingly found involved in the processes of phenotypic divergence and local adaptation (Berg et al. 2017, Mérot et al. 2018, Westram et al. 2018, Wellband et al. 2019).

To date, empirical work focusing on the contribution of genomic background (e.g., standing genetic variation and chromosomal rearrangement) to adaptive variation in the presence of high gene flow have been limited to a restricted number of study systems (e.g., Tine et al. 2014, Bernatchez 2016, Le Moan et al. 2019, Barth et al. 2017, Petterson et al. 2019). Marine organisms are excellent biological models to address this issue since many marine species experience highly heterogeneous marine conditions potentially acting as selective agents, yet displaying very weak genetic differentiation due to large *N_e_* and high connectivity associated with strong dispersal capacities (Palumbi 1992, Bradbury et al. 2008, Laporte et al. 2016, Selkoe et al. 2016, Xuereb et al. 2018). Capelin (*Mallotus villosus*) is a key-forage fish species (Buren et al. 2014) that display such characteristics typical of marine organisms. It has high genetic diversity and weak local genetic structure, as previously documented using mtDNA, microsatellite and AFLP markers that suggests large *N_e_* and high gene flow (Præbel et al. 2008, Colbeck et al. 2011). At a larger scale, three genetically-divergent, and parapatric glacial lineages have been reported in the North Atlantic and Arctic seas without apparent physical or geographic barriers separating them (Dodson et al. 2007). Within each lineage, capelin experiences spatiotemporally variable sea conditions that could be important selective drivers (Carscadden et al. 2001, Rose 2005). Following the pre-nuptial migration, adults actively choose to reproduce at sites located in a continuum spanning from the intertidal (i.e., beach spawning sites) to the benthic zone from 1 to 280 m (i.e., demersal-spawning sites) that strongly differ in terms of biotic and abiotic conditions (Nakashima & Wheeler 2002). Environmental variation is especially high in the intertidal zone where the survival and/or development of embryos and larvae depends on temperature, salinity, and trophic productivity (Frank & Leggett 1981a, 1981b, 1982; Leggett et al. 1984, Præbel et al. 2013, Purchase 2018). The position along the gradient of water depth as well as the sea conditions prevailing in the intertidal zone, are expected to exert strong selective pressure on local populations. This offers a suitable system to investigate connectivity and adaptation at different scales in a marine ecosystem, between and within lineages. Yet, the studies that have previously focused on inter- and intra-lineage genetic variation in the capelin were undertaken before the ‘genomic era’, which hampered the investigation of complex scenarios of divergence history and the search of molecular signals (including chromosomal rearrangements) associated with adaptation to local sea conditions.

In this study, we examined how standing genetic variation maintained by chromosomal rearrangements and shared polymorphism among the three aforementioned glacial lineages underlies local adaptation to sea conditions prevailing in the spawning sites of capelin. We used a comprehensive dataset of genome-wide single nucleotide polymorphism (SNP) from 1,310 fish collected at 34 spawning sites throughout the Northwest Atlantic and Arctic waters. First, we confirmed the existence of the three glacial lineages (Northwest Atlantic lineage, NWA; Greenland lineage, GRE; and Arctic lineage, ARC) identified by Dodson et al. (2007) by analyzing the pattern of genetic diversity and differentiation in the whole study area (**Fig.1**). Then, we inferred the demographic history of the three lineages using joint Site Frequency Spectrum (jSFS) for each pair of lineages. We first statistically compared four alternative modes of divergence: Strict Isolation (SI), Isolation-with-Migration (IM), Ancient Migration (AM), and Secondary Contact (SC). Importantly, we took into account the confounding effect of barrier to gene flow, that affects the rate of migration (Barton & Bengsson, 1986) and the confounding effect of linked selection (due to background selection and hitchhiking of linked neutral alleles) that can be approximated as a reduction in local effective population size, *N_e_* (Charlesworth et al. 1993). We also quantified demographic parameters of interest, including migration rate (m), *Ne,* and divergence time). Then, we analyzed in detail the genetic structure within the NWA lineage (**Fig.1**). In particular, we detected the molecular signature of putative chromosomal rearrangements (absent in the other glacial lineages) that best fit a pattern of chromosomal fusion. We examined the molecular signature of local adaption by identifying outlier loci putatively associated with the spawning site position along the water depth gradient (three categories of sites: beach spawning sites, demersal shallow-water sites, and demersal deep-water sites) and environmental variables in beach spawning sites including temperature and trophic production. We also evaluated the contribution of standing genetic variation to local adaptation by testing whether allele sharing of the candidate loci resulted from low rate of gene flow or shared ancestral polymorphism. Finally, we investigated the role of the genomic architecture on local adaptive variation by determining the proportion of candidate loci captured by the putative chromosomal arrangement.

**Fig. 1.**
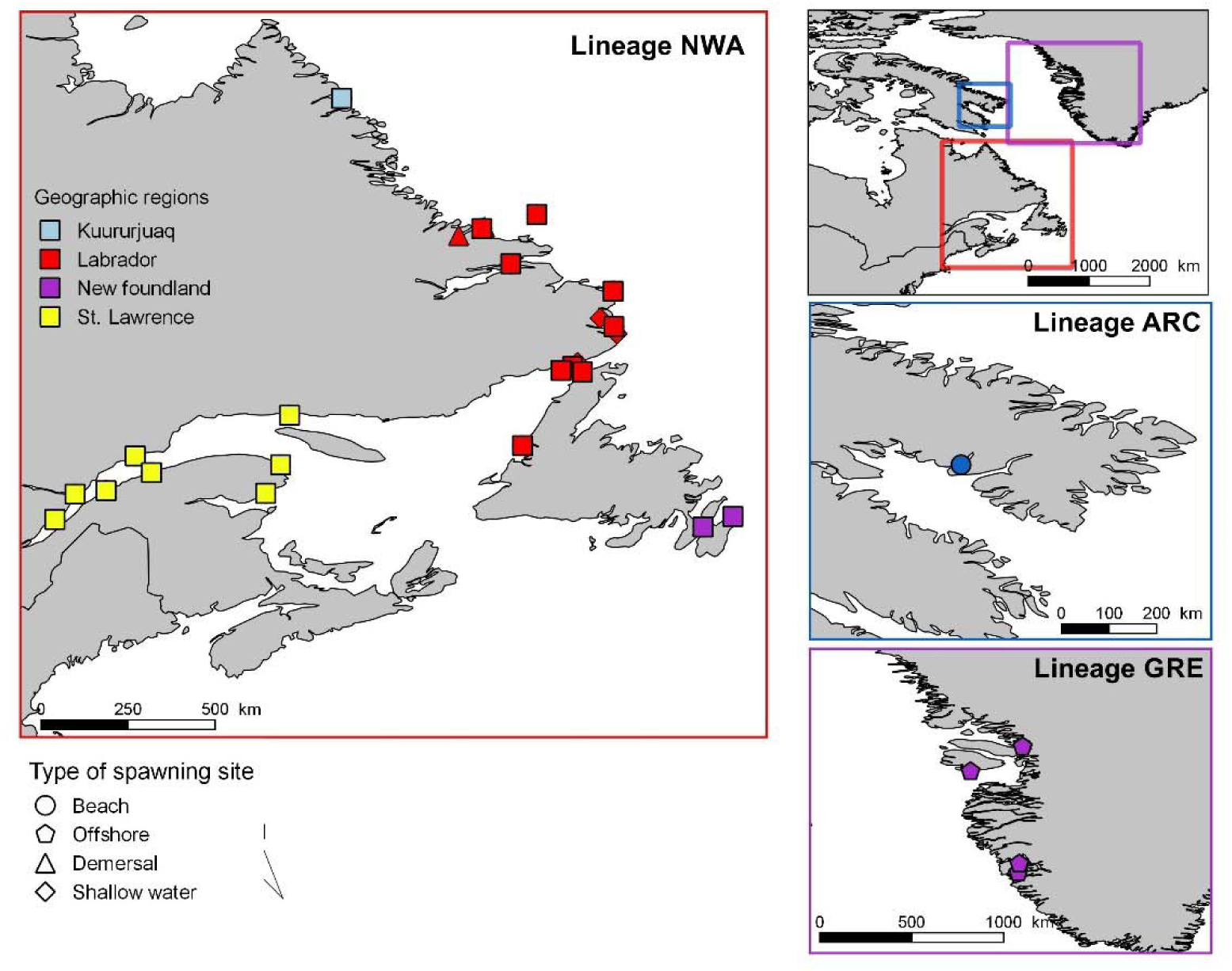
Map of the study area. A total of 31 sampling locations were considered in the distribution range of three glacial lineages of capelin (NWA, ARC and GRE). In the NWA lineage distribution range, we sampled individuals according to three types of spawning sites: beach-spawning sites, shallow-water demersal sites, and deep-water demersal sites. In the ARC lineage, the two sampling sites are very close (few kilometers) and cannot be distinguished on the map.

## Materials and methods

### Genome sequencing and draft assembly

A total of 47 Gbp (∼99x coverage) of sequencing data was generated on the Pacific Biosystems Sequel sequencing machine. Flye version 2.3.3 (Kolmogorov et al. 2019) was run on the PacBio sequencing data specifying a genome size of 700 Mbp (which is between 0.5x and 2.0x of expected genome size of related marine fishes; http://www.genomesize.com/) and otherwise default options. We ran BUSCO version 3.0.1 to assess genome assembly completeness (Simão et al. 2015). While this first version of the capelin genome is not at the chromosome level assembly, it was used in the context of this study to map our RAD sequence reads as well as documenting the synteny with other fish genomes (see below). We annotated the capelin draft genome with the published transcripts of *Clupea harengus* (NCBI assembly accession: ASM96633v1), a related species, using GAWN v0.3.3 (https://github.com/enormandeau/gawn) in order to find transcripts that were physically close to candidate outlier loci. GAWN used GMAP version 2018-07-04 to align the transcripts to the genome.

### Synteny analysis

To estimate the chromosome-scale position of each SNP, we investigated the synteny between the the capelin contigs and four chromosome-level genome assembly of related species, available in the NCBI database, *Esox Lucius* (Rondeau et al. 2014, GenBank assembly accession: GCA_004634155.1)*, Dicentrarchus labrax* (Tine et al. 2014, GCA_000689215.1)*, Sparus aurata* (Pauletto et al. 2018, GCA_003309015.1), and *Takifugu rubripes* (Kai et al. 2011, GCA_000180615.2). The four genomes were first aligned to each other using D-Genies (Cabanettes & Klopp 2018), revealing that similar sequences generally conserve a common location at the chromosome scale, albeit with important rearrangements within each chromosome. This allowed inferring 24 orthologous chromosomes between the four species taken as reference (**Supplementary material**, **Fig. S2**) and to take into account fusion/fission events (**Supplementary material**, **Table S1**). Chromosomes were numbered from 1 to 24 according to their size and their synteny with the *Esox lucius* genome.

All five genomes were masked for repeated sequences using RepeatMasker (Chen 2004) with the *Danio* database. Then, we used the protein sequence aligner *promer* from Mummer 4.0 (Marçais et al. 2018) to localize syntenic regions, *i.e.* protein sequence matches, between the capelin genome and each of the four genomes. Protein sequence matches were filtered to keep only alignments longer than 200 amino acids with similarity over 50%. Then, for each contig of the capelin genome, we calculated the proportion of its length aligning to each chromosome of the four reference genomes. The overall distribution of matches between capelin contigs and the reference chromosomes showed that most contigs aligned primarily to a single orthologous chromosome in the four species, indicating relatively conserved chromosome-scale organization (**Supplementary material**, **Fig. S2**, **Table S1**). We relied on this observation to attribute each of the capelin contig to one of the 24 orthologous chromosomes, i.e. to the chromosome with the longest total alignment across the four species. Contigs were ordered by chromosome and then by size for graphical representations. When no best match to a chromosome was found, or if the best alignment fell on scaffolds from the four reference genomes that were not assembled into chromosome, the contig was placed in the category “Unplaced” (chrU).

### Sampling area and Genotyping-By-Sequencing approach

#### Sampling area and molecular analyses

A total of 1,359 capelins were sampled from 31 sites in the Northwest Atlantic, both in Canadian and Greenland waters (**Fig.1** and **Table 1**), which were expected to include representatives from the three lineages, according to a previous study using mitochondrial DNA and microsatellites genetic markers (Dodson et al. 2007). We sampled 25 spawning sites within the presumed NWA lineage including three demersal shallow-water sites, three demersal deep-water sites, and 19 beach spawning sites. These sites were located in four geographic regions (**Fig.1**): Kuururjuaq, Labrador, Newfoundland, and St. Lawrence. In parallel, we sampled two sites (beach-spawning fish) within the presumed range of the ARC lineage and four offshore sampling sites in the range of the GRE lineage. The median sample size was 46.5 (range: 19 to 50) individuals per site. The fish were collected, sexed, and a piece of fin was preserved in RNAlater.

**Table 1.**
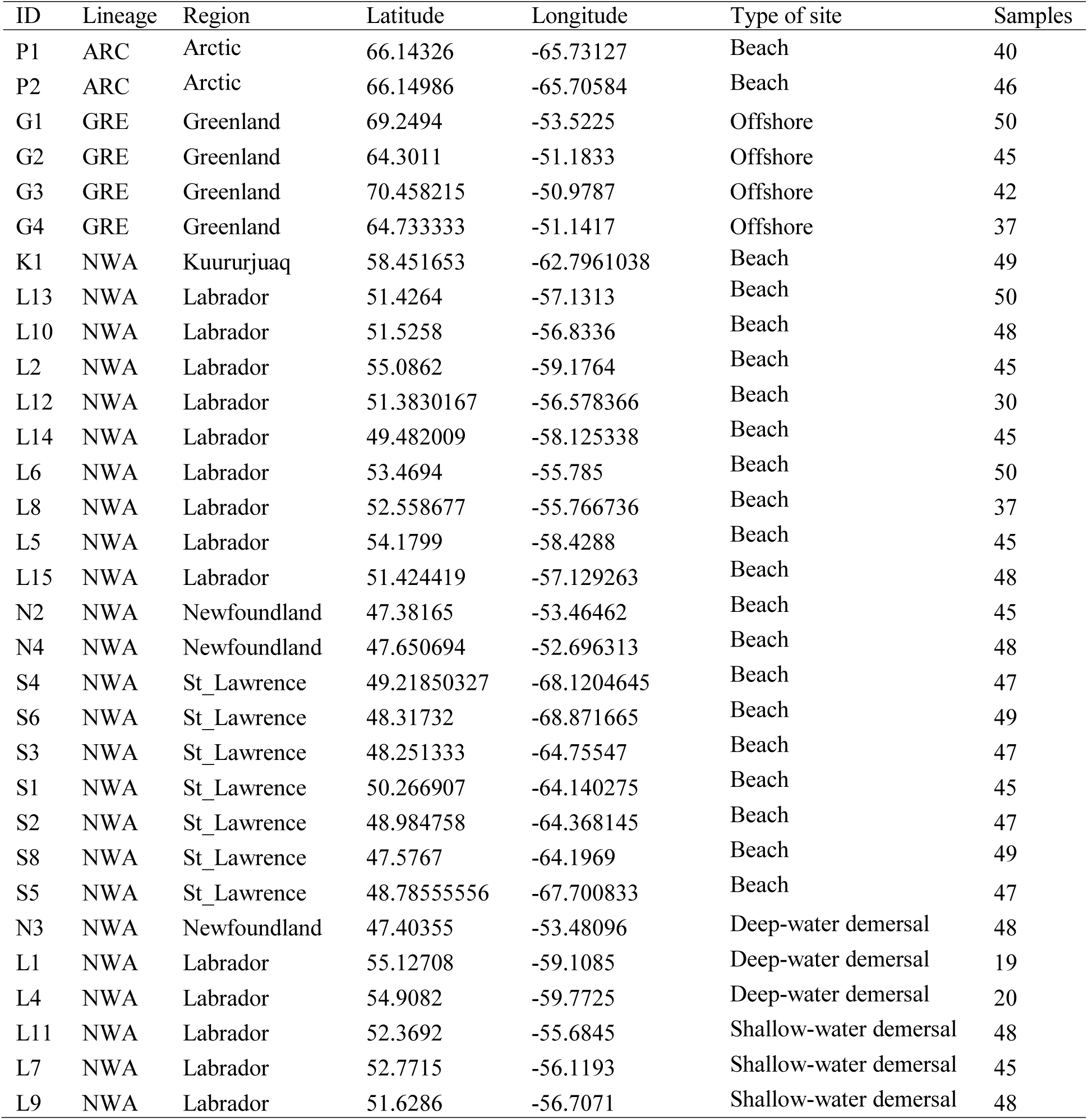
Characteristics of the 31 sampling sites.

Both DNA extractions and library preparation were performed following protocols fully described elsewhere (e.g. Moore et al. 2017; Rougemont et al. 2019). Briefly, DNA was extracted with a salt-based method and an RNAse A (Qiagen) treatment was applied following the manufacturer’s recommendation. DNA quality was assessed using gel electrophoresis. DNA was quantified using a NanoDrop spectrophotometer (Thermo Scientific) and then using QuantiT Picogreen dsDNA Assay Kit (Invitrogen). Concentration of DNA was normalized to 20 ng/μl. Libraries were constructed following a double-digest RAD (restriction site-associated DNA sequencing) protocol modified from Mascher et al. (2013). Genomic DNA was digested with two restriction enzymes (*Pst*I and *Msp*I, Poland et al. 2013) by incubating at 37°C for two hours followed by enzyme inactivation by incubation at 65°C for 20 min. Sequencing adaptors and a unique individual barcode were ligated to each sample using a ligation master mix including T4 ligase. The ligation reaction was completed at 22°C for 2 hours followed by 65°C for 20 min to deactivate the enzymes. Samples were pooled in multiplexes of 48 individuals, ensuring that individuals from each sampling location were sequenced as part of at least six different multiplexes to avoid pool effects. Libraries were size-selected using a BluePippin prep (Sage Science), amplified by PCR and sequenced on the Ion Proton P1v2 chip. Eighty-two individuals were sequenced per chip.

#### DNA sequencing, genotyping, and diversity analysis

Barcodes were removed using cutadapt (Martin 2011) and trimmed to 80 bp, allowing for an error rate of 0.2. They were then demultiplexed using the ‘process_radtags’ module of Stacks v1.44 (Catchen et al. 2013) and aligned to the capelin draft genome assembly using bwa-mem (Li 2013) with default parameters, as detailed below. Then, aligned reads were processed with Stacks v.1.44 for SNPs calling and genotyping. The ‘pstacks’ module was used with a minimum depth of 3 and up to 3 mismatches were allowed in the catalog creation. We then ran the ‘populations’ module to produce a vcf file that was further filtered using python (https://github.com/enormandeau/stacks_workflow) and bash scripts. SNPs were kept if they displayed a read depth greater than 5 and less than 70 (to control for paralogy). Then, SNPs present in a least 70 % in each sampling location and with heterozygosity lower than 0.60 were kept to further control for paralogs and HWE disequilibrium. The resulting vcf file comprised 550,724 SNPs spread over 27,442 loci and was used for demographic inferences with ∂a∂i (See Supplementary material). This vcf file was then subsampled to meet the different assumptions (in particular the use of unlinked SNPs) of the models underlying the different population genetic analyses performed below. We therefore kept one single SNPs per locus (the one with the highest minor allele frequency), further remove SNPs with r^2^ values higher than 0.2 using plink in sliding windows of 50 SNPs. Finally, we excluded any SNPs not present in at least 90 percent of the total dataset resulting in a final vcf of 25,904 high quality SNPs.

### Hierarchical patterns of population genetic structure admixture

#### Genetic differentiation between glacial lineages

We examined genetic differentiation among lineages using the whole dataset (i.e., the 31 sampling sites, Table 1). A principal component analysis (PCA) was performed to investigate the molecular variation among individuals using the R package *Adegenet* (Jombart et al. 2008). Then, we quantified ancestry and admixture proportion using the *snmf* function included in the R package LEA (Frichot & François 2015), with K-values from 1 to 10. We estimated ancestry coefficients levels of K and obtained cross-validations using the cross-entropy criterion with 5% of masked genotypes. Next, we investigated the level of genetic differentiation among sampling sites using the *F_ST_* estimator (Weir & Cockerham 1984) in the STAMPP R package along with 95% confidence intervals using 1000 bootstraps (Pembleton et al. 2013).

#### Geographical and habitat-dependent genetic structure within lineages

A second PCA was executed for each of the three lineages separately to examine within-lineage genetic variation. Admixture analyses were conducted only on the NWA lineage to examine intra-lineage genetic structure. We did not analyze intra-lineage structure for ARC and GRE because of the limited number of sampling sites and the absence of any genetic structure based on PCA results (**Supplementary material**, **Fig. S**4). Moreover, we examined *F_ST_* estimates among all sampling sites, and between demersal and beach spawning sites more specifically.

In addition, we tested for patterns of Isolation-By-Distance (IBD) within the NWA lineage separately for neutral SNPs and SNPs putatively under divergent selection. Due to the limitations of differentiation-based methods and the potentially high false positive rates when searching for outlier loci under divergent selection (Thornton & Jensen 2007, Narum & Hess 2011), we used two distinct approaches to identify neutral markers and markers under selection: 1) an *F_ST_*-based outlier approach implemented in BAYESCAN (Foll & Gaggiotti 2008) that allows detecting neutral markers, markers under balancing or purifying selection, and markers under divergent selection; 2) a hierarchical modeling approach implemented in PCADAPT (2017) permitting to segregate neutral loci and markers under divergent selection. BAYESCAN was run using 5,000 output iterations, a thinning interval of 10, 20 pilot runs of length 10,000, and a burn-in period of 50,000, with prior odds of the neutral model of 10,000 as suggested by Lotterhos and Whitlock (2014). We recorded all loci with a q-value of 0.2 or less, which equates to a false discovery rate of 20%. PCADAPT was run following Luu et al.’s (2017) recommendations. IBD was tested using a linear model in which pair-wise *F_ST_* (*F_ST_*/(1-*F_ST_*)) was included as the response variable and the Euclidean distance (z-scored) was incorporated as an explanatory term. The significance of the effect of Euclidean distance was assessed using adjusted *R^2^* and ANOVA with *F*-test.

### Population structure as revealed by haplogroup distribution in the NWA lineage

Within the NWA lineage, we detected three genetic clusters using the PCA hereafter called haplogroup1, haplogroup2, and haplogroup3, which occurred in all sampling sites across the range of the NWA lineage but that were totally absent in the other two lineages; this pattern was also supported by a clustering analysis performed for the NWA lineage alone (see Results). This was the typical signature of a chromosomal rearrangement (e.g., Berg et al. 2017, Wellband et al. 2019) and we performed the following analyses to confirm this hypothesis. We identified a set of SNPs mainly located in Chr2 and Chr9 associated with these haplogroups based on their loadings along the PC1 axis (**Supplementary material**, **Fig. S4**). We tested whether the heterozygosity of those SNPs differed between haplogroups using a beta regression model; heterozygosity was included in the model as the dependent variable and the haplogroup as discrete explanatory variable. Furthermore, for each sampling site, we tested for Hardy-Weinberg equilibrium when considering the haplogroups as different karyotypes of a putative rearrangement using a chi-square test and a ternary plot implemented in the R package HardyWeinberg (Graffelman et al. 2015).

Next, we examined if the assignment probability of individuals to the haplogroups depended on their sex. We built a logistic regression models for each haplogroup in which individual assignment to the haplogroup was coded as a binary response variable (0 = not assigned to the haplogroup; 1 = assigned) and sex as a discrete explanatory variable. Moreover, we investigated how the frequency of the haplogroup within sampling sites was affected by the types of spawning site (deep-water and shallow-water demersal sites were merged into a single category to increase the power of the analysis). In addition, we examined how bottom temperature and chlorophyll concentration in beach spawning sites influenced haplogroup frequency. The marine data layers were downloaded from Bio-ORACLE (http://www.bio-oracle.org/) and the two environmental variables were extracted using the R package sdmpredictors (Bosch et al. 2017); we present the environmental data in **Supplementary material**, **Table S 14**. We used linear model where the frequency of haplogroup 1 and 2 [(2 × *N_haplogroup1_* + *N_haplogroup2_*) / *N_total_*)] was included as the response variable and the type of spawning sites, temperature, and chlorophyll concentration as explanatory variables. Each explanatory variable was introduced separately in the model to avoid model over-fitting. The significance of explanatory variable effects was evaluated using adjusted *R^2^* and ANOVA with *F*-test.

### Demographic history of divergence in δaδi and identification of inter-lineage outliers

#### Divergence history of the three capelin lineages

We compared the jSFS of our data comprising all three capelin lineages (NWA, ARC, GRE) to those simulated under different demographic models using the diffusion approximation implemented in δaδi (Gutenkunst et al. 2009). We compared four alternative models of historical divergence previously developed (Tine et al. 2014). These include (*i*) a model of Strict Isolation (SI), (*ii*) a model of divergence with continuous gene flow or Isolation with Migration (IM), (*iii*) a model of divergence with initial migration or Ancient Migration (AM) and (*iv*) a model of Secondary Contact (SC) (**Supplementary material, Figure S3**). All models were characterized by the split of an ancestral population of size N*anc* at time *T_split_*into two daughter populations of respective size *N*_1_ and *N*_2_. There is no gene flow between the two populations under the SI model whereas the remaining models incorporate gene flow at a constant rate at each generation. Gene flow is modeled as *M_ij_ = 2Nref.m_ij_* and can be asymmetric, so that two independent migration rates *m_12_* (from population 2 to 1) and *m_21_* (from population 1 to 2) were modeled, with *Nref* being the size of the reference population. Under AM, gene flow occurred between *T_split_*and *T_am_* and was followed by a period of strict isolation. Under IM, the two diverging populations exchange migrants at a constant rate at each generation from *T_split_* to present time. Under the SC model, the population evolved in strict isolation from *T_split_*to *T_sc,_* the time at which a secondary contact had been taking place continuously up to the present time.

These models incorporate the effects of selection at linked sites locally affecting *Ne* and the effect of differential introgression affecting the rate of migration (*m*) along the genome (Tine et al. 2014, Rougemont et al. 2017, Le Moan et al. 2016, Rougeux et al. 2017). Heterogeneous introgression was modeled using two categories of loci occurring in proportions *P* (i.e. loci with a migration rates *m_12_* and *m_21_*) and *1-P* (i.e. loci with a reduced effective migration rates m*e_12_*and *me_21_*) across the genome. The same procedure was used to account for linked selection by defining two categories of loci (Q and 1-Q) with varying effective size. Then, the effect of linked selection was quantified using a Hill-Robertson scaling factor (*Hrf*, Rougeux et al. 2017) relating the effective population size of loci influenced by selection (*Nr = Hrf* * *Ne*) to that of neutral loci (*Ne*).

Three jJSFS for all three pairs of lineages identified (NWA vs. ARC, NWA vs. GRE, ARC vs. GRE) were constructed by merging all localities from the ARC lineage together, all localities from the GRE lineage, and randomly chosen localities within the NWA lineage showing non-significant genomic differentiation. Random sampling within NWA lineage was further reduced to match the number of individuals equal to that of the other two lineages. Details of the filtering procedure are provided in Supplementary materials (see the section “Filtering procedure for demographic inference with δaδi”) along with the jSFS. Three optimization steps consisting of 1) “hot” simulated annealing 2) “cold” simulated annealing and 3) BFGS -Broyden–Fletcher– Goldfarb–Shanno- algorithm were used for model fitting following Tine et al. (2014). Twenty-five replicates were performed for each model and the replicate with the lowest AIC value by model was kept. Model choice was performed using Akaike information criterion (AIC), ΔAIC and AIC weights. Standard deviations around parameter estimates were obtained using the Godambe Information Matrix (Coffman et al. 2015) with 100 bootstraps dataset. We attempted to estimate biological parameters assuming a generation time of 3.8 years (Dodson et al. 2007) and a standard mutation rate of value of 1e-8 mutation/bp/generation. Scripts to reproduce the analysis are available on GitHub at: https://github.com/QuentinRougemont/DemographicInference.

To account for the possible confounding effect of the two chromosomal blocks (on separate Chr2 and Chr9) likely representing a chromosomal rearrangement on our model choice and parameter estimates, we repeated the analysis in two ways. First, we excluded the whole Chr2 and Chr9 from the jSFS and repeated the δaδi analysis between all lineages. Second, to make sure that the occurrence of genetic differentiation between groups did not bias our model choice, we separated individuals from the NWA clade into three haplogroups based on their coordinates from the PCA. We then created a new set of pairwise jSFS including all SNPs and repeated all δaδi analyses by comparing the haplogroups 1 and 3 of the NWA lineage against GRE and ARC lineages. We repeated this same analysis after excluding the chr2 and chr9 from the jSFS. Last, to better understand the origin of these three haplogroups, we constructed a final jSFS for the haplogroup1 *vs* haplogroup3 that we hypothesized to be homokaryotes (haplogroup 2 hypothetically being the heterokaryote, see Results), each for a different version of the putative chromosomal rearrangement (see the section *Results*). We then performed a set of demographic inference by fitting the same set of demographic models.

#### Coalescent simulation and identification of inter-lineage outliers

We used the demographic parameter estimates along with their standard deviation under each best model for each pair of lineages to perform 4,000,000 coalescent simulations using the simulator Ms (Hudson 2002). Here, we assumed a strictly neutral model with homogeneous population size and homogeneous migration rate. For each SNP, we computed levels of expected heterozygosity (He) as well as Weir & Cockerham *F_ST_* estimator between lineages. As proposed by Beaumont & Nichols (1996) we computed *F*_ST_ quantiles in heterozygosity intervals of 0.025. This enabled us to detect candidate loci potentially under divergent selection between lineages, but with a more realistic and complex demographic model specifically inferred for our focal species. This method is similar to that used in Le Moan et al. (2016) and Rougemont et al. (2017) but also considers the uncertainty surrounding parameter estimates (Leroy et al. 2019). Loci departing from our neutral envelope (outside the 99.99^th^ upper quantiles of conditioned *F_ST_* distribution) were inferred as candidate outliers. Finally, to performed gene annotation and better identify candidate loci we used the sequences of the markers potentially under divergent selection between the three lineages to perform a blast (Altschul et al. 1990) analysis on the NCBI NR DATABASE using an e-value threshold of 1x10-10 and a minimum length of 90 bp.

### Searching for signals of local adaptation pattern within the NWA lineage

#### Genotype-environment associations and local adaptation to the type of spawning sites

We examined the molecular signature of local adaptation to spawning sites within the NWA lineage only, since not enough sampling sites were available within the other two lineages. More specifically, we aimed at detecting outlier loci associated with types of breeding sites distributed along a water depth gradient: the beach spawning site located in the intertidal zone, the demersal spawning site in shallow water (from 2 to 5 m), and the demersal spawning site in deeper water (from to 10 to 20 meters). As recommended by Forester et al. (2018), we used a combination of latent factor mixed models (LFMM; Frichot et al. 2013, Frichot & François 2015) and partial redundancy analysis (pRDA; ‘rda’ function in ‘vegan’ R package; Lasky et al. 2012, Forrester et al. 2018) to detect candidate loci.

We ran LFMMs, a GEA approach that controls for population structure using latent factors, using updated parameters as recommended by the authors (http://membres-timc.imag.fr/Olivier.Francois/LEA/tutorial.htm). The analysis was conducted in the R package LEA. Models were parameterized using the K-value (K = 2) obtained in our previous clustering analyses in LEA. By doing so, we accounted for the genetic variation associated with the haplogroups in our analyses of genotype-environment associations. We ran each model 10 times with 5,000 iterations and a burnin of 2,500. The median of the squared z-scores was used to rank loci and to calculate a genomic inflation factor to evaluate model fit (François et al. 2016).

We used a partial redundancy analysis (pRDA) to detect multi-locus outliers that were associated with the type of spawning sites after controlling for the haplogroups (see Legendre & Legendre 2012, Laporte et al. 2016, Le Luyer et al. 2017, Forester et al. 2018 for details). Standard deviation of markers scores was multiplied by 3.5 to establish the multi-locus outlier detection p-value at 0.001 (Forester et al. 2018). We retained the outliers that were detected by both LFMM and pRDA to reduce the number of false positives as recommended by Forester et al. (2018) and represented the overlap with a Venn diagram.

For each outlier locus associated to the type of spawning site, we identified the transcripts present within a 1 kbp window around the loci by using BEDtools intersect v2.26.0 (Quinlan & Hall 2010). We then used go_enrichment (https://github.com/enormandeau/go_enrichment) to annotate the same transcriptome by blasting the transcripts on the Swissprot database (blast tools 2.7.1, Swissprot from 2018-05-01). We also tested for the presence of over and under-represented GO terms using GOAtools (v0.6.1, pval = 0.05) and filtered the outputs of GOAtools to keep only GO terms with levels 3 or less and with an FDR value equal to or below 0.1.

#### Genotype-environment associations and local adaptation to environmental conditions in beach spawning sites

We investigated the molecular signature of local adaptation to local environmental conditions across beach spawning sites within the NWA lineage (i.e., 19 sampling sites). We considered two environmental variables: temperature and chlorophyll concentration as a proxy of trophic productivity, which may have dramatic effects on embryonic and larval performances (Frank & Leggett 1981a, 1981b, 1982; Leggett et al. 1984; but see Murphy et al. 2018). The correlation between temperature and chlorophyll concentration (*r* = 0.47) was relatively low. For this reason, these two variables were simultaneously introduced in the pRDA analyses presented below. We used the same approach than the one used for detecting the molecular signature of adaptation to spawning sites, that is we determined candidate loci associated with three variables using the combination of LFMM and pRDA. In LFMM, the variables were introduced one by one in the models to identify the candidate loci associated to each variable. We then used the same procedure described for spawning sites analyses for go_enrichment and blasting.

## Results

### Draft genome assembly and statistics

The resulting draft genome assembly was 490 Mbp with a N50 contig size of 230 Kbp. Running BUSCO with the *Actinopterygii* dataset on the assembly found 4070 complete, 209 fragmented and 305 missing genes out of a total of 4584 genes searched. The genome assembly is available at DOI: 10.6084/m9.figshare.9752558. Further in-depth analysis of the genome will be provided elsewhere as its main purpose here was to provide a reference for reads mapping.

#### Genetic differentiation between glacial lineages

The raw vcf from stacks contained 1,081,533 SNPs spread on 67,305 loci before filtration. After filtration we kept a final set of 25,904 SNPs spread on 25,904 loci (see details in methods).

A high and heterogeneous level of differentiation was found along the genome for the comparison between lineages (**Supplementary material**, **Fig.S1**). The first and second axes of the PCA explained 8.2% of the genetic variation (**Fig.2A**) and they supported the pattern found in the. The ARC linage was the most divergent among all three lineages. The clustering analysis also supported the existence of three lineages (K = 3) with the absence of admixture among lineages (**Fig.2C**). This pattern suggested very limited, migration, if any, between each pair of lineages despite the absence of physical barriers between them. Accordingly, measures of net sequence divergence (*d*_a_) which, unlike, are not affected by variation in the common ancestor revealed a level of net divergence that was about twice higher between the ARC and the GRE lineages (*d*_a_ = 0.00146) and between the ARC and NWA lineages (*d*_a_ = 0.00140) than between the GRE and NWA clades (*d*_a_ = 0.00067). Moreover, the mean was 0.2435 between NWA and GRE, 0.3783 between NWA and ARC lineages, and 0.4106 between GRE and ARC (**Fig.2C**)., thus revealing a very high level of genetic differentiation for a marine species (DeWoody & Avise 2005).

**Fig. 2.**
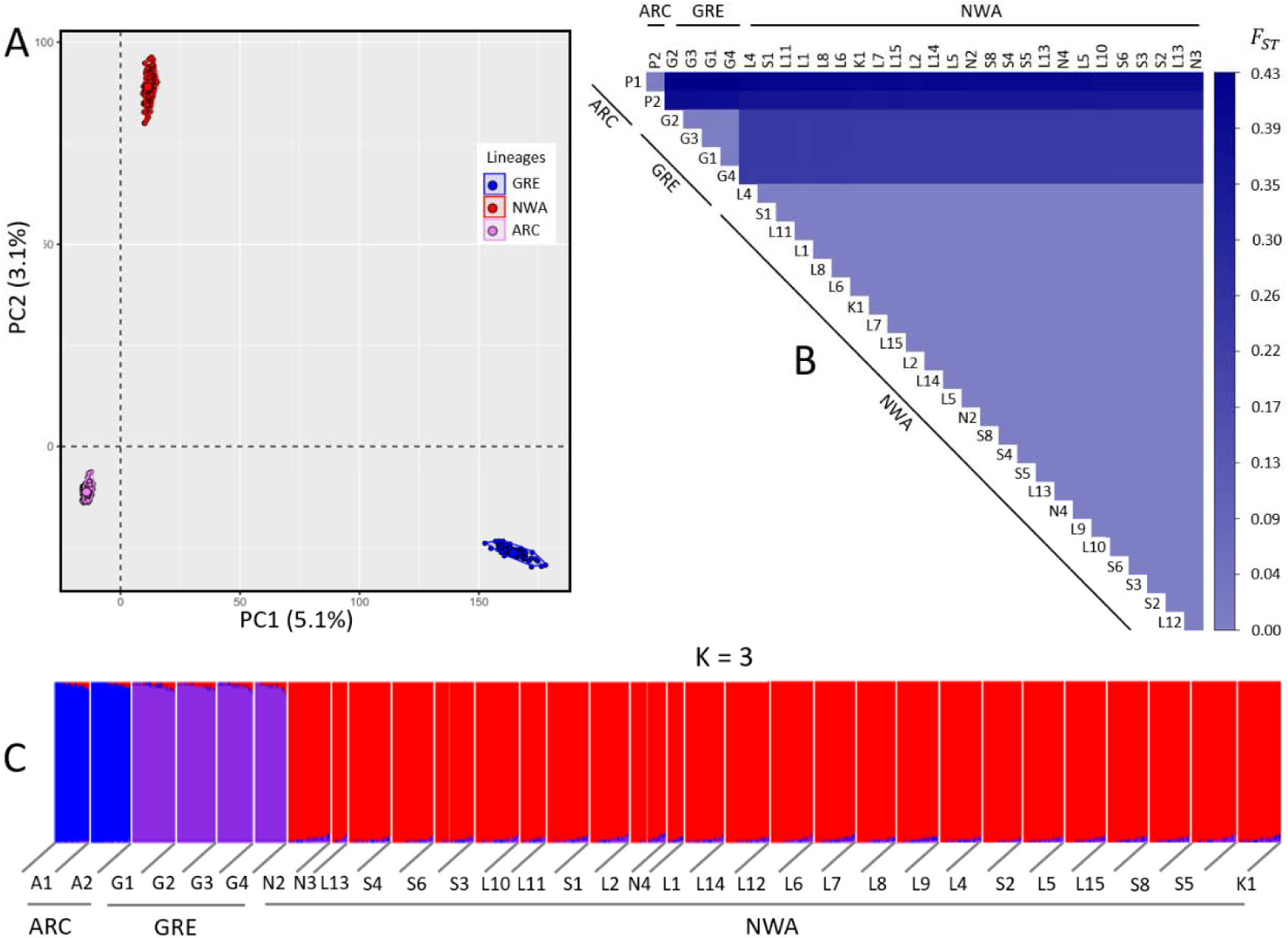
Genetic structure and differentiation among the three lineages (NWA, GRE, and ARC) of capelin. (A) Principal Component Analysis (axis 1 and 2) showing the genetic variation among the lineages. (B) Heatmap of among and within three lineages. (C) Clustering analysis performed using LEA R package.

**Fig. 3.**
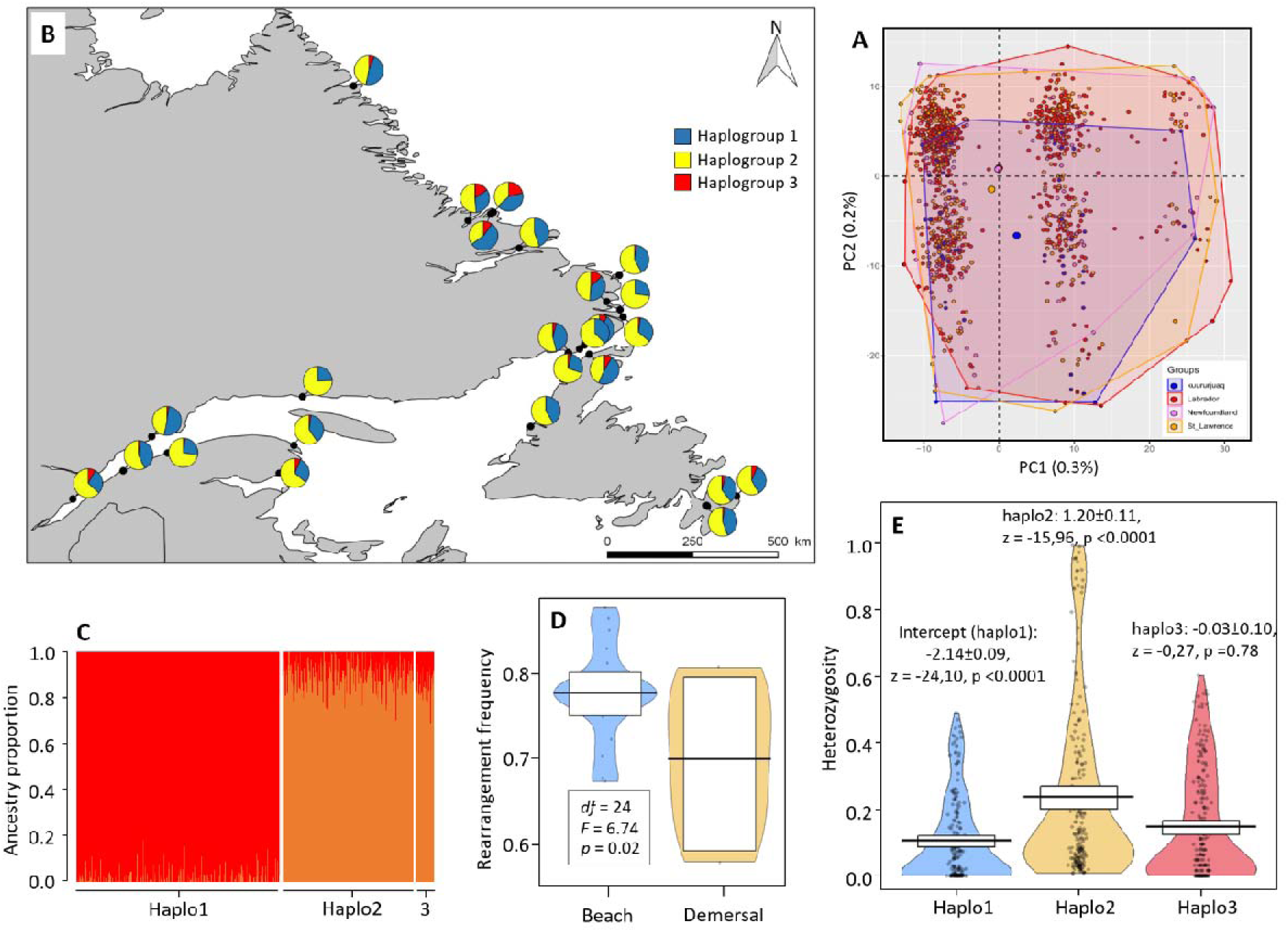
Spatial variation of the putative chromosomal rearrangement (involving chromosomes 2 and 9) within the NWA lineage of capelin. (A) Principal Component Analysis (axis 1 and 2) showing the three clusters of individuals corresponding to the three haplogroups. (B) Relative proportion of the three haplogroups among the 25 sampled spawning throughout the distribution area of the NWA lineage. (C) Ancestry proportion ordered by haplogroups from clustering analyses performed using LEA R package. All individuals of haplogroup1 are contained in the “red” genetic cluster red whereas all individuals of the haplogroups 2 and 3 are admixed between the “orange” and “red” genetic cluster. (D) Frequency of the rearrangement in the demersal (deep-water sites + shallow-water sites) and beach spawning sites; we provide the outputs of an ANOVA (degrees of freedom df, F statistics, and p-value) testing the effect of the type of spawning site on the rearrangement frequency. (E) Heterozygosity analysis focused on the contigs contained in the potential chromosomal rearrangement in chromosomes Chr2 and Chr9. We provide the outputs of the beta regression model (the haplogroups were coded as a discrete variable in the model): intercept for haplogroup1, and the slope coefficients for haplogroup2 and haplogroup3, and their associated standard error and statistical test (z statistic and p-value).

#### Geographical and habitat-dependent genetic structure within lineages

At the intra-lineage level, the pair-wise *F_ST_* values between sampling sites were three order of magnitude lower than observed between the three lineages (**Fig.2B**). The mean *F_ST_* was 0.0033 within the GRE lineage (ranging from 0.0021 to 0.0054) and 0.0044 in the ARC lineage (only two sampling sites in this lineage). Within NWA lineage, the mean *F_ST_* was 0.0010 (ranging from 0 to 0.0090); 91% of the *F_ST_* estimates had a *p*-*value* lower than 0.01 and 95% CI that did not include 0 (see **Supplementary material**, **Table S13**), indicating a very weak yet significant population structure, as typically observed in marine species. Within the NWA lineage, the mean *F_ST_* among shallow-water spawning sites and among deep-water demersal sites was 0.0019 and 0.0017, respectively whereas it was 0.0029 among beach spawning sites. The mean *F_ST_* between shallow-water demersal sites and beach sites and between deep-water demersal sites and beach sites was 0.0031 and 0.0027, respectively. This suggests a more pronounced genetic connectivity among demersal than beach spawning capelin as well as a more restricted gene flow between demersal and beach spawning capelin than within demersal capelin.

We then tested for IBD within the NWA lineage, using neutral loci and loci under selection (**Fig.5**). Using BAYESCAN, we identified 12,278 neutral SNPs, 2,462 SNPs under balancing selection, and 20 SNPs under divergent selection. Using PCADAPT, we detected 17,317 neutral SNPs and 43 SNPs under divergent selection. After removing the SNPs under balancing selection, we compared the neutral SNPs and SNPs putatively under divergent selection defined with the two approaches (**Fig.5A**). A total of 12,060 SNPs (97.7% of the total of SNPs) were identified as neutral by the BAYESCAN and PCADAPT while the number of SNPs identified as “under divergent selection” was low (8 SNPs, 0.01%). We used the 12,060 SNPs to examine neutral IBD and we used the SNPs detected as outliers by at least one of the two methods to examine non-neutral IBD. We detected a weak signal of IBD using the two sets of markers (with SNPs under divergent selection: *F* = 5.38, = 0.01, *p* = 0.02; with neutral SNPs: *F* = 3.89, = 0.01, *p* = 0.05) (**Fig.5B**). This result suggests that besides the effect of spawning mode on patterns of population structure, geographic distance also plays a role, albeit very modest.

**Fig. 4.**
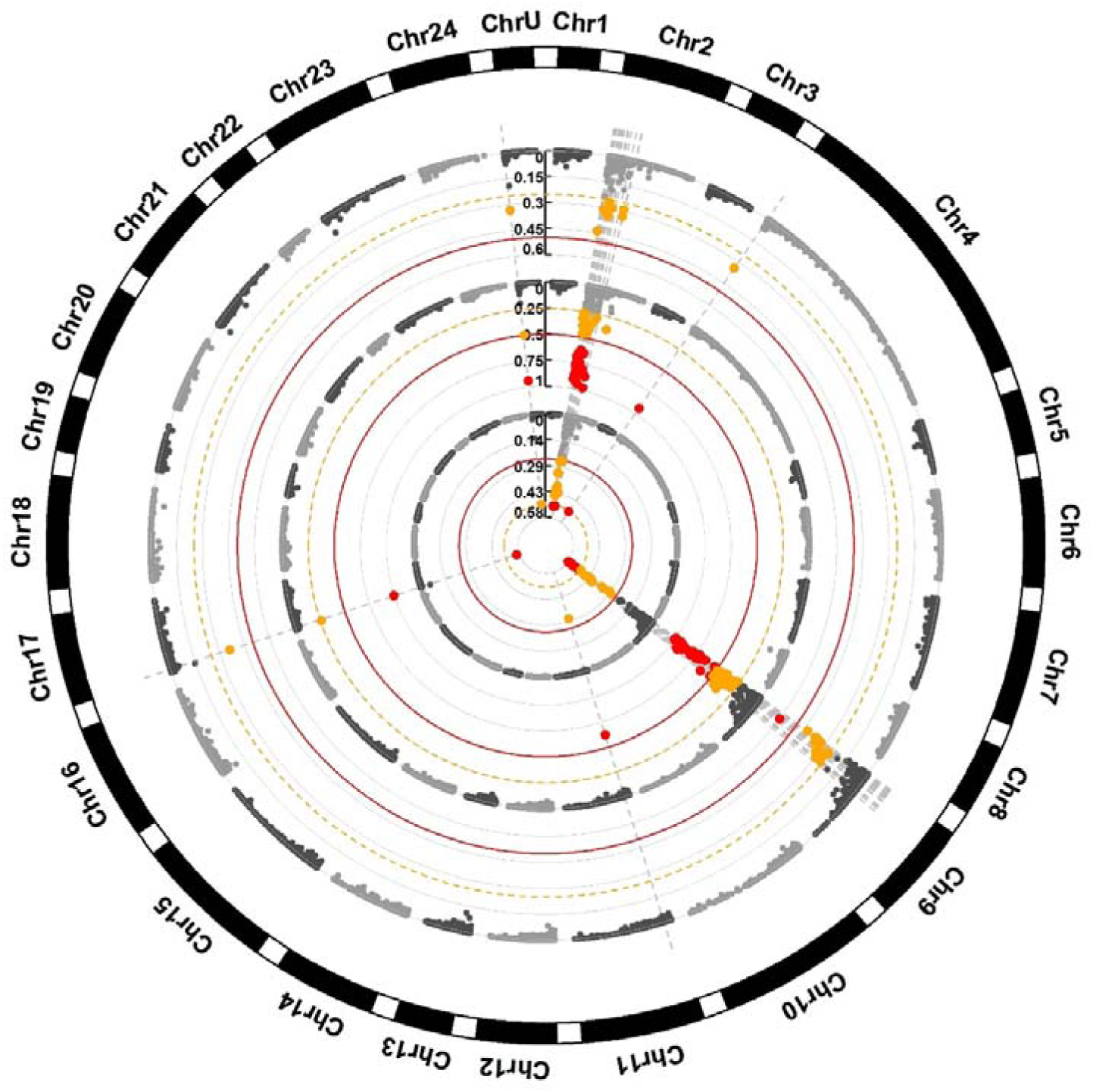
Manhattan plots of the *F_ST_* values between the three haplogroups within the NWA lineage. The external circle shows the *F_ST_* between haplogroups 1 and 2. The medium circle presents the *F_ST_* between haplogroups 1 and 3. The central circle shows the *F_ST_* between haplogroups 2 and 3. Capelin contigs were placed into 24 ancestral chromosomes based on synteny with 4 related species. Contigs that were not assigned to a chromosome were aggregated in the group ChrU. As the order of the contigs within the chromosomes was not conserved and not always consistent among genome comparison, contigs were ranked according to their size. SNPs having *F_ST_* less than 0.25 are shown in grey, SNPs with *F_ST_* ranging from 0.25 to 0.50 are shown in yellow, and SNPs with *F_ST_* higher than 0.50 are shown in red.

**Fig. 5.**
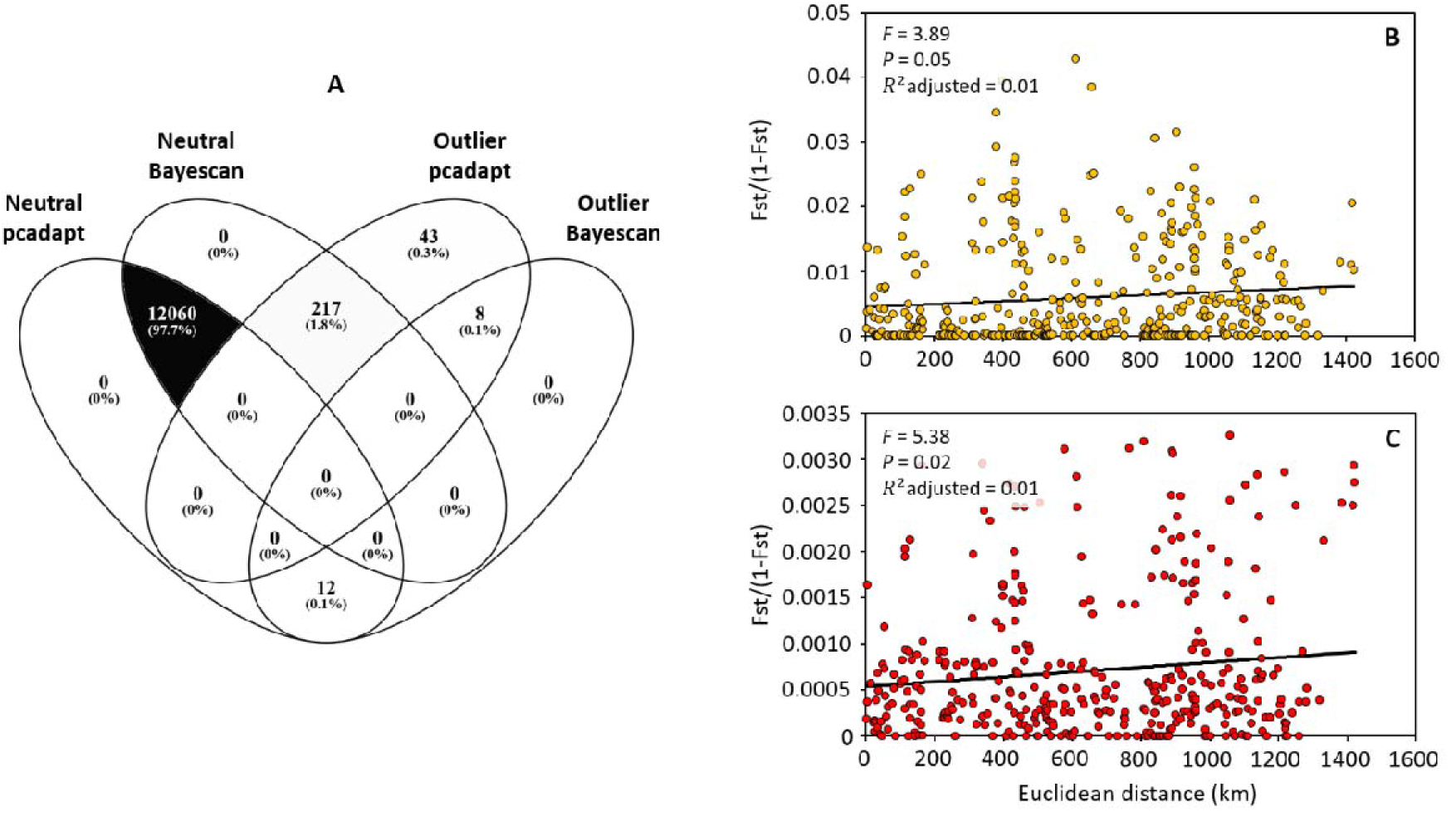
Within the NWA lineage, Outlier detection and Isolation-By-Distance (IBD) analyses for neutral SNPs and SNPs under selection. (A) Venn diagram showing the neutral markers and markers under selection detected with PCADAPT and BAYESCAN. (B) IBD with neutral markers. (C) IBD with markers under divergent selection.

#### Evidence for a major chromosomal rearrangement associated with population within the NWA lineage

The PCA performed for the GRE and ARC lineages did not display any genetic structure (**Supplementary material**, **Fig.S4**) either. By contrast, the PCA revealed a pronounced genetic structure within the NWA lineage (**Fig.3A**), with three genetic clusters along PC1 (explaining 0.3% of the variation). The distribution of the clusters did not show any geographical pattern (**Fig.3B**, but see below). A set of 248 SNPs displayed the highest loading along the first axis. Mapping of these 248 SNPs revealed that 85% of them were preferentially located on ancestral chromosomes Chr2 (n = 74 SNPs) and Chr9 (n = 139 SNPs) whereas an average of two SNPs was mapped on the remaining chromosomes. The PCA performed for Chr2 and Chr9 separately showed similar clustering patterns (**Supplementary material**, **Fig.S4**). Altogether, these observations led us to hypothesize that such genetic substructure may be due to a chromosomal inversion in Chr2 and Chr9 that are likely fused in the capelin. We thus considered the three clusters as three haplogroups. Based on the PCA, 57% of the individuals were assigned to the haplogroup1, 38% to the haplogroup2, and 5% to the haplogroup3.

SNPs with the highest levels of differentiation between haplogroups were also predominantly located on Chr2 and Chr9 (**Fig.4**). The highest genetic differentiation was observed between haplogroups1 and 3, with a genome-wide *F_ST_* of 0.0047 but a *F_ST_* of 50 times higher (0.29) when we only considered the SNPs located in the Chr2-Chr9 rearrangement. This was higher than the differentiation observed between the haplogroup and the two other lineages for the same genomic region (*F_ST_*group1 and GRE = 0.055; *F_ST_* group1 and ARC = 0.0124; *F_ST_* group3 vs GRE = 0.072; *F_ST_*group3 vs ARC = 0.148). The beta regression model also indicated that the heterozygosity of the SNPs from Chr2 and Chr9 differed among haplogroups (**Fig.3E**): it was highest in haplogroup2 and was relatively similar in haplogroup1 and haplogroup3. The clustering approach also supported the existence of those haplogroups. We detected two genetic groups (K = 2) spread over all the sampling sites. All individuals of the haplogroup 1 are contained in one genetic group whereas all the individuals of the haplogroup2 and haplogroup3 are admixed with the other genetic group (**Fig.3C**). Moreover, when considering haplogroup1 and haplogroup3 as the homokaryotes and haplogroup2 as the heterokaryote for the putative rearrangement, there was no deviation from Hardy-Weinberg equilibrium at any sampling site (Supplementary material, Table S12), suggesting mendelian inheritance of markers found in this rearrangement. Altogether these observations suggest that haplogroup 2 may represent a group of heterozygotes bearing two versions, perhaps partially non-recombining of haploblocks that exist at homozygote state in the haplogroups 1 and 3.

Haplogroups 1 and 2 were present in 100% of the sampling sites, while haplogroup 3 occurred in only 78% of the sites. The frequency of haplogroups varied among sampling sites (**Fig.3B**): the proportion of individuals in haplogroup 1 ranged from 35 to 73%, the proportion ranged from 24 to 52% for haplogroup 2, and from 0 to 21% for haplogroup 3. Logistic models indicated that the probability that an individual was assigned to haplogroup 1 or haplogroup 2 (encompassing 95% of the individuals) was not influenced by its sex (haplogroup 1: *F* = 0.33, *p*-value = 0.33; haplogroup 2: *F* = 0.69, *p*-value = 0.41). The probability that an individual was assigned to haplogroup 3 was sex-dependent (*β_males_* = -0.97±0.34, *F* = 8.73, *p*-value = 0.003) although this was likely due to the small number of individuals assigned to haplogroup 3. Furthermore, linear models indicated that the frequency of the putative rearrangement differed between the type of sites (**Fig.3D**). The frequency of haplogroups 1 and 2 combined was higher in beach spawning sites than in demersal sites (both deep-water and shallow-water sites combined) (*df* = 24, *F* = 6.74, *p* = 0.02). By contrast, rearrangement frequency was not influenced by temperature (*df* = 18, *F* = 0.67, *p* = 0.42) or chlorophyll concentration (*df* = 18, *F* = 0.08, *p* = 0.83).

### Demographic history of divergence

#### History of divergence among glacial lineages

The divergence history inferred with ∂a∂i revealed the occurrence of ongoing gene flow between all three lineages whereas models of strict isolation were not supported (**Fig.6**; for the model selection procedure, see **Supplementary material, Table S3**). In two inter-lineage comparisons, GRE *vs* NWA lineages and GRE *vs* ARC lineages, the models of secondary contacts including both linked selection and variable introgression rate among loci (“SC2N2m”) received the highest statistical support (wAIC = 1; **Supplementary material, Table S3, Fig.6**). In the comparison of the NWA *vs* ARC lineages, the model of isolation with migration including both linked selection and variable introgression rate among loci (“IM2N2m”) received the highest statistical support (wAIC = 1; **Supplementary material, Table S3**). We verified if non-modelled structure due to the presence of the putative chromosomal re-arrangement defined above may have biased our results. We found that excluding the Chr2 and Chr9 from the analysis did not affect the model choice with the IM2N2m still being the best model between NWA and ARC (wAIC = 1, Supplementary Material, Table S3, Fig S8). The SC2N2m also remained the best between NWA and GRE (wAIC = 1, Supplementary Material, Table S3, Fig S8). After separating the NWA lineage into three haplogroups (corresponding to the different versions of the arrangement) and repeating the ∂a∂i analysis, the SC2N2m model was still supported in comparison against the GRE lineage (wAIC = 1; **Supplementary material, Table S3**) providing very good fit to the data (**Fig.7**). The IM2N2m was also still supported when comparing the three NWA haplogroups against the ARC lineage (wAIC = 1; **Supplementary material, Table S3**; **Fig.7**). The model choice remained the same after removing the chromosome 2 and 9 (wAIC = 1; **Supplementary material, Table S3**). Detailed parameter estimates when splitting the NWA and when excluding the Chr2 and Chr9 were largely similar to those obtained globally and often fall within the confidence intervals obtained globally (**Supplementary Table S4**). Therefore, we focused mainly on parameter estimates obtained globally presented in **Table 2**. Parameter estimates revealed that the NWA and ARC lineages diverged approximately 3.8 MyA, [95% CI: 3.6 – 3,8 MyA]. The GRE was inferred to have diverged from the ARC lineage approximately 2.7 MyA, [95% CI: 2.1 – 3,2MyA). The NWA vs GRE comparison indicated a divergence approximately ∼1.8 MyA [95% CI: 0.8 – 2,8 MyA). These splits time were older when removing the Chr2 and Chr9 and when separating by haplogroup with respective averaged value of 4,3 MyA for the NWA vs ARC split and 2.5 MyA for the NWA vs GRE lineages (**Supplementary material, Table S4**) suggesting a more recent time to the most recent common ancestor (TMRCA) for the Chr2 and Chr9. In all models of secondary contact, the period of gene-flow represented from 22% to 48% for the total divergence time, translating into long periods of gene flow. This gene flow ranged from 1.3e^-7^ to 1.6^e-6^ across all comparisons and was highly asymmetric and heterogeneous. Indeed, under SC on average, half of the genome displayed reduced introgression rate, while our inference under IM2N2m indicated that 99% of the genome displayed reduced introgression, with this latter value decreasing to 88% when separating by haplogroup and excluding Chr2 and Chr9. The rate of introgression under IM2N2m between the NWA and the ARC lineage appears to be an order of magnitude lower than in other comparisons (**Supplementary material, Table S4**). Finally, in all comparisons, approximately half of the genome appears potentially under the effect of selection at linked sites and the hrf factor indicated that the effective population size in these regions would be approximately 10 to 24% of the total effective population size. According to our estimates, these effective population sizes were high, varying from approximately 220,000 (sd = 7,600) in the ARC lineage under IM2N2m to more than 1,330,000 (sd = 82,000) in the NWA lineage under the same model, in line with the very high levels of polymorphisms observed across loci (**Table 2**; see also **Supplementary material, Table S4** for details of parameters estimates when separating by haplogroups).

**Fig. 6.**
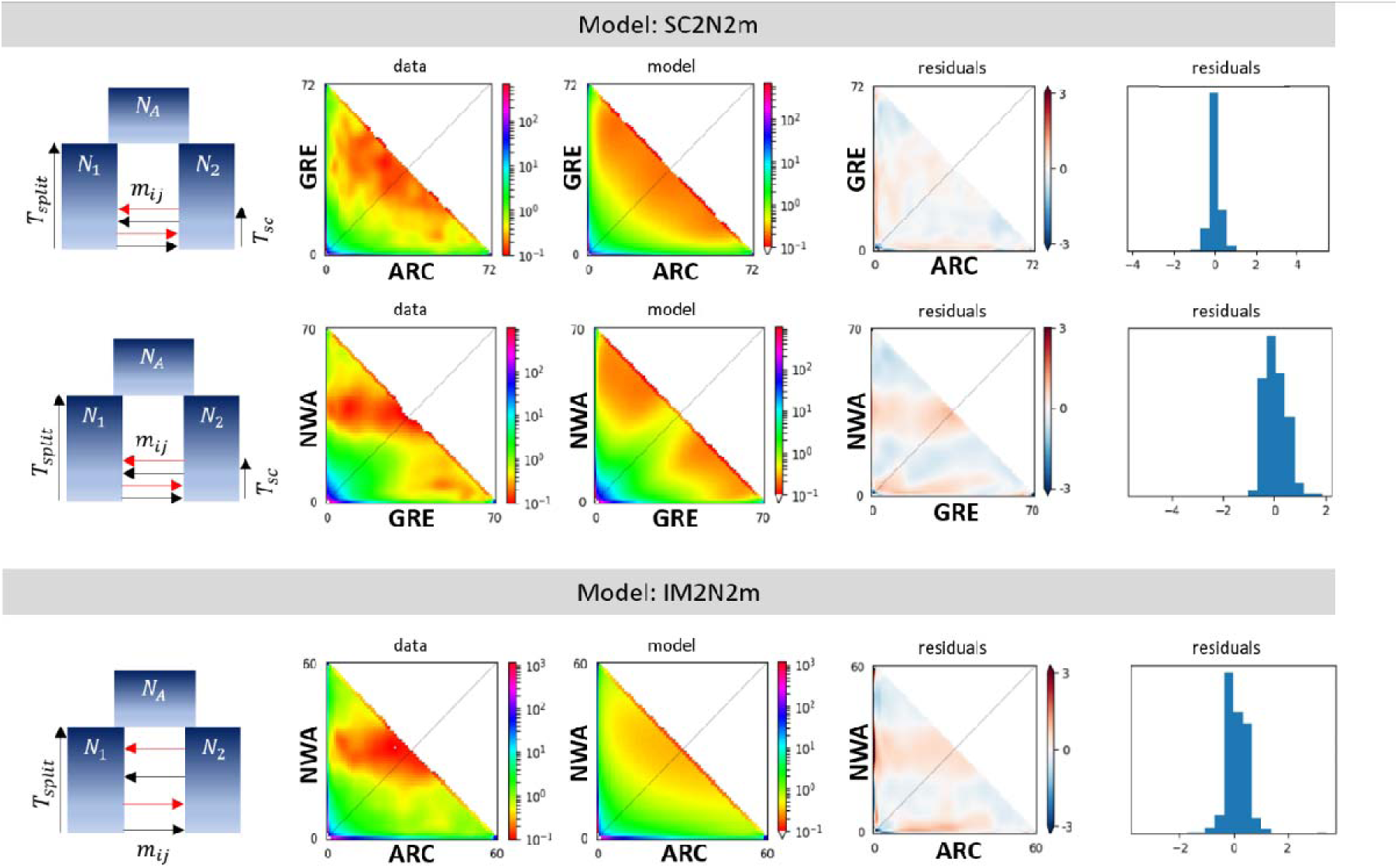
Observed and best fitted jSFS obtained from ∂a∂i (left panels) and residuals of the fitted models (right panels). The best model was the SC2N2m (model of secondary contacts including both linked selection and variable introgression rate among loci) in the comparison between ARC vs GRE and GRE vs NWA whereas the IM2N2m model (model of isolation with migration including both linked selection and variable introgression rate among loci) best fits the data in the ARC vs NWA comparison.

**Fig. 7.**
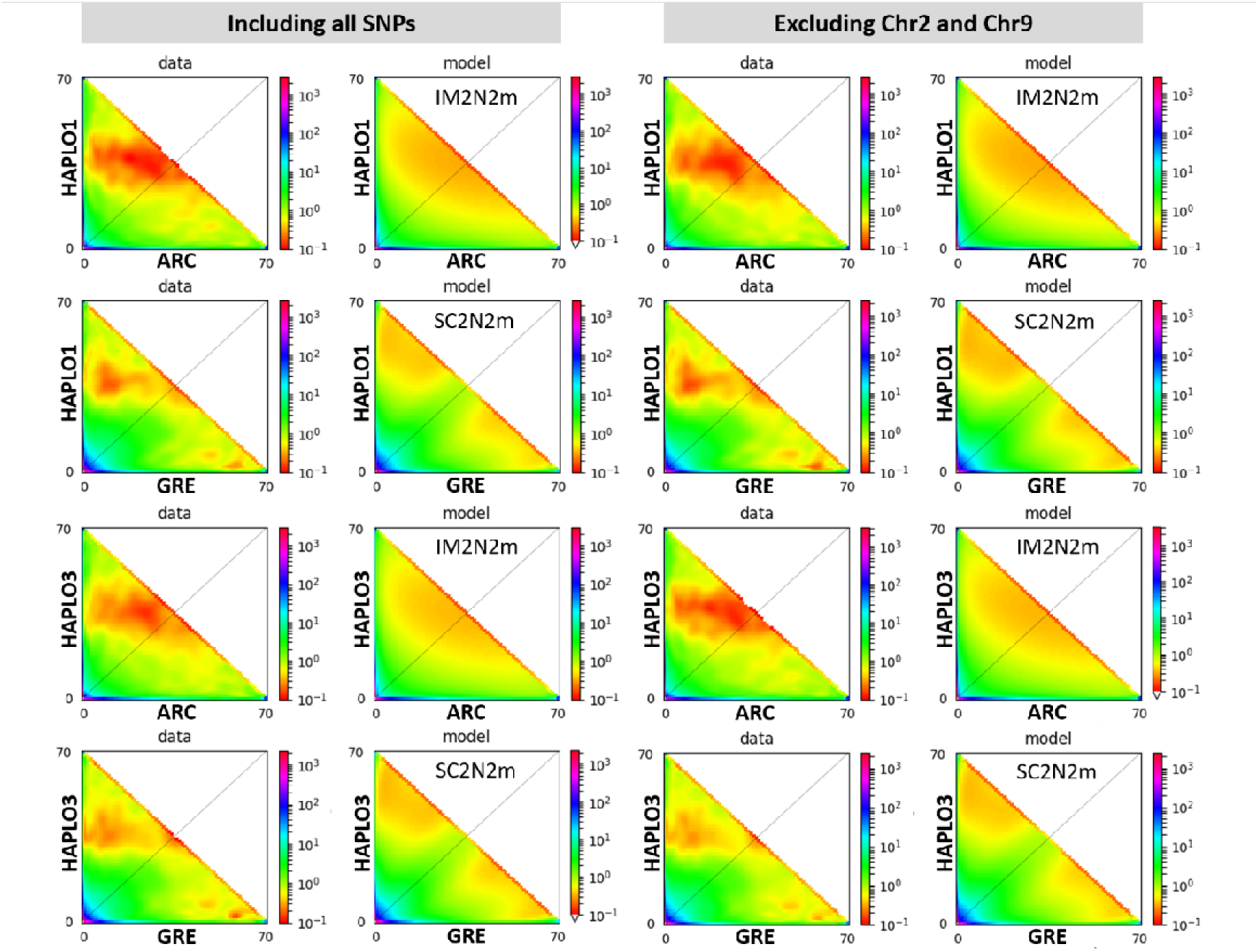
Observed and best fitted jSFS obtained from ∂a∂i split by haplogroups, including all the SNPs (left parts) and excluding SNPs mapping into Chr2 and Chr9. (The best model was the same as in Fig 6, indicating that our inferred models were not biased by a chromosomal arrangement or any other structure associated with the haplogroups within the NWA lineage, see also supplementary figure S7). The haplogroup 2 was heterozygous between haplogroup1 and 3 and therefore was excluded from the analysis.

**Table 2.**
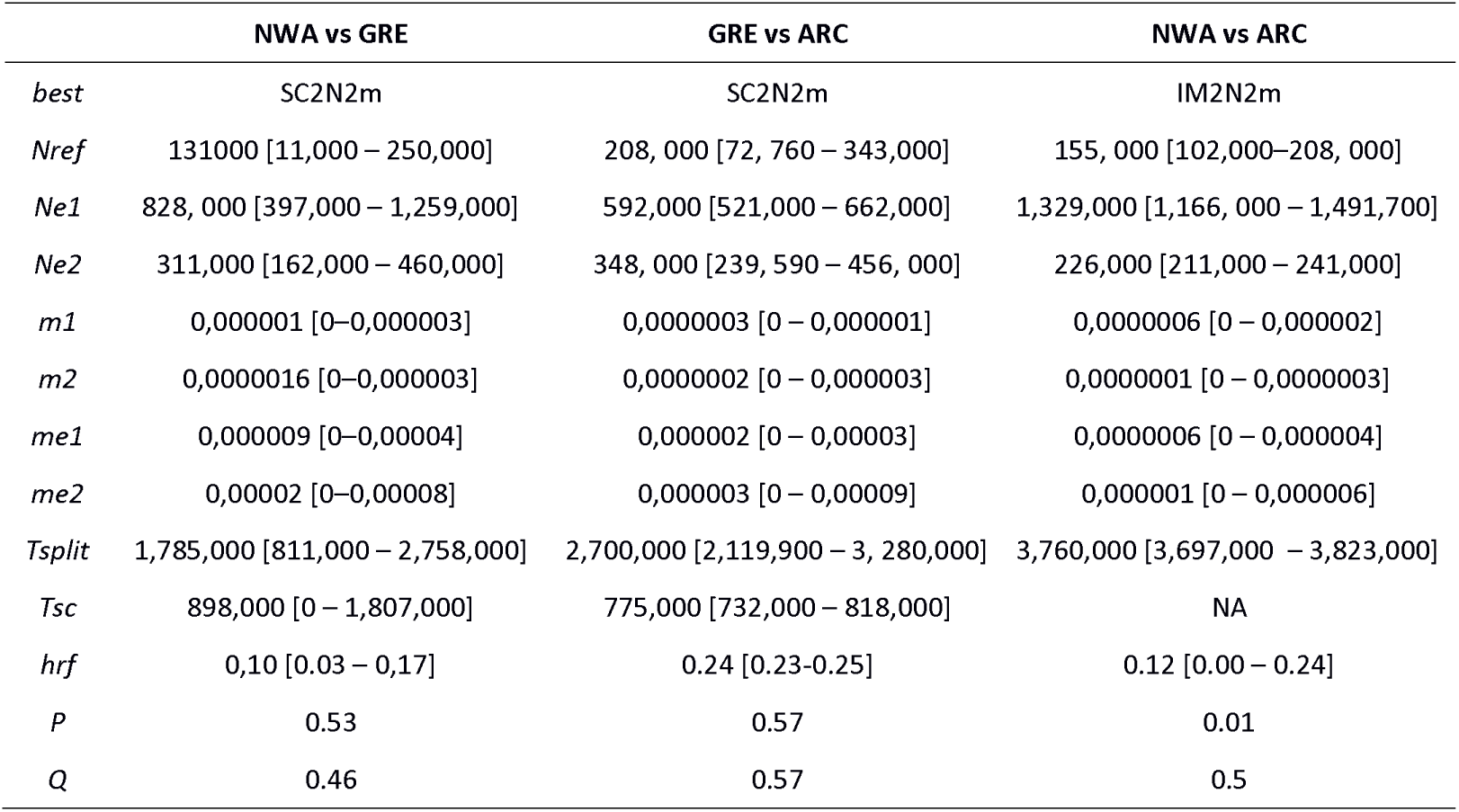
Parameter estimates under the best demographic models for each of the three lineages: Ne1 and Ne2, effective population size of the compared pair; m1 ← 2 and m2 ← 1, migration from population 2 to population 1 and migration from population 1 into population 2. me12 and me21, effective migration rate estimated in the most differentiated regions of the genome, Tsplit: Time of split of the ancestral population in the two daughter species; Tsc, duration of the secondary contact; P, proportion of the genome freely exchanged (1-P provides the proportion of the genome non-neutrally exchanged); Q, proportion of the genome with a reduced Ne due to selection at linked sites (1-Q provides the proportion of the genome neutrally exchanged); hrf = Hill-Robertson factor representing the reduction of Ne in the region (1-Q) with reduced Ne. Values in brackets provide confidence intervals around the estimated parameters.

#### Candidate loci for reproductive isolation among lineages

After excluding markers with reduced heterozygosity (He <0.10), a total of 2085, 622 and 330 markers departed from neutrality in the GRE vs ARC, NWA vs ARC and NWA vs GRE comparisons respectively. These were all randomly distributed across the reconstructed chromosomes (**Supplementary material**, **Table S7**). Of these, 276 were shared outliers between GRE *vs* ARC and NWA *vs* ARC, 130 were shared between NWA *vs* ARC and NWA *vs* GRE and 114 were shared between GRE *vs* ARC and NWA *vs* GRE, respectively. A total of seven outliers were shared in all three pairwise comparisons, such that only few outliers should be affected by background selection (see Discussion). A total of 118 outliers were fixed (i.e. *F_ST_* = 1) between GRE and ARC lineages, none were fixed in the two other pairwise comparisons and *F_ST_* values were less extreme in comparisons involving the NWA clades. A total of 688 SNPs yielded significant blast (e-value <1e^-10^ and length >90), with a total of 43 putative non-synonymous mutations. These significant hits were also randomly distributed in the genome (**Supplementary material, Table S7**) and associated with various biological processes (**see Supplementary material, Table S8**). The SNPs associated to the putative chromosomal rearrangement on Chr2 and Chr9 were significantly enriched with outliers when compared to the rest of the genome for the comparison between GRE and NWA (Fisher exact test, p < 0.0001) as well as between ARC and NWA (Fisher test, *p*-value = 0.0006) but not in the comparison between ARC and GRE (Fisher test, *p*-value = 0.7038), as expected given this rearrangement is absent in these lineages.

#### History of divergence between haplogroups within the NWA lineage

Reconstruction of the divergence history between homokaryote haplogroup1 and haplogroup3 supported a model of Secondary Contact including both linked selection and variable introgression rate among loci (SC2N2m, wAIC = 1; **Supplementary material, Table S5**). Here, it is noteworthy that all alternative versions of the secondary contact model were better than all other possible models (AM, IM, SI), even if they included linked selection or barriers to gene flow. Parameters estimates revealed that the divergence time of the group fall into the estimated divergence time for the three lineages (Tsplit ∼ 3.1 MyA CI = 2.7-3.52 MyA; **Supplementary material, Table S6**). Estimates of effective population size remained large in haplogroup1 (i.e. >500,000), but tended to be smaller than estimates for the whole NWA lineage when compared to the remaining lineages (**Table 2**; **Supplementary material, Table S6**). Haplogroup3 displayed a lower effective population size 189,000 [CI: 163,000–214,000]. Here estimates of migration rate were at least 200 times higher than between lineages, when considering the smallest difference in migration rate. This migration was strongly asymmetric with 2.6 times higher migration from haplogroup1 to haplogroup3. Finally, the proportion of the genome neutrally exchanged was approximately 95% while the proportion of the genome with a neutral effective population size was also approximately 95% (**Supplementary material, Table S6**).

### Genotype-environment associations in the NWA clade

#### Outlier loci associated to the type of spawning sites

In the pRDA, genetic variance was significantly explained by the type of spawning site (i.e., beach spawning, shallow-water demersal, and deeper-water demersal sites; overall significance, *df* = 2, *F* = 1.58, *p* < 0.001). We detected 407 and 1,737 SNPs outliers associated to the type of spawning site using pRDA (overall significance, *df* = 2, *F* = 1.12, *p* < 0.001) and LFMM, respectively (**Fig.8**). A total of 108 SNPs were detected by the two methods; i.e., 26% of the SNPs detected with pRDA were also detected using LFMM. Respectively 5% and 9% of the 108 loci were located in the genomic regions of Chr2 and Chr9 related to the haplogroups (i.e. that we defined as the genomic regions mapped on Chr2 and Chr9 and displaying high loadings on the PC1 axis of the principal component analysis performed on the NWA lineage). The outlier loci were in excess in these genomic regions compared to the rest of the genome (see the outputs of the Fisher test in **Table 3**). Furthermore, 85% and 55% of the outlier SNPs were polymorphic (allelic frequency (AF) higher than 0) in the GRE (mean AF = 0.28) and ARC lineages (mean AF = 0.26), respectively. Fisher tests (**Table 3**) indicated an excess of shared polymorphism within both lineages at outlier loci associated with the type of spawning site. We identified 112 transcripts in the 1Kbp window around the outlier loci. These transcripts were enriched in various cellular processes (e.g., bounding membrane of organelle, intracellular vesicle), biological processes (e.g., protein metabolic process, cellular macromolecule metabolic process), and molecular function (e.g., ubiquitin ligase complex, protein domain specific binding) (**Supplementary material**, **Table S9**).

**Fig. 8.**
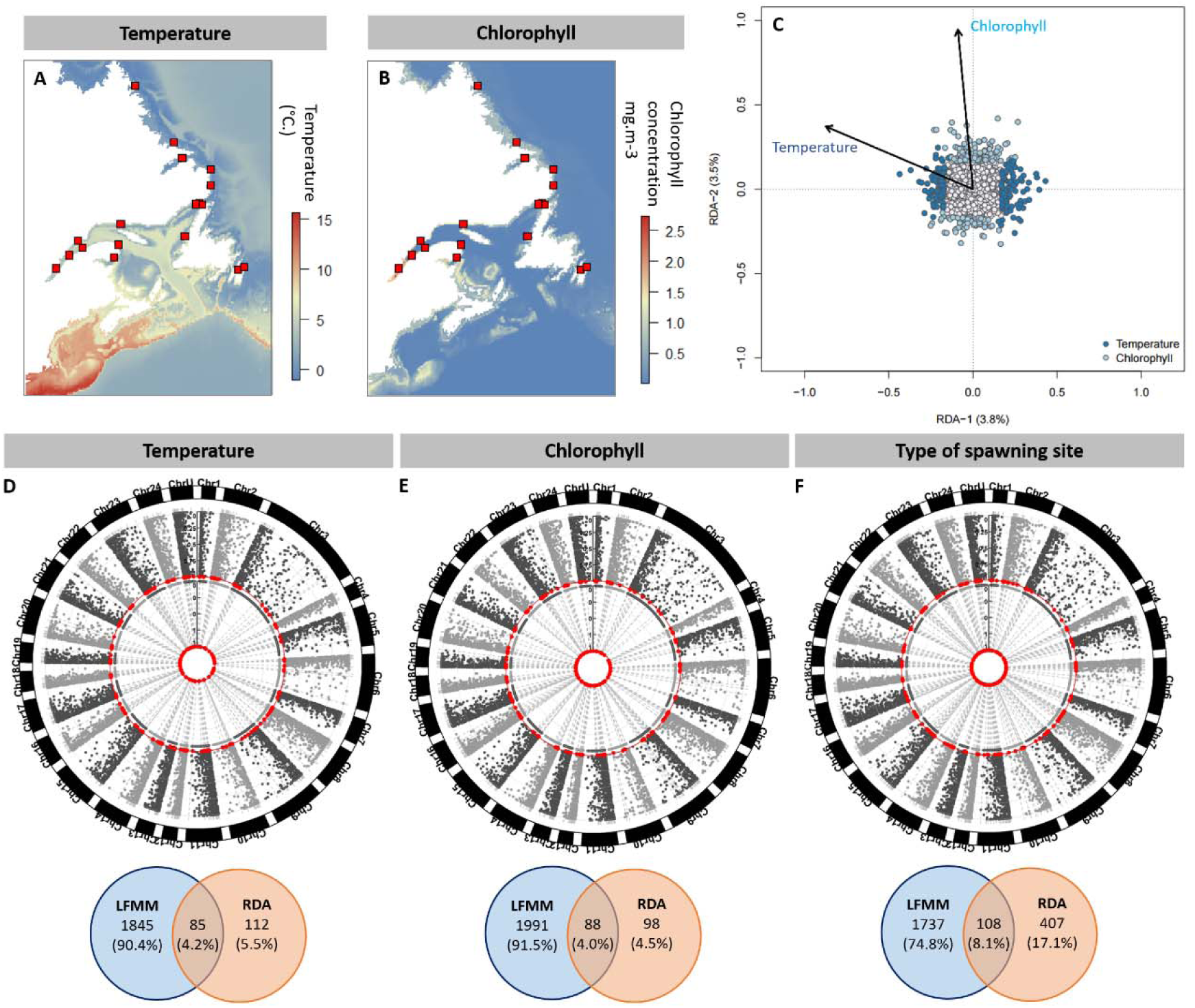
Adaptation to local environmental conditions. The two maps (A and B) shows spatial variation of temperature and chlorophyll concentration; the correlation between those two variables was relatively low (r = 0.47). RDA plot (C) highlights SNP loadings and environmental association; outlier loci are colored. (D-F) Manhattan plots shows p-values associated with SNPs outliers (shown in red) in LFMM analyses (external circle) and the evaluation of SNPs (0 = not outlier, 1 = outlier) from the pRDA (internal circle). Venn diagrams show the outlier loci detected with either LFMMs or pRDA, or both – the percentage corresponds to the proportion of outliers detected by each method and the two methods combined. Overall, our analysed showed that SNP outlier are spread throughout the genome although, for temperature and spawning-site, we detected an excess of outliers in the genomic regions contained in the putative chromosomal rearrangement (Table 3).

**Table 3.** Enrichment of SNPs outliers in the potential chromosomal rearrangement in chromosome Chr2 and Chr9 and in shared polymorphism with ARC and GRE lineages. Also presented are the outputs of Fisher tests (odd ratio and p-values).

|  | Chlorophyll |  | Temperature |  | Type of spawning site |  |
| --- | --- | --- | --- | --- | --- | --- |
|  | Ratio | p | Ratio | p | Ratio | p |
| Enrichment |  |  |  |  |  |  |
| Chromosomal rearrangement | 0.35 | 0.16 | 0.10 | <0.0001 | 0.09 | <0.0001 |
| Shared polymorphism with ARC | 0.31 | <0.0001 | 0.42 | 0.0001 | 0.31 | <0.0001 |
| Shared polymorphism with GRE | 0.10 | <0.0001 | 0.19 | <0.0001 | 0.18 | <0.0001 |
| Shared polymorphism with GRE and ARC | 0.12 | <0.0001 | 0.36 | <0.0001 | 0.22 | <0.0001 |

#### Outlier loci associated to the temperature and chlorophyll concentration within beach-spawning sites

In the pRDA, the genetic variance was significantly explained by the two environmental variables (overall significance, *df* = 2, *F* = 1.09, *p* < 0.001). A total of 197 and 1,930 SNPs outliers were potentially associated with temperature based on pRDA and LFMM, respectively (**Fig.8**). A total of 85 SNPs were identified by the two methods; i.e., 43% of the SNPs identified with pRDA were also detected using LFMM. Among those 85 SNPs, 7% and 2% respectively occurred within the genomic regions of Chr2 and Chr9 comprised in the haplogroups. Fisher tests indicated the outlier loci were in excess in these regions compared to the rest of the genome (**Table 3**). Furthermore, 76% and 47% of the SNPs displayed allelic frequency higher than 0 in the GRE (mean AF = 0.220) and ARC lineages (mean AF = 0.223), respectively. Fisher tests showed an excess of shared polymorphism for both lineages at these outlier loci (**Table 3**). Within the 1Kbp window around the outlier loci, we detected 77 transcripts that were enriched in the cellular processes (e.g., transferase complex, ubiquitin ligase complex), biological processes (e.g., regulation of growth, response to radiation), and molecular functions (e.g., ubiquitin-like protein transferase activity, ubiquitinyl hydrolase activity) (**Supplementary material**, **Table S9**).

We also identified 186 and 2,079 SNPs outliers potentially associated with the chlorophyll concentration with pRDA and LFMM respectively (**Fig.8**). A total of 88 SNPs were identified using the two approaches; i.e., 47% of the SNPs detected with pRDA were also detected using LFMM. Among those 88 SNPs, 2% and 5% were present in the genomic regions Chr2 and Chr9 comprised in the haplogroups. However, the outlier loci were not found to be in excess in these genomic regions (**Table 3**). Furthermore, 91% and 55% of the SNPs displayed allelic frequency higher than 0 in the GRE (mean = 0.26) and ARC lineages (mean = 0.23) respectively. Fisher tests showed an excess of shared polymorphism for both lineages in outlier loci (**Table 3**). We detected 93 transcripts within the 1Kbp window around the outlier loci. These transcripts were enriched in the cellular processes (e.g., cyclin-dependent protein kinase holoenzyme complex, intracellular membrane-bounded organelle), biological processes (e.g., swim bladder inflation, cell cycle phase transition), and molecular functions (e.g., kinase regulator activity) (**Supplementary material**, **Table S10**).

## Discussion

Our study revealed a complex demographic history of divergence among the three capelin glacial lineages in North Atlantic and Arctic oceans. Models of secondary contact including both linked selection and variable introgression rates among loci received highest statistical support. They showed that the lineages diverged from 3.8 to 1.8 MyA depending on the pair of clades considered, and we identified loci potentially involved in their reproductive isolation in absence of physical barrier to gene flow between them. These observations support the hypothesis that these lineages may in fact represent cryptic species of capelin or at the very least they indicate that these lineages are part of an ongoing speciation process. Within lineages, our analyses revealed large *N_e_* and a low genetic divergence among sampling sites at both historical (*F*_ST_-based approach) and contemporary (clustering method) time scales. In the NWA lineage, a likely chromosomal rearrangement that appeared early during the inter-lineage divergence process resulted in the existence of three haplogroups spread in most spawning sites of the NWA lineage but showing differences in frequencies between capelin from different spawning habitats. Genotype-environment associations revealed molecular signatures of local adaptation to local sea conditions prevailing at spawning sites. Importantly, our study emphasized that shared polymorphism among glacial lineages and chromosomal rearrangements occurring in the NWA clade broadly contributed to the genetic polymorphism involved in local adaptation in the presence of high gene flow.

### An ongoing speciation process among lineages

Our modelling approach supports the hypotheses that i) the three lineages have diverged in allopatry and accumulated barriers to gene flow resulting in heterogenous differentiation, and ii) that such differentiation is also driven by linked selection either in the form of background selection (Charlesworth et al. 1993) or selective sweep and hitchhiking (Smith & Haigh 1974). Overall, demographic reconstruction suggests a relatively long period of divergence, compatible with a divergence level comparable to those observed between species (Hedges et al. 2015, Roux et al. 2016). This divergence has resulted in strong contemporary genome-wide differentiation randomly distributed across scaffolds. Accordingly, we found that candidate genomic outliers for reproductive isolation were spread randomly along the chromosomes, as expected under the genic view of speciation and gradual accumulation of Dobzhansky-Muller incompatibilities (DMI) (Coyne & Orr 2004, Wu 2001). Under this model, the majority of barriers are endogenous, and exogeneous barriers (i.e. environmental barriers) are expected to have a minor role in the reproductive isolation of these lineages (Bierne et al. 2011). This divergence has apparently resulted in strong barriers to gene flow, explaining very low current levels of gene exchange among lineages, despite the complete absence of physical barriers to gene flow. To our knowledge such high levels of restricted gene flow between parapatric lineages have rarely been documented in a strictly marine species (Le Moan et al. 2016, Gagnaire et al. 2018).

The statistical support for the model of divergence with gene flow (IM) with heterogeneous drift and migration (i.e. IM2N2m model) between NWA and ARC. Under this model, we found that levels of gene-flow were close to zero with less than one migrant being exchanged every generation (Table 2). In such conditions, this model is very close to a model of strict isolation. An alternative, but non-exclusive explanation is that the isolation signal may have been lost if the period of secondary contact was too long relative to the period of isolation. For instance, here, the timing of secondary contact represented 20% of the total divergence time, in the two other model comparisons. This long period of gene-flow is above the ideal conditions under which the secondary contact signal can still be detected (Alcalla & Vuilleumier 2014, Roux et al. 2016).

Separating the analysis by haplogroups and excluding Chr2 and Chr9 for the NWA lineage indicated that our results were robust to putative confounding effects by a potential chromosomal rearrangement. This separate analysis, together with the confidence intervals around our parameter estimates suggested that the three lineages diverged during the Pleistocene glacial oscillations, as proposed by Dodson et al. (2007) and as observed in other large-scale comparison in Atlantic salmon (*Salmo salar*) (Rougemont & Bernatchez, 2018), and in the mussel species complex *Mytilus galloprinciallis* and *Mytilus edulis* (Roux et al. 2014). The ARC and NWA diverged first, followed by a split of the GRE from the ARC lineage. We hypothesized that a more recent split time between NWA and GRE could result from 1) an underestimation of the divergence time between NWA and GRE – GRE diverged approximately at the same time from both the NWA and ARC, as supported by overlapping confidence intervals (Table 2). Alternatively, (2) the observed net levels of sequence divergence (lower between NWA and GRE), the PCA (**Supplementary material**, **Fig. S10**), and the more recent divergence could also be compatible with the hypothesis of non-modelled ancestral introgression events between the two lineages that could globally result in lower divergence than expected.

The action of selection and the accumulation of barriers to gene flow are not the only factors driving heterogeneous differentiation along the genome. Our modelling approach indicated that approximately 45-50% of the genome was undergoing linked selection (Table 2) and that effective population size (*N_e_)* was locally reduced to approximately 10-20% of its size due to Hill-Robertson interference (Table 2). The role of such a linked selection process, unrelated to speciation, is increasingly recognized as being implicated in observed patterns of genome-wide divergence (Cruickshank & Hahn 2014, Burri 2017), and has been empirically demonstrated elsewhere (e.g., Burri et al. 2015, Roux et al. 2016, Wang et al. 2016, Rougemont & Bernatchez, 2018, Stankowski et al. 2019). Admittedly, further estimates of recombination rate variation along the genome will be needed to help disentangling how linked selective processes are currently affecting genome-wide divergence patterns. Here, the inference that 50% of the genome can be affected by this process should be kept in mind when interpreting putative outliers inferred from the approaches below, which might be highly affected by these confounding factors (Lotterhos 2019).

### A potential chromosomal rearrangement in the NWA lineage

Our PCA analyses highlighted the existence of three haplogroups spread into almost all sampling sites of the NWA lineage which are differentiated only at limited number of co-localized SNPs in the genome. This result suggests the presence of a chromosomal rearrangement with a lower recombination, although PCA is not the most effective tool to detect structural variants (Li & Ralph 2019; but see Ma & Amos 2012). Indeed, the signal left by inversions cannot easily be distinguished from long haplotypes under balancing selection or simply regions of reduced recombination (Lotterhos 2019). Here however, our synteny analysis indicates that the loci included in the putative chromosomal rearrangement are located in Chr2 and Chr9 of the *Esox lucius* genome. The PCA executed for Chr2 and Chr9 separately also displayed the same haplogroup pattern. Assuming that Chr2 and Chr9 are completely fused in the capelin, this strongly suggests the existence of a polymorphic chromosomal inversion in the NWA lineage.

Our analyses indicated that the chromosomal rearrangement is monomorphic (i.e., absence of the three haplogroups) in ARC and GRE lineages. Given the large *N_e_* of these lineages, it is unlikely that the rearrangement has lost its polymorphism due to the effect of genetic drift during lineage divergence. The most probable hypothesis is that the rearrangement appeared in the NWA lineage ∼ 2.8 MyA after the split from the ARC lineage but almost concomitantly with the GRE split from the NWA. Then, the rearrangement would have spread into the NWA lineage, perhaps during *N_e_* reduction events associated with the first glaciations of the Gelasian stage (2.6 MyA) (Rio et al. 1998). Gene flow and introgression between NWA and GRE lineage would explain the fact that genetic differentiation between haplogroup 1 and haplogroup3 was higher than the differentiation between each haplogroup and the GRE lineage. Further data and modeling will be required to formally support or reject this hypothesis.

The three haplogroups occur at variable frequency among sampling sites, with haplogroups 1 and 2 being most prevalent (on average 57 and 38% of the individuals per site respectively). By contrast, haplogroup 3 was much less common (5% of the individuals) and was absent from a small number of sampling sites. Given the large effective size and high gene flow observed in capelin, such widespread polymorphism may be neutral and/or transient. Yet, large chromosomal rearrangements do not easily get established or maintained because they prevent the purge of deleterious mutations, and thus polymorphism patterns have frequently been associated to balancing selection, either due to overdominance, frequency dependence or spatially-varying selection (Kirkpatrick 2010, Guerrero & Hahn 2017, Wellenreuther & Bernatchez 2018, Faria et al. 2019). In the case of the capelin, the absence of heterozygote excess over Hardy-Weinberg expectation for the putative heterokaryote (haplogroup2) does not suggest any overdominance effect. Yet, the fact that an excess of outlier markers was found within the rearrangement and that the frequency of haplogroups depends on the type of spawning sites suggest that selection may contribute in maintaining this polymorphism. Future work addressing the genetic architecture of this putative rearrangement, its ancestry, its distribution and investigating its possible association with phenotype would be needed to better understand its implication in the evolution of the NWA lineage.

### High gene flow among spawning sites within lineages

Our study highlighted a tenuous, yet significant genetic variation within NWA lineage. These results are also congruent with many studies on marine organisms (excluding reef species) that reported low genetic variation across large geographic areas (e.g., fishes: Lamichhaney et al. 2012; crustaceans: Benestan et al. 2015; molluscs: Van Wyngaarden et al. 2017; for a review, see Kelley et al. 2016). The weak intra-lineage genetic differentiation within NWA partly results from large historical effective population sizes in the three lineages – several million individuals according to our demographic inferences. Furthermore, the low values of pairwise *F*_ST_ and the slight IBD detected using neutral markers both indicate high dispersal among spawning sites that led to pronounced historical gene flow. However, we detected slight differences in *F*_ST_ values among the three types of breeding modes: *F*_ST_ were higher between demersal and beach spawning sites than between the two types of demersal sites (i.e., shallow and deep-water sites). This pattern suggests a more restricted gene flow between demersal and beach spawning capelin than within demersal spawning capelin. In addition, our analyses revealed that the frequency of haplogroup 3 was higher in demersal spawning sites than in beach spawning sites. Altogether, these results suggest high but non-random gene flow among spawning environments, resulting in a fine-scale genetic structure in the NWA lineage.

### Habitat matching choice and adaptation to local marine conditions

Beyond the weak genetic structure at the scale of the NWA range, our results suggest that the type of spawning site (beach site, demersal shallow-water site, and demersal deep-water site) is an important predictor of genetic variation at loci under putative divergent selection. As adult capelins were captured at spawning sites during spawning, this pattern likely results from a habitat match choice (Edelaar et al. 2008). This mechanism implies that individuals disperse towards reproductive sites that maximize their breeding performances, which allows local adaptation despite high gene flow (Edelaar et al. 2008, Jacob et al. 2017). As individual habitat choice does not always match optimal environment conditions (i.e., partial habitat matching) or because a population may contain both specialist and generalist individuals (Jacob et al. 2017), it may result in a weak genetic structure due to large effective population size and/or gene flow between the two habitats.

The choice of spawning site taking place during capelin pre-spawning migration could be genotype-dependent to maximize offspring fitness at embryonic and larval stages under particular environmental conditions. The spawning sites of beach-spawning capelin located in the intertidal zone are exposed to a higher environmental stochasticity than those of demersal-spawning fishes. Highly variable meteorological factors such as solar radiation and wind strength and direction broadly affect embryonic and larval growth and survival as well as swimming performance during larval dispersal (Frank & Leggett 1981a, 1981b, 1982; Leggett et al. 1984). In addition, beach spawning sites display more variable and higher water temperatures than demersal sites (Nakashima & Wheeler 2002), which may sometimes lead to dramatic mortality events before larval hatching (Leggett et al. 1984). Because of the stressful environmental conditions in the intertidal zone, survival at early stages is negatively correlated with the time spent by eggs and larvae in this area (Frank & Leggett 1981a). Interestingly, a previous study has suggested that embryos from beach-spawning capelin have a faster growth rate than those of demersal capelins (Nakashima & Wheeler 2002), which might indicate local adaptation allowing an accelerated development in the intertidal zone. In parallel, our results indicate that candidate loci potentially under divergent polygenic selection and associated to the type of spawning site are involved in a broad range of biological processes (**Supplementary material**, **Table S9**) driving cell resistance to environmental stress. Taken together, these results suggest the existence of a habitat matching choice and an adaptation to local environmental conditions prevailing at beach and demersal spawning sites.

Our study also suggests the existence of a similar mechanism of adaptation to the conditions of temperature and trophic productivity experienced in beach spawning sites. We detected two markers under putative divergent selection associated with water temperature and localized in genomic regions (a narrow window of 1 Kbp) that contain genes involved in growth regulation (thermo-dependent in fishes, Pauly 1980) and response to UV radiation (Supplementary material, Table S4). Furthermore, a locus associated with trophic productivity is implicated in the expansion of fish swim bladder, an organ playing a critical role in buoyancy, movement behavior, and energy expenditure (Woolley & Qin 2010). An increased buoyancy control by an inflated swim bladder contributes to reduce energy expenditure related to movement and increases larval survival during dispersal (Woolley & Qin 2010), which could be particularly beneficial in limited food environments. Overall, these results are consistent with previous studies on capelin showing that water temperature and trophic productivity strongly affect embryonic and larval growth and survival, as well as the swimming capacity of larvae during the drift phase following hatching (Frank & Leggett 1981a, 1981b, 1982; Leggett et al. 1984). Taken together, these studies and ours strongly suggest that temperature and trophic productivity are important selective agents driving local adaptation in capelin, as reported in other marine fishes (Bradbury et al. 2010), crustaceans (Benestan et al. 2016), and molluscs (De Wit & Palumbi 2013).

### Contribution of genomic background to local adaption

Our analyses showed that standing genetic variation likely plays a central role in local adaptation to sea conditions, as predicted by theory (Barrett & Schluter 2008, Hedrick 2013). Indeed, 85% and 55% of the markers putatively under divergent selection and associated with the type of spawning site were polymorphic in GRE and ARC lineages, respectively. In addition, 76% and 47% of the candidate outliers related to temperature, as well as 91% and 55% of the outliers associated with trophic productivity, were also polymorphic in GRE and ARC lineages respectively. Importantly, we consistently found an excess of shared polymorphism with both lineages in this set of outlier loci. Therefore, these results strongly suggest that adaptation to local environmental condition is facilitated by a high level of shared polymorphism among glacial lineages that was maintained over time by large effective population size and high gene flow among reproduction sites across the distribution range of each lineage.

Our analyses also indicated that a non-negligible proportion of outlier markers associated with the type of spawning site (8%), temperature (7%), and trophic productivity (2%) is located in the putative chromosomal rearrangement. For both the type of spawning site and temperature, the outlier loci were in excess in the genomic regions contained in the chromosomal rearrangement compared to the rest of the genome. These results are thus consistent with previous work highlighting the important role of chromosomal polymorphism in genome and phenotype evolution, as well as in local adaptation in marine fishes (Roesti et al. 2015, Sodeland et al. 2016, Barth et al. 2017, Lehnert et al. 2019). In particular, studies showed that chromosomal arrangements may be involved in migratory behavior (Pearse et al. 2014, Kirubakaran et al. 2016, Berg et al. 2017) and habitat selection (Barth et al. 2019). In parallel, other works highlighted that chromosomal polymorphism may facilitate adaptation to local conditions including temperature despite high gene flow (Barth et al. 2017, Wellband et al. 2019).

### Conclusion

Our study shows that a thorough understanding of the interaction of divergence processes and chromosomal polymorphism allows to better understand processes of local adaptation in species with high dispersal ability. It also emphasizes the importance of quantifying the relative contribution of intra- and inter-lineage gene flow, historical population size, selection, and chromosomal rearrangement in the retention of genetic diversity in marine species. Improving our knowledge about those processes is critical toward better evaluating and predicting the evolvability as well as the probability of evolutionary rescue in marine organisms in a context of climate change and global increase of anthropogenic pressures.

## Acknowledgments

We thank biologists and technicians of the Department of Fisheries and Oceans Canada for their implication as well as all everyone who contributed to sampling throughout the study area. This research was funded by a Strategic Project Grant from the Natural Sciences and Engineering Research Council of Canada (NSERC) to L. Bernatchez, M. Clément and P. Sirois, a financial contribution of Ressources Aquatiques Québec and was also supported by in-kind contribution from many other organisations: Department of Fisheries and Oceans Canada, Nunatsiavut Government, NunatuKavut Community Council, Labrador Fishermen’s Union Shrimp Company, Department of Fisheries and Aquaculture – Government of Newfoundland and Labrador, World Wildlife Fund Canada, St. Lawrence Global Observatory, Parc Marin du Saguenay–Saint-Laurent, and the Greenland Institute of Natural Resources. Whole genome sequencing and construction of the draft capelin genome assembly were funded by the Research Council of Norway (RCN) through the Nansen Legacy project (RCN no. 276730) and the ComparaCod project (RCN no. 222378). PacBio library creation and high throughput sequencing were carried out at the Norwegian Sequencing Centre (NSC), University of Oslo, Norway. Genome assembly was performed on the Abel Supercomputing Cluster (Norwegian metacenter for High Performance Computing (NOTUR) and the University of Oslo) operated by the Research Computing Services group at USIT, the University of Oslo IT-department (http://www.hpc.uio.no/).

## Author contributions

H.C. and Q.R. made the statistical analyses and wrote the paper. M.L., C.M., E.N., and Y.D. contributed to the bioinformatics and statistical analyses. L.B., P.S., M.CA., and M.CL. initiated the project, and L.B. conceptualized and coordinated the work. T.J. collected the DNA samples in the GRE lineage. The generation of the draft capelin genome assembly was initiated and coordinated by S.J, and samples provided by K.P. The whole genome sequencing DNA extraction was performed by S.N.K.H and construction of the draft genome assembly was done by O.K.T and S.N.K.H.

