## Supplementary_material for "Standing genetic variation and chromosomal rearrangements facilitate local adaptation in a marine fish"

Hugo Cayuela*, Quentin Rougemont* et al. **Standing genetic variation linked to ancestral polymorphism and chromosomal rearrangement facilitates local adaptation in a marine fish**

**Supplementary material**

**Filtering procedure for demographic inference with δaδi**

The full dataset was filtered separately to obtained unbiased site frequency spectrum to performed demographic inferences. A minimal filtering was performed to avoid biasing our inferences. From the full vcf containing 550,724 SNPs, we kept one SNP per locus and further filter the vcf to remove long distance LD by keeping SNPs with r^2^ < 0.2 using plink (option --indep-pairwise 50 10 0.2). For the GRE and ARC lineages, all samples sites were merged and considered as a single population. For the NWA lineage we combined a set of four sample sites, showing low Fst (<0,0030) and low missing rates, in order to obtain similar number of individuals as for the GRE and ARC. We then splitted the vcf by population and remove any SNPs with more than 10% of missing data and excluded SNPs departing strongly from HWE (p-value of 0.0001) to remove putatively remaining paralogs. We then remove non-polymorphic markers between any pair of populations and subsampled all our pairwise site frequency spectrum in δaδi to remove missing data. The final number of SNPs ranged from 9,500 to 17,000 depending on the pairs. To estimate demographic parameters we estimated the full length of sequenced RAD loci and corrected it for the final number of retained SNPs. All site frequency spectrum will be available on dryad.

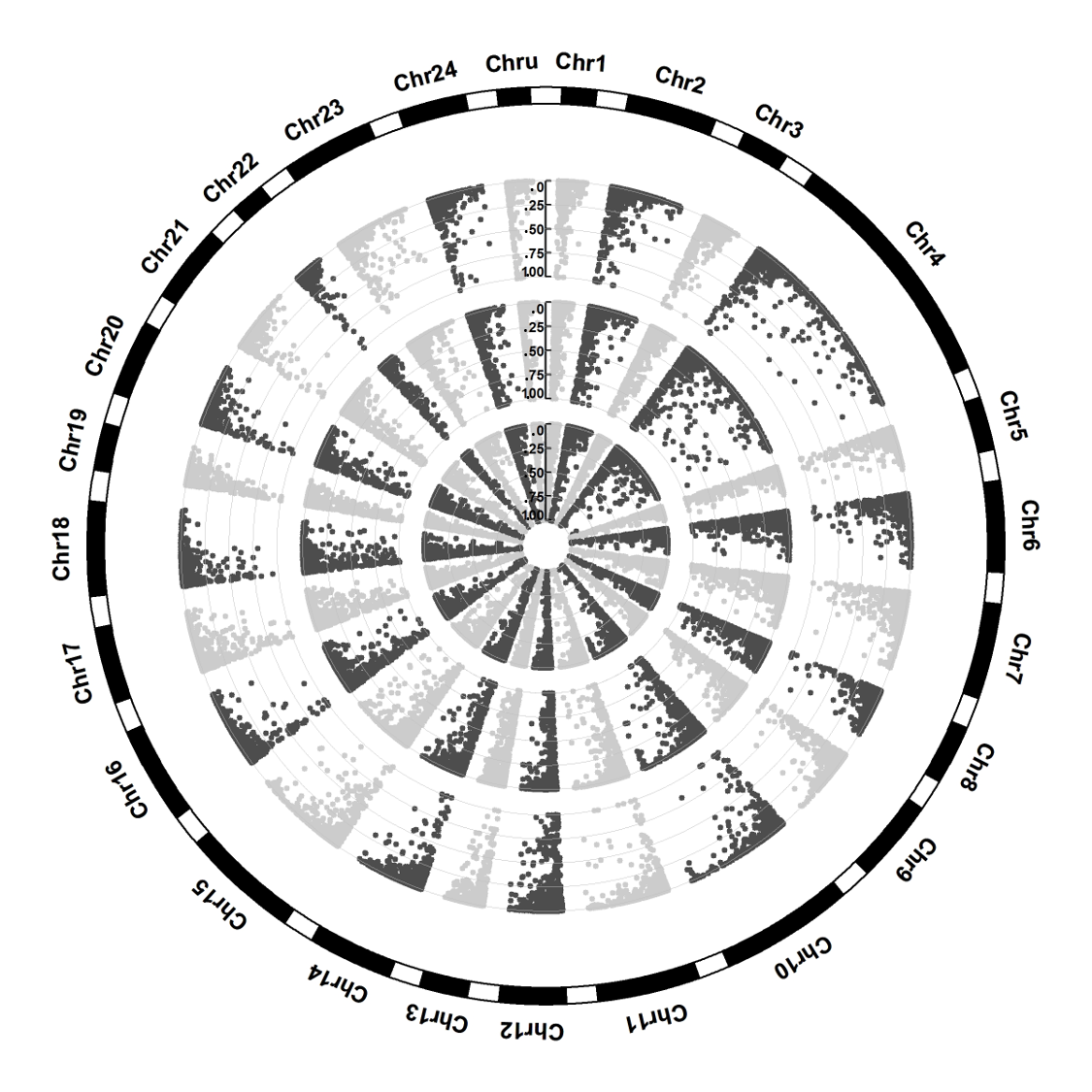

**Fig. S1**. Manhattan plots of the $F_{ST}$ values between the three lineages (ARC, NWA, and GRE) of capelin. The outer circle shows the $F_{ST}$ between ARC and GRE lineages. Medium circle presents the $F_{ST}$ between ARC and NWA lineages. Central circles show the $F_{ST}$ between GRE and NWA lineages. The 24 ancestral chromosomes are shown at the external circle, and contigs that were not assigned to a chromosome were aggregated in the group ChrU.

**Fig. S2**. Heatmap showing the proportion of each contig of capelin (*Mallotus villosus*) that aligns, at the protein level, with the four chromosome-scale genome assemblies analysed. Each line represents one *M. villosus* contig. Each column represents a chromosome in the reference genomes, with letters referring to the compared species, *Esox lucius* (E)*, Dicentrarchus labrax* (D)*, Sparus aurata* (S)*,* and *Takifugu rubripes* (T), and with numbers (1-24) referring to the arbitrary chromosome names and correspondence given in table S1. U refers to scaffolds from the four reference genomes that were not assembled into chromosomes. Note the high level of synteny between the genome of *Mallotus villosus* and the genomes of *Esox Lucius* and *Dicentrarchus labrax*.

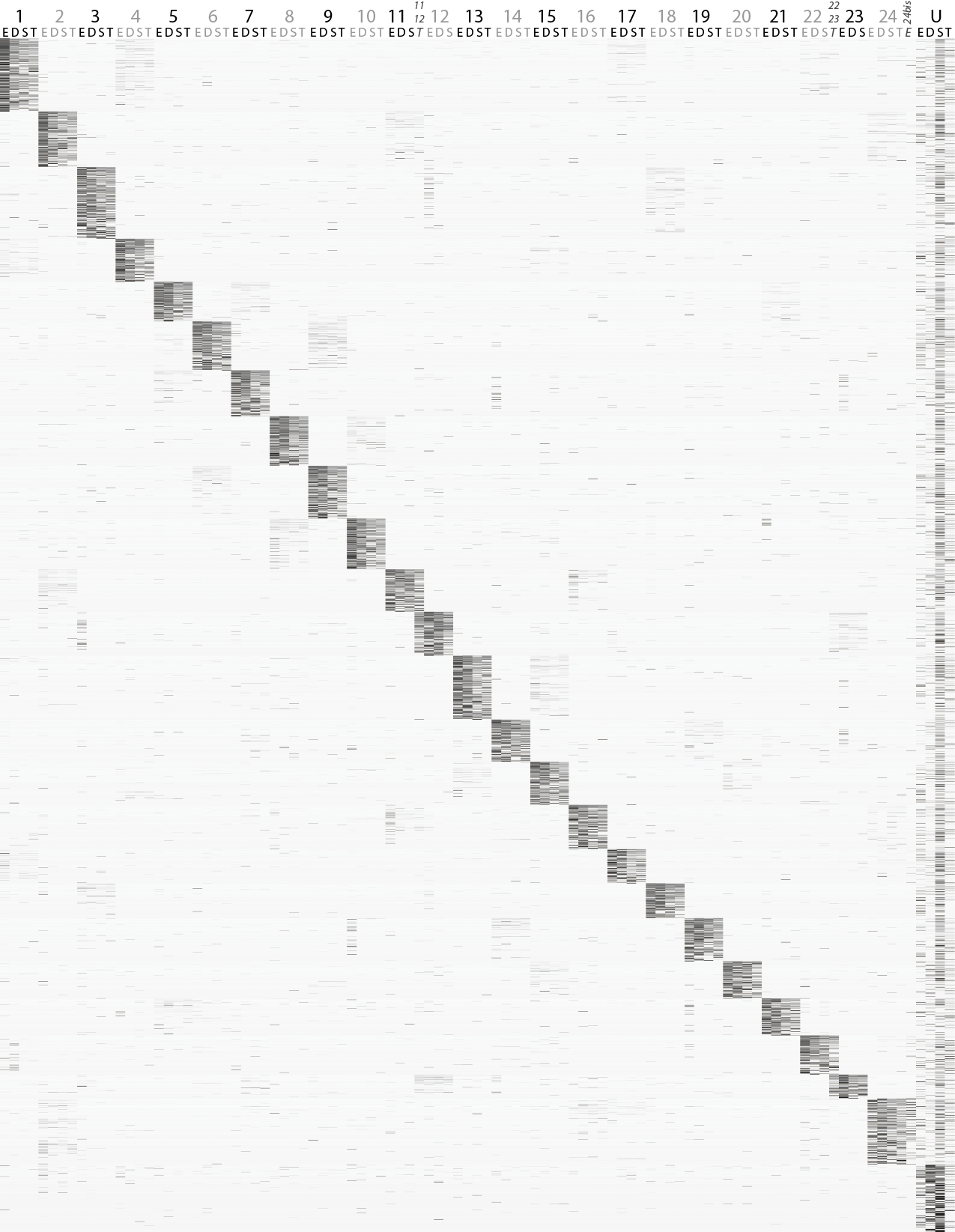

**Figure S3.** Compared demographic models for capelin. Four major demographic models are compared, a model of strict isolation (SI), a model of Ancient Migration (AM) a model of Isolation with Migration (IM) and a model of Secondary Contact (SC). Each model allows for heterogeneous effective population size (as depicted by the variable coalescent width). Models with gene flow (AM, IM, SC) further allow for heterogeneous migration rate to account for the accumulation of barrier to gene flow during the divergence process.

**Figure extracted from Rougemont & Bernatchez (2018).**

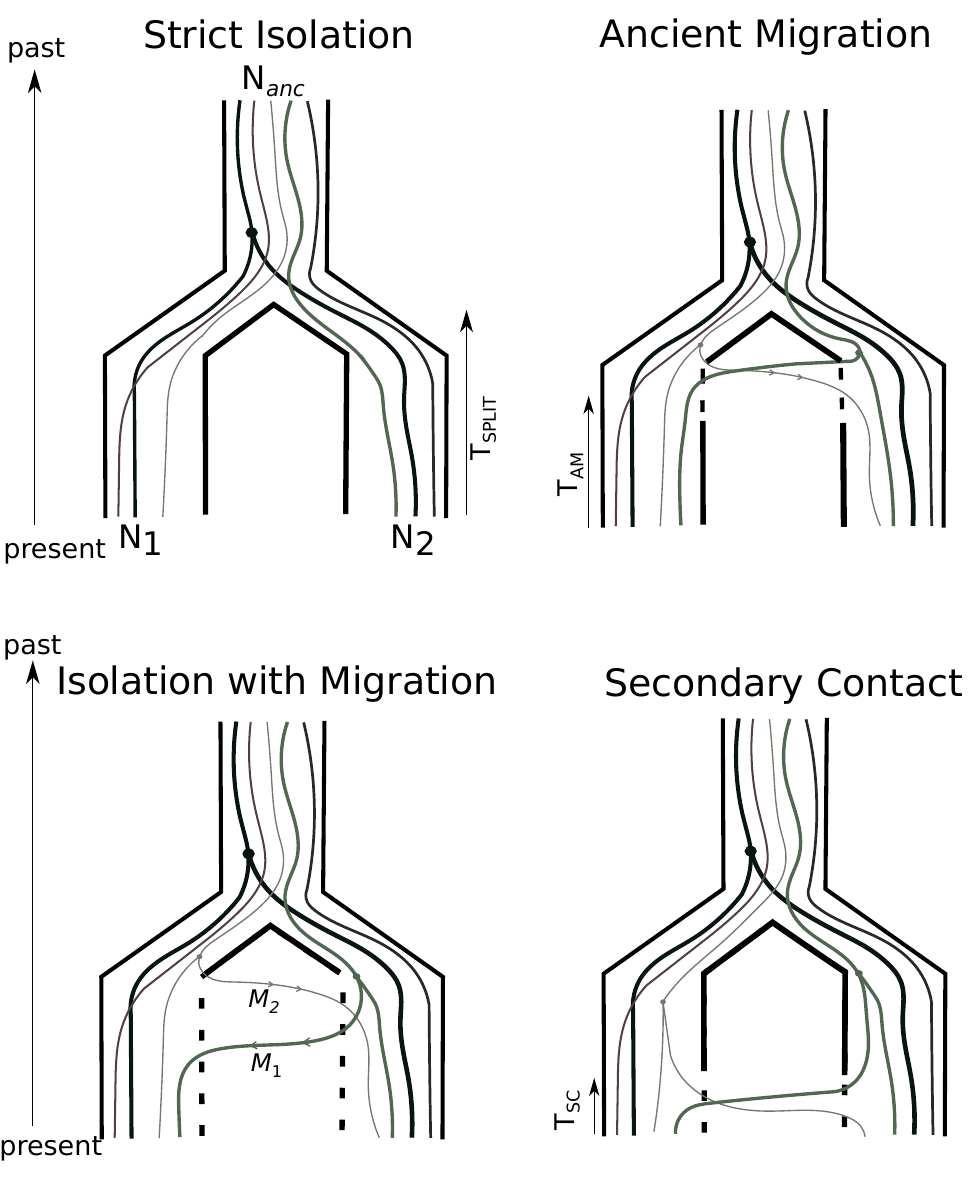

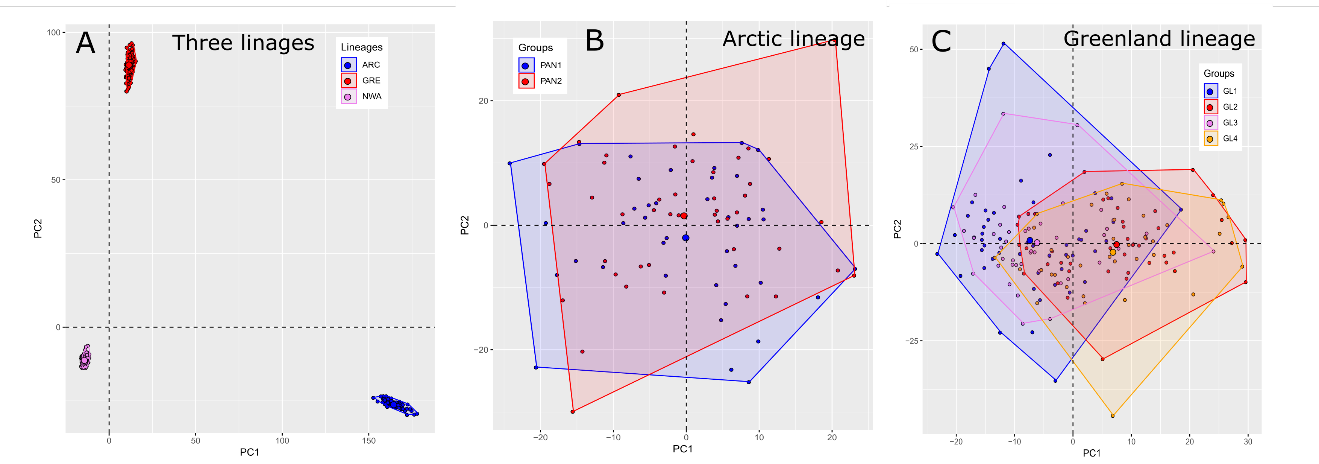

**Fig.S4**. Genetic structure and differentiation among and within the three lineages (NWA, GRE, and ARC) of *Mallotus villosus*. (A) Principal Component Analysis (axis 1 and 2) showing the genetic variation among the lineages. Principal Component Analysis showing the genetic variation within the arctic (B) and Greenland (C) lineages. Two and four sampling sites were respectively considered in the Arctic lineage (PAN1 and PAN2) and the Greenland lineage (GL1, GL2, GL3, and GL4).

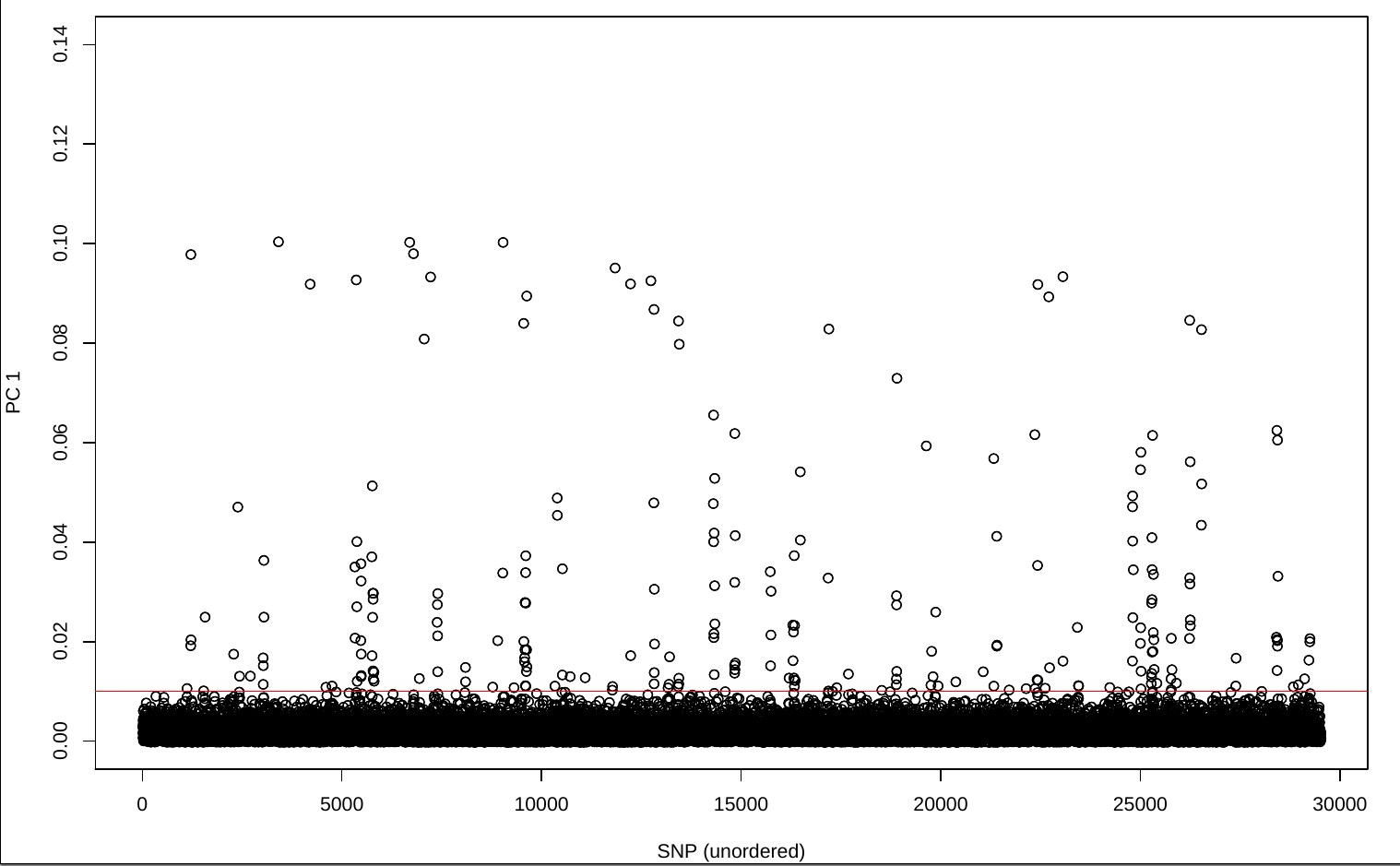
**Figure S5:** SNPs loading along the first axis discriminating three groups. SNPs with the highest loadings (load > 0.10 red line) were mapped preferentially to the chr2 and chr9.

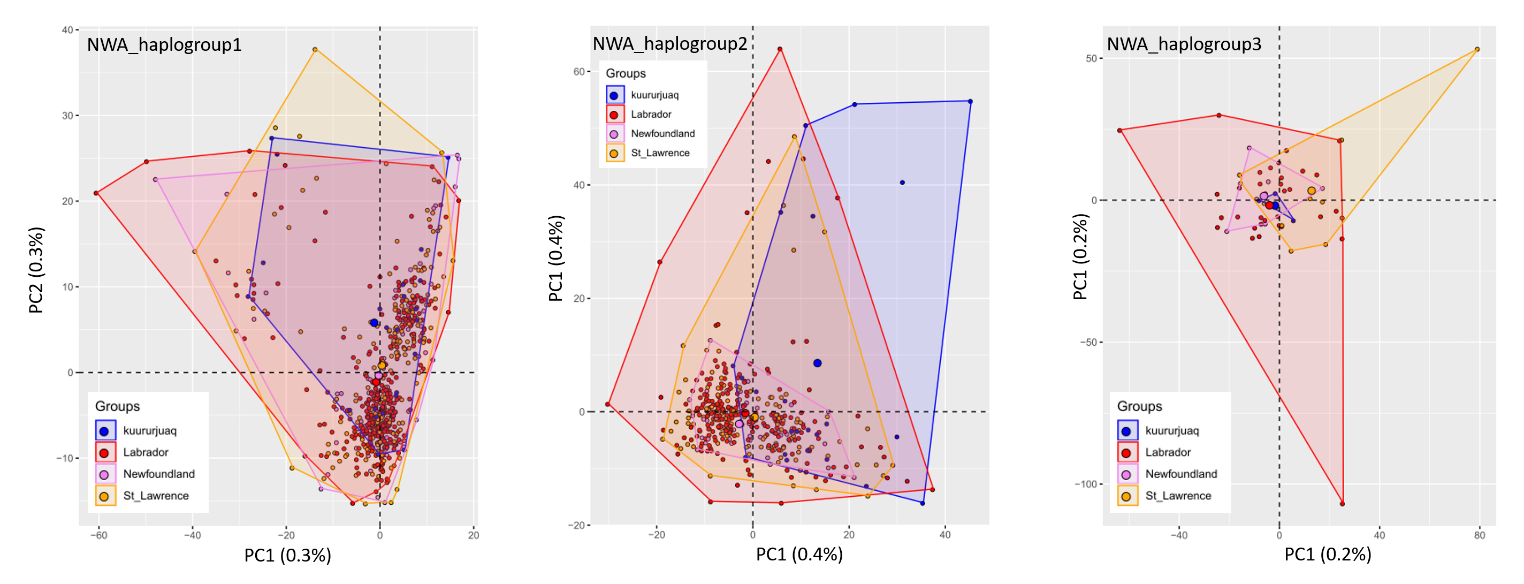

**Fig.S6**. Genetic structure and differentiation within the three haplogroups within the Northwest Atlantic (NWA) lineage of *Mallotus villosus*.

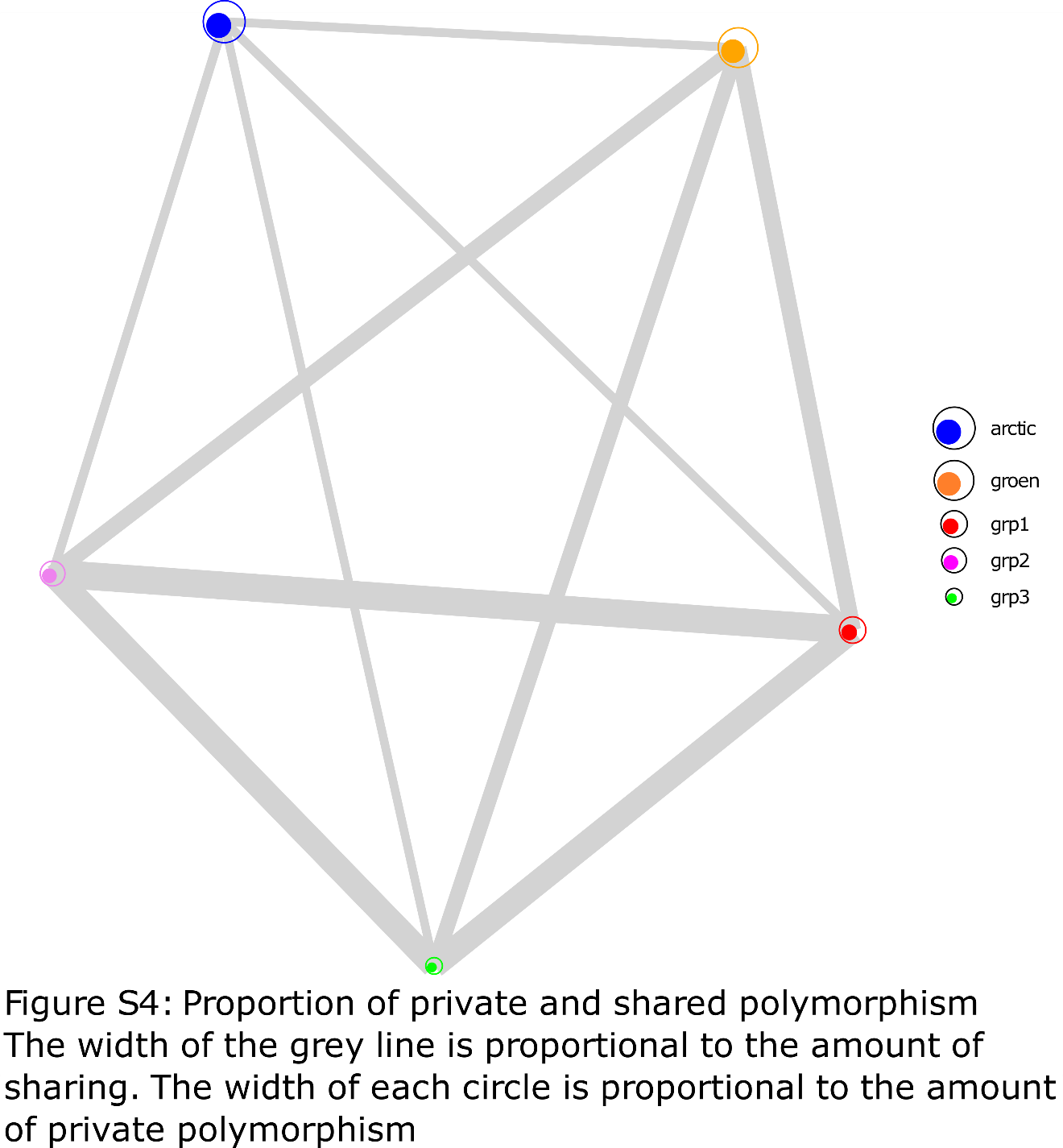

**Fig.S7**. Proportion of private and shared polymorphism in the Arctic and Greenland lineages, and in the three haplogroups (grp1, grp2, and grp3) of northwest Atlantic lineage (NWA). The width of the grey lines is proportional to the amount of polymorphism sharing. The size of the circles is proportional to the amount of private polymorphism.

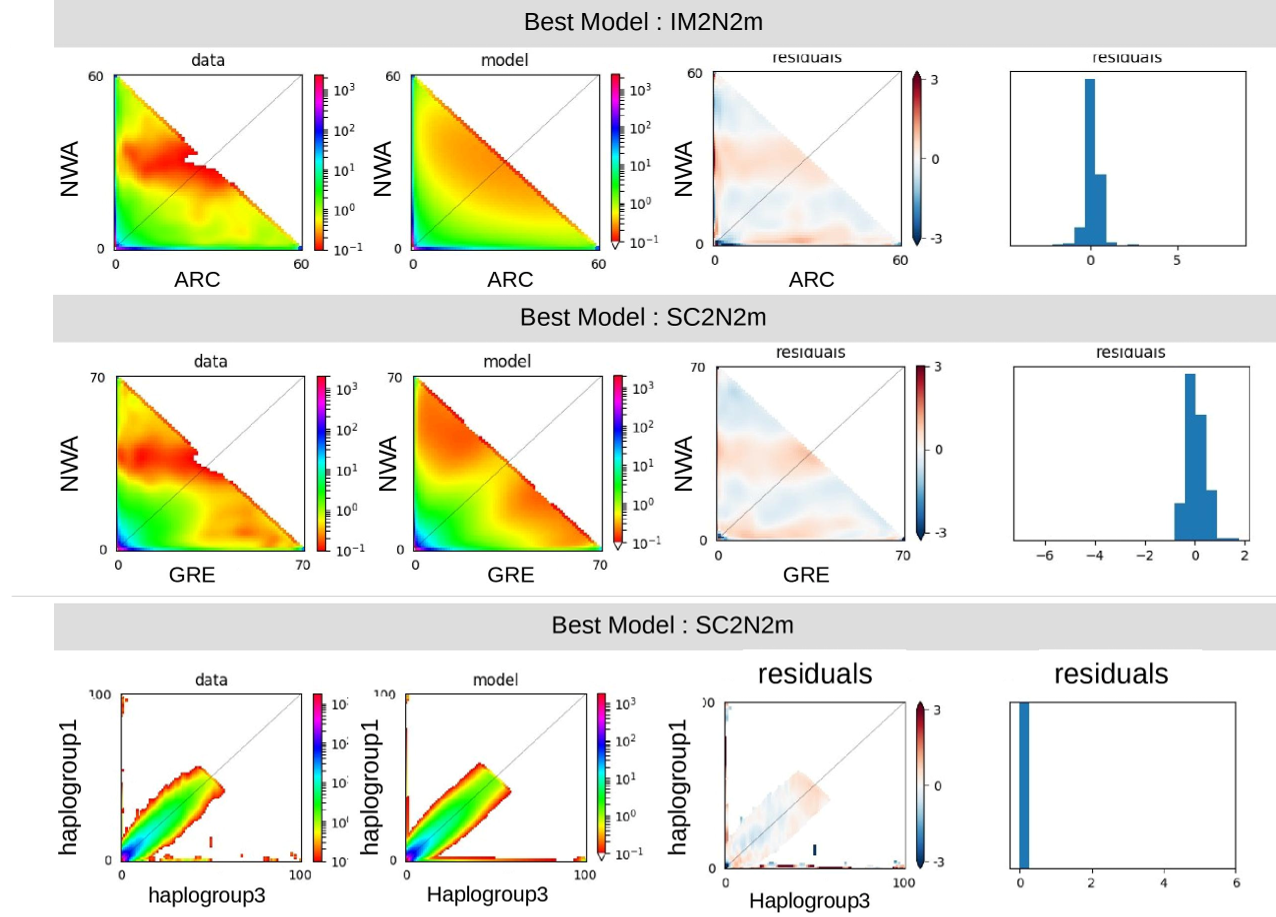

**Fig. S8.** Observed and fitted JSFS between NWA and GRE and ARC excluding the rearrangement and JSFS between haplogroup 1 and haplogroup 3 under the best model along with the residuals.

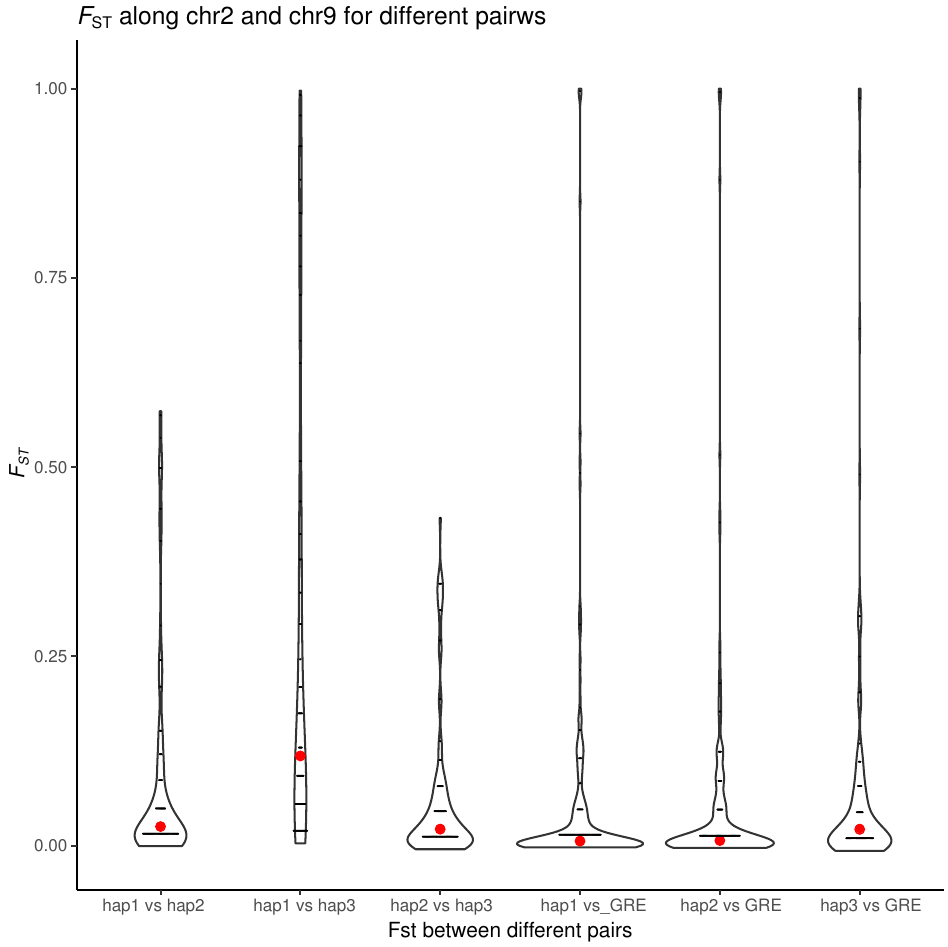

**Fig. S9.** Violin Plot displaying Weir & Cockerham (1984) Fst estimator for each 212 SNPs of the Chr2 and Chr9 with high loading along axis 1 of the PCA. Fst values between haplotgroup are depicted in violin plot 1-3 (lef) and Fst values between haplogroups and the GRE lineage are depicted in violin plot 4-6 (rigth part). The red dots represent the median Fst value. The Fst is lower between GRE and the haplogroup(s) than among haplogroup.

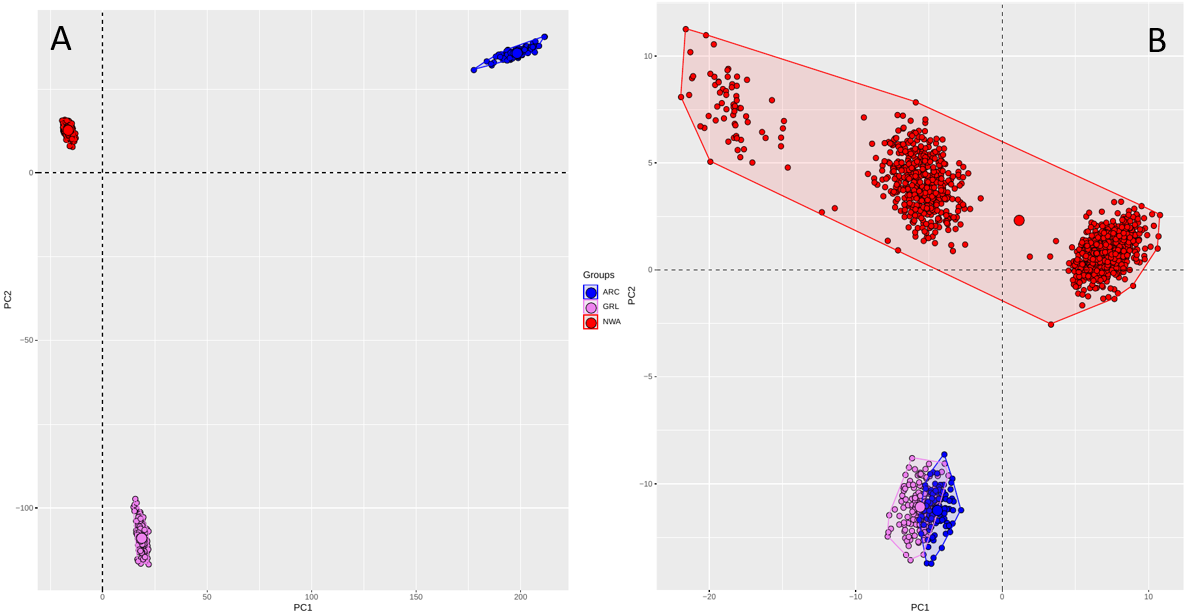

**Figure S10.** A) PCA including all the SNP (25904) except the 212 SNPs with highest loadings on PC1 of figure 3A. The distance among lineages are increased as compared to the PCA with all data (Fig 2A). B) PCA with only the SNP with the highest loading in Figure 3A but including all three lineages (n = 1474 individuals). Distances are reduced.

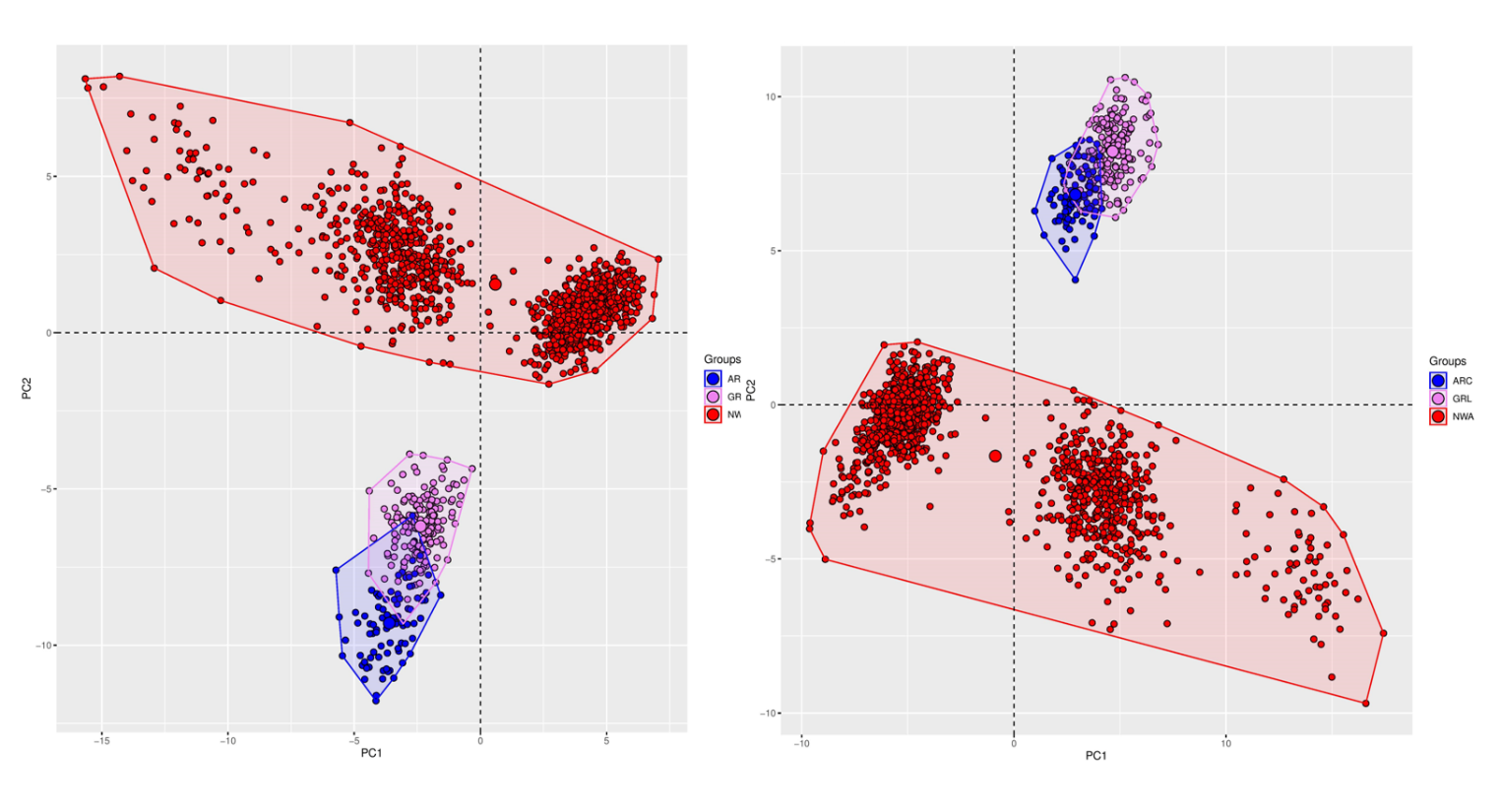

**Figure S11.** PCA performed by Chr2 (on the left) and Chr9 (on the right) showing that the same haplogroup pattern.

**Table S1**. Correspondence between the longest scaffolds in the four chromosome-level genome assembly of related species used to order *Mallotus villosus*’ contigs: *Esox lucius* (Rondeau et al. 2014), *Dicentrarchus labrax* (Tine et al. 2014), *Sparus aurata* (Pauletto et al. 2018), and *Takifugu rubripes* (Kai et al. 2011). Correspondence is based on observed synteny when first aligned to each other using D-Genies (Cabanettes & Klopp 2018). Cells in grey indicated chromosomal fusion relatively to *Esox lucius*.

| Arbitrary_name | *Esox lucius* (E) | *Dicentrarchus labrax* (D) | *Sparus aurata* (S) | *Takifugu rubripes* (T) |
| --- | --- | --- | --- | --- |
| Chr_1 | NC_025974.3 | HG916831.1 | CM010265.1 | NC_018904.1 |
| Chr_2 | NC_025978.3 | HG916848.1 | CM010275.1 | NC_018894.1 |
| Chr_3 | NC_025987.3 | HG916833.1 | CM010269.1 | NC_018896.1 |
| Chr_4 | NC_025968.3 | HG916830.1 | CM010254.1 | NC_018900.1 |
| Chr_5 | NC_025986.3 | HG916846.1 | CM010260.1 | NC_018898.1 |
| Chr_6 | NC_025984.3 | HG916837.1 | CM010258.1 | NC_018908.1 |
| Chr_7 | NC_025969.3 | HG916845.1 | CM010256.1 | NC_018902.1 |
| Chr_8 | NC_025980.3 | HG916840.1 | CM010257.1 | NC_018910.1 |
| Chr_9 | NC_025979.3 | HG916841.1 | CM010259.1 | NC_018892.1 |
| Chr_10 | NC_025981.3 | HG916836.1 | CM010264.1 | NC_018895.1 |
| Chr_11 | NC_025972.3 | HG916838.1 | CM010272.1 | NC_018890.1 |
| Chr_12 | NC_025983.3 | HG916832.1 | CM010261.1 | NC_018890.1 |
| Chr_13 | NC_025975.3 | HG916844.1 | CM010263.1 | NC_018909.1 |
| Chr_14 | NC_025982.3 | HG916829.1 | CM010268.1 | NC_018891.1 |
| Chr_15 | NC_025970.3 | HG916827.1 | CM010273.1 | NC_018911.1 |
| Chr_16 | NC_025973.3 | HG916828.1 | CM010267.1 | NC_018893.1 |
| Chr_17 | NC_025971.3 | HG916839.1 | CM010270.1 | NC_018903.1 |
| Chr_18 | NC_025977.3 | HG916849.1 | CM010255.1 | NC_018901.1 |
| Chr_19 | NC_025985.3 | HG916834.1 | CM010274.1 | NC_018905.1 |
| Chr_20 | NC_025988.3 | HG916835.1 | CM010271.1 | NC_018899.1 |
| Chr_21 | NC_025990.3 | HG916850.1 | CM010266.1 | NC_018907.1 |
| Chr_22 | NC_025991.3 | HG916843.1 | CM010262.1 | NC_018897.1 |
| Chr_23 | NC_025989.3 | HG916842.1 | CM010276.1 | NC_018897.1 |
| Chr_24 | NC_025976.3 | HG916847.1 | CM010253.1 | NC_018906.1 |
| Chr_U | NC_025992.3 | HG916847.1 | CM010253.1 | NC_018906.1 |

**Table S2**. Effect of haplogroups on SNPs heterozygosity. Estimates of the beta regression model.

| Parameter | Estimate | Std error | z-value | P-value |
| --- | --- | --- | --- | --- |
| Intercept | -2.38677 | 0.03296 | -72.417 | < 2e-16 |
| Haplogroup 1 | 0.20586 | 0.03808 | 5.406 | 6.46e-08 |
| Haplogroup 2 | 0.15851 | 0.03809 | 4.161 | 3.17e-05 |
| Phi coeffcient | 3.58938 | 0.09428 | 38.07 | <2e-16 |

**Table S3**. AIC, deltaAIC and AICw model choice for δaδi. The best models for each pair of populations are highlighted in bold italic. pop1 and pop2 shows the name of the population pairs compared (GRE = Groenland, ARC = Arctic, NWA =North West Atlantic). The ‘full” dataset included all the SNPs. The dataset “genome-wide” included all the SNPs but the NWA lineage was split into three groups according to their coordinates along PC1 axes. The data set “no-chr2 chr9” excluded all the SNPs falling on the chromosome 2 and chromosome 9 and the NWA lineage was split by groups. Models abbreviations: SC = Secondary Contact, IM = Isolation w. Migration, AM = Ancient Migration, SI = Strict Isolation. Suffix 2N = heterogeneous population size. 2M =heterogeneous migration rate. All alternative version of the SC, IM, AM and SI including or not heterogeneity in migration rate and effective population size were tested.

| **pop1** | **pop2** | **dataset** | **model** | **AIC** | **∆**  **AIC** | **AIC**  **weight** |  | **pop1** | **pop2** | **dataset** | **model** | **AIC** | **∆**  **AIC** | **AIC**  **weight** |
| --- | --- | --- | --- | --- | --- | --- | --- | --- | --- | --- | --- | --- | --- | --- |
| ***GRL*** | ***ARC*** | ***full*** | ***SC2N2m*** | ***4583*** | ***0*** | ***1,0*** |  | ***GRL*** | ***haplo3*** | ***genome wide*** | ***SC2N2m*** | ***6086*** | ***0*** | ***1,0*** |
| GRL | ARC | full | IM2N2m | 4614 | 31 | 0,0 |  | GRL | haplo3 | genome wide | SC2N | 6322 | 236 | 0,0 |
| GRL | ARC | full | SC2m | 4648 | 64 | 0,0 |  | GRL | haplo3 | genome wide | SC2m | 6421 | 334 | 0,0 |
| GRL | ARC | full | SC2N | 4661 | 77 | 0,0 |  | GRL | haplo3 | genome wide | AM2N | 6432 | 345 | 0,0 |
| GRL | ARC | full | AM2N | 4661 | 78 | 0,0 |  | GRL | haplo3 | genome wide | IM2N | 6442 | 356 | 0,0 |
| GRL | ARC | full | IM2N | 4669 | 86 | 0,0 |  | GRL | haplo3 | genome wide | IM2N2m | 6448 | 362 | 0,0 |
| GRL | ARC | full | IM2m | 4748 | 164 | 0,0 |  | GRL | haplo3 | genome wide | AM2N2m | 6636 | 550 | 0,0 |
| GRL | ARC | full | AM2m | 4749 | 166 | 0,0 |  | GRL | haplo3 | genome wide | IM2m | 6704 | 617 | 0,0 |
| GRL | ARC | full | AM2N2m | 4786 | 202 | 0,0 |  | GRL | haplo3 | genome wide | AM2m | 6705 | 618 | 0,0 |
| GRL | ARC | full | SC | 5478 | 894 | 0,0 |  | GRL | haplo3 | genome wide | SI2N | 9242 | 3156 | 0,0 |
| GRL | ARC | full | AM | 5520 | 936 | 0,0 |  | GRL | haplo3 | genome wide | SC | 9287 | 3201 | 0,0 |
| GRL | ARC | full | IM | 5521 | 937 | 0,0 |  | GRL | haplo3 | genome wide | IM | 9613 | 3526 | 0,0 |
| GRL | ARC | full | SI2N | 6130 | 1547 | 0,0 |  | GRL | haplo3 | genome wide | AM | 9615 | 3528 | 0,0 |
| GRL | ARC | full | SI | 7629 | 3045 | 0,0 |  | GRL | haplo3 | genome wide | SI | 13677 | 7591 | 0,0 |
| ***NORTH*** | ***GRL*** | ***full*** | ***SC2N2m*** | ***4934*** | ***0*** | ***1,0*** |  | ***ARC*** | ***haplo3*** | ***no chr2 chr9*** | ***IM2N2m*** | ***5596*** | ***0*** | ***1,0*** |
| NORTH | GRL | full | AM2N | 5174 | 240 | 0,0 |  | ARC | haplo3 | no chr2 chr9 | SC2N2m | 5696 | 100 | 0,0 |
| NORTH | GRL | full | SC2N | 5178 | 244 | 0,0 |  | ARC | haplo3 | no chr2 chr9 | AM2N | 5723 | 127 | 0,0 |
| NORTH | GRL | full | IM2N | 5195 | 261 | 0,0 |  | ARC | haplo3 | no chr2 chr9 | SC2N | 5764 | 168 | 0,0 |
| NORTH | GRL | full | SC2m | 5278 | 344 | 0,0 |  | ARC | haplo3 | no chr2 chr9 | IM2N | 5768 | 172 | 0,0 |
| NORTH | GRL | full | AM2N2m | 5357 | 424 | 0,0 |  | ARC | haplo3 | no chr2 chr9 | SC2m | 5880 | 284 | 0,0 |
| NORTH | GRL | full | IM2N2m | 5373 | 439 | 0,0 |  | ARC | haplo3 | no chr2 chr9 | AM2N2m | 5890 | 294 | 0,0 |
| NORTH | GRL | full | IM2m | 5467 | 533 | 0,0 |  | ARC | haplo3 | no chr2 chr9 | IM2m | 5977 | 381 | 0,0 |
| NORTH | GRL | full | AM2m | 5469 | 535 | 0,0 |  | ARC | haplo3 | no chr2 chr9 | AM2m | 5977 | 381 | 0,0 |
| NORTH | GRL | full | SC | 7033 | 2099 | 0,0 |  | ARC | haplo3 | no chr2 chr9 | SC | 6718 | 1122 | 0,0 |
| NORTH | GRL | full | AM | 7537 | 2603 | 0,0 |  | ARC | haplo3 | no chr2 chr9 | AM | 6740 | 1144 | 0,0 |
| NORTH | GRL | full | SI2N | 7657 | 2723 | 0,0 |  | ARC | haplo3 | no chr2 chr9 | IM | 6780 | 1184 | 0,0 |
| NORTH | GRL | full | IM | 7734 | 2800 | 0,0 |  | ARC | haplo3 | no chr2 chr9 | SI2N | 7941 | 2345 | 0,0 |
| NORTH | GRL | full | SI | 11513 | 6579 | 0,0 |  | ARC | haplo3 | no chr2 chr9 | SI | 9314 | 3718 | 0,0 |
| ***NORTH*** | ***ARC*** | ***full*** | ***IM2N2m*** | ***3998*** | ***0*** | ***1,0*** |  | ***ARC*** | ***haplo1*** | ***no chr2 chr9*** | ***IM2N2m*** | ***5611*** | ***0*** | ***1,0*** |
| NORTH | ARC | full | AM2N | 4017 | 20 | 0,0 |  | ARC | haplo1 | no chr2 chr9 | SC2N2m | 5678 | 67 | 0,0 |
| NORTH | ARC | full | IM2N | 4033 | 35 | 0,0 |  | ARC | haplo1 | no chr2 chr9 | AM2N | 5744 | 133 | 0,0 |
| NORTH | ARC | full | SC2N2m | 4036 | 38 | 0,0 |  | ARC | haplo1 | no chr2 chr9 | SC2N | 5779 | 168 | 0,0 |
| NORTH | ARC | full | SC2N | 4036 | 38 | 0,0 |  | ARC | haplo1 | no chr2 chr9 | IM2N | 5785 | 174 | 0,0 |
| NORTH | ARC | full | AM2N2m | 4158 | 160 | 0,0 |  | ARC | haplo1 | no chr2 chr9 | SC2m | 5911 | 300 | 0,0 |
| NORTH | ARC | full | SC2m | 4221 | 223 | 0,0 |  | ARC | haplo1 | no chr2 chr9 | AM2N2m | 5913 | 302 | 0,0 |
| NORTH | ARC | full | IM2m | 4341 | 343 | 0,0 |  | ARC | haplo1 | no chr2 chr9 | IM2m | 6007 | 396 | 0,0 |
| NORTH | ARC | full | AM2m | 4342 | 345 | 0,0 |  | ARC | haplo1 | no chr2 chr9 | AM2m | 6008 | 397 | 0,0 |
| NORTH | ARC | full | SC | 4956 | 959 | 0,0 |  | ARC | haplo1 | no chr2 chr9 | SC | 6756 | 1145 | 0,0 |
| NORTH | ARC | full | AM | 5009 | 1011 | 0,0 |  | ARC | haplo1 | no chr2 chr9 | AM | 6786 | 1175 | 0,0 |
| NORTH | ARC | full | IM | 5019 | 1022 | 0,0 |  | ARC | haplo1 | no chr2 chr9 | IM | 6824 | 1213 | 0,0 |
| NORTH | ARC | full | SI2N | 5610 | 1612 | 0,0 |  | ARC | haplo1 | no chr2 chr9 | SI2N | 8058 | 2447 | 0,0 |
| NORTH | ARC | full | SI | 7078 | 3080 | 0,0 |  | ARC | haplo1 | no chr2 chr9 | SI | 9473 | 3862 | 0,0 |
| ***ARC*** | ***haplo1*** | ***genome wide*** | ***IM2N2m*** | ***5842*** | ***0*** | ***1,0*** |  | ***GRL*** | ***haplo1*** | ***genome wide*** | ***SC2N2m*** | ***6131*** | ***0*** | ***1,0*** |
| ARC | haplo1 | genome wide | SC2N2m | 5942 | 100 | 0,0 |  | GRL | haplo1 | genome wide | SC2N | 6367 | 236 | 0,0 |
| ARC | haplo1 | genome wide | AM2N | 5976 | 134 | 0,0 |  | GRL | haplo1 | genome wide | SC2m | 6475 | 344 | 0,0 |
| ARC | haplo1 | genome wide | SC2N | 6018 | 176 | 0,0 |  | GRL | haplo1 | genome wide | AM2N | 6494 | 363 | 0,0 |
| ARC | haplo1 | genome wide | IM2N | 6020 | 179 | 0,0 |  | GRL | haplo1 | genome wide | IM2N | 6495 | 363 | 0,0 |
| ARC | haplo1 | genome wide | SC2m | 6161 | 319 | 0,0 |  | GRL | haplo1 | genome wide | IM2N2m | 6514 | 383 | 0,0 |
| ARC | haplo1 | genome wide | AM2N2m | 6177 | 335 | 0,0 |  | GRL | haplo1 | genome wide | AM2N2m | 6768 | 636 | 0,0 |
| ARC | haplo1 | genome wide | IM2m | 6283 | 441 | 0,0 |  | GRL | haplo1 | genome wide | IM2m | 6795 | 664 | 0,0 |
| ARC | haplo1 | genome wide | AM2m | 6284 | 442 | 0,0 |  | GRL | haplo1 | genome wide | AM2m | 6796 | 665 | 0,0 |
| ARC | haplo1 | genome wide | SC | 7147 | 1305 | 0,0 |  | GRL | haplo1 | genome wide | SC | 9423 | 3291 | 0,0 |
| ARC | haplo1 | genome wide | AM | 7182 | 1341 | 0,0 |  | GRL | haplo1 | genome wide | SI2N | 9537 | 3405 | 0,0 |
| ARC | haplo1 | genome wide | IM | 7223 | 1382 | 0,0 |  | GRL | haplo1 | genome wide | IM | 9775 | 3644 | 0,0 |
| ARC | haplo1 | genome wide | SI2N | 8579 | 2737 | 0,0 |  | GRL | haplo1 | genome wide | AM | 9777 | 3646 | 0,0 |
| ARC | haplo1 | genome wide | SI | 10191 | 4349 | 0,0 |  | GRL | haplo1 | genome wide | SI | 14125 | 7994 | 0,0 |
| ***ARC*** | ***haplo3*** | ***genome wide*** | ***IM2N2m*** | ***5822*** | ***0*** | ***1,0*** |  | ***GRL*** | ***haplo3*** | ***no chr2 chr9*** | ***SC2N2m*** | ***5892*** | ***0*** | ***1,0*** |
| ARC | haplo3 | genome wide | SC2N2m | 5913 | 91 | 0,0 |  | GRL | haplo3 | no chr2 chr9 | SC2N | 6102 | 210 | 0,0 |
| ARC | haplo3 | genome wide | AM2N | 5945 | 123 | 0,0 |  | GRL | haplo3 | no chr2 chr9 | IM2N | 6205 | 313 | 0,0 |
| ARC | haplo3 | genome wide | SC2N | 5995 | 172 | 0,0 |  | GRL | haplo3 | no chr2 chr9 | SC2m | 6208 | 315 | 0,0 |
| ARC | haplo3 | genome wide | IM2N | 6004 | 182 | 0,0 |  | GRL | haplo3 | no chr2 chr9 | AM2N | 6209 | 317 | 0,0 |
| ARC | haplo3 | genome wide | SC2m | 6118 | 296 | 0,0 |  | GRL | haplo3 | no chr2 chr9 | IM2N2m | 6256 | 364 | 0,0 |
| ARC | haplo3 | genome wide | AM2N2m | 6123 | 301 | 0,0 |  | GRL | haplo3 | no chr2 chr9 | AM2N2m | 6428 | 535 | 0,0 |
| ARC | haplo3 | genome wide | AM2m | 6225 | 403 | 0,0 |  | GRL | haplo3 | no chr2 chr9 | IM2m | 6469 | 577 | 0,0 |
| ARC | haplo3 | genome wide | IM2m | 6226 | 403 | 0,0 |  | GRL | haplo3 | no chr2 chr9 | AM2m | 6471 | 579 | 0,0 |
| ARC | haplo3 | genome wide | SC | 7080 | 1257 | 0,0 |  | GRL | haplo3 | no chr2 chr9 | SC | 8634 | 2742 | 0,0 |
| ARC | haplo3 | genome wide | AM | 7103 | 1281 | 0,0 |  | GRL | haplo3 | no chr2 chr9 | SI2N | 8844 | 2952 | 0,0 |
| ARC | haplo3 | genome wide | IM | 7145 | 1323 | 0,0 |  | GRL | haplo3 | no chr2 chr9 | IM | 8945 | 3052 | 0,0 |
| ARC | haplo3 | genome wide | SI2N | 8365 | 2543 | 0,0 |  | GRL | haplo3 | no chr2 chr9 | AM | 8947 | 3054 | 0,0 |
| ARC | haplo3 | genome wide | SI | 9914 | 4092 | 0,0 |  | GRL | haplo3 | no chr2 chr9 | SI | 12674 | 6782 | 0,0 |
| ***GRL*** | ***haplo1*** | ***no chr2 chr9*** | ***SC2N2m*** | ***5926*** | ***0*** | ***1,0*** |  | ***NWA*** | ***ARC*** | ***no chr2 chr9*** | IM2N2m | 3928 | 0. | 1 |
| GRL | haplo1 | no chr2 chr9 | SC2N | 6126 | 200 | 0,0 |  | *NWA* | *ARC* | *no chr2 chr9* | AM2N | 3983 | 55 | 0 |
| GRL | haplo1 | no chr2 chr9 | SC2m | 6235 | 309 | 0,0 |  | *NWA* | *ARC* | *no chr2 chr9* | IM2N | 4005 | 77 | 0 |
| GRL | haplo1 | no chr2 chr9 | AM2N | 6243 | 318 | 0,0 |  | *NWA* | *ARC* | *no chr2 chr9* | SC2N | 4006 | 78 | 0 |
| GRL | haplo1 | no chr2 chr9 | IM2N | 6253 | 327 | 0,0 |  | *NWA* | *ARC* | *no chr2 chr9* | SC2N2m | 4006 | 78 | 0 |
| GRL | haplo1 | no chr2 chr9 | IM2N2m | 6293 | 368 | 0,0 |  | *NWA* | *ARC* | *no chr2 chr9* | AM2N2m | 4110 | 182 | 0 |
| GRL | haplo1 | no chr2 chr9 | AM2N2m | 6452 | 527 | 0,0 |  | *NWA* | *ARC* | *no chr2 chr9* | SC2m | 4169 | 241 | 0 |
| GRL | haplo1 | no chr2 chr9 | IM2m | 6519 | 594 | 0,0 |  | *NWA* | *ARC* | *no chr2 chr9* | IM2m | 4247 | 319 | 0 |
| GRL | haplo1 | no chr2 chr9 | AM2m | 6522 | 596 | 0,0 |  | *NWA* | *ARC* | *no chr2 chr9* | AM2m | 4249 | 320 | 0 |
| GRL | haplo1 | no chr2 chr9 | SC | 8763 | 2837 | 0,0 |  | *NWA* | *ARC* | *no chr2 chr9* | SC | 4779 | 850 | 0 |
| GRL | haplo1 | no chr2 chr9 | SI2N | 9002 | 3076 | 0,0 |  | *NWA* | *ARC* | *no chr2 chr9* | AM | 4805 | 877 | 0 |
| GRL | haplo1 | no chr2 chr9 | IM | 9087 | 3161 | 0,0 |  | *NWA* | *ARC* | *no chr2 chr9* | IM | 4821 | 892 | 0 |
| GRL | haplo1 | no chr2 chr9 | AM | 9089 | 3163 | 0,0 |  | *NWA* | *ARC* | *no chr2 chr9* | SI2N | 5663 | 1735 | 0 |
| GRL | haplo1 | no chr2 chr9 | SI | 12986 | 7060 | 0,0 |  | *NWA* | *ARC* | *no chr2 chr9* | SI | 6674 | 2745 | 0 |
| ***NWA*** | ***GRE*** | ***no chr2 chr9*** | SC2N2m | 4733 | 0 | 1 |  |  |  |  |  |  |  |  |
| *NWA* | *GRE* | *no chr2 chr9* | IM2N2m | 4890 | 156 | 0 |  |  |  |  |  |  |  |  |
| *NWA* | *GRE* | *no chr2 chr9* | SC2N | 4946 | 212 | 0 |  |  |  |  |  |  |  |  |
| *NWA* | *GRE* | *no chr2 chr9* | AM2N | 4963 | 229 | 0 |  |  |  |  |  |  |  |  |
| *NWA* | *GRE* | *no chr2 chr9* | IM2N | 4973 | 239 | 0 |  |  |  |  |  |  |  |  |
| *NWA* | *GRE* | *no chr2 chr9* | SC2m | 5040 | 306 | 0 |  |  |  |  |  |  |  |  |
| *NWA* | *GRE* | *no chr2 chr9* | AM2N2m | 5133 | 399 | 0 |  |  |  |  |  |  |  |  |
| *NWA* | *GRE* | *no chr2 chr9* | IM2m | 5218 | 485 | 0 |  |  |  |  |  |  |  |  |
| *NWA* | *GRE* | *no chr2 chr9* | AM2m | 5220 | 487 | 0 |  |  |  |  |  |  |  |  |
| *NWA* | *GRE* | *no chr2 chr9* | SC | 7027 | 2294 | 0 |  |  |  |  |  |  |  |  |
| *NWA* | *GRE* | *no chr2 chr9* | IM | 7245 | 2511 | 0 |  |  |  |  |  |  |  |  |
| *NWA* | *GRE* | *no chr2 chr9* | AM | 7247 | 2513 | 0 |  |  |  |  |  |  |  |  |
| *NWA* | *GRE* | *no chr2 chr9* | SI2N | 7458 | 2724 | 0 |  |  |  |  |  |  |  |  |
| *NWA* | *GRE* | *no chr2 chr9* | SI | 10690 | 5956 | 0 |  |  |  |  |  |  |  |  |

**Table S4.** Detailed parameter estimates from ∂a∂i for a) all data, b) split by haplogroups and c) excluding the chr2 and chr9. SD provides the standard errors around the parameter estimates.

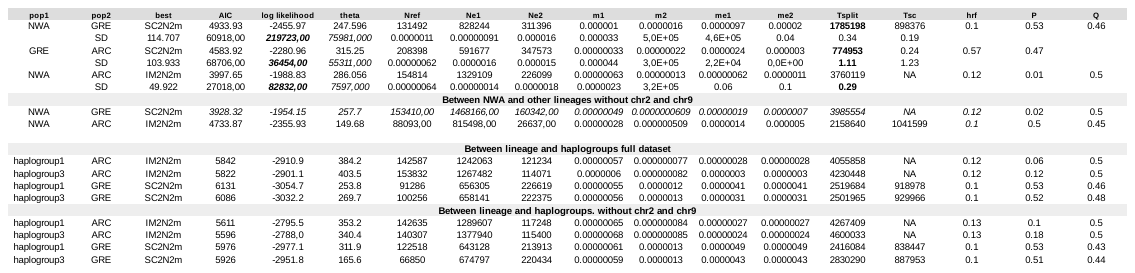

**Table S5: AIC, ∆AIC and AIC weights for model selection among the two haplogroups.** The SC2N2m model received the highest support based on all metrics, followed by other alternative version of the SC. The SC2N2m model assumes heterogeneous gene flow (suffix 2m) and heterogeneous effective population size (suffix 2N) Other models were not supported.

| **haplogroup1** | **haplogroup2** | **model** | **AIC** | **deltaAIC** | **AICweights** |
| --- | --- | --- | --- | --- | --- |
| ***haplogroup1*** | ***haplogroup3*** | ***SC2N2m*** | ***4773.4*** | ***0*** | ***1*** |
| haplogroup1 | haplogroup3 | SC2m | 4818.8 | 45.5 | 0 |
| haplogroup1 | haplogroup3 | SC2N | 5049.6 | 276.3 | 0 |
| haplogroup1 | haplogroup3 | SC | 5116.3 | 343.0 | 0 |
| haplogroup1 | haplogroup3 | IM2N2m | 6051.9 | 1278.5 | 0 |
| haplogroup1 | haplogroup3 | SI2N | 6669.5 | 1896.1 | 0 |
| haplogroup1 | haplogroup3 | AM2N2m | 6870.6 | 2097.3 | 0 |
| haplogroup1 | haplogroup3 | SI | 7151.1 | 2377.7 | 0 |
| haplogroup1 | haplogroup3 | IM | 7155.4 | 2382.1 | 0 |
| haplogroup1 | haplogroup3 | AM | 7157.7 | 2384.4 | 0 |
| haplogroup1 | haplogroup3 | IM2N | 7160.0 | 2386.6 | 0 |
| haplogroup1 | haplogroup3 | AM2N | 7162.4 | 2389.0 | 0 |
| haplogroup1 | haplogroup3 | IM2m | 7162.7 | 2389.4 | 0 |
| haplogroup1 | haplogroup3 | AM2m | 7164.1 | 2390.8 | 0 |

**Table S6:** Estimates of demographic parameters under the best models (SC2N2m) between each haplogroup1 and 3 corresponding defined based on the PCA (see methods). Legend is the same as in Table 1 in main text.

|  | **haplogroup 1 vs haplogroup 3** |
| --- | --- |
| best | SC2N2m |
| Nref | 43700 |
| Ne1 | 527700 [45700–598000] |
| Ne3 | 189000 [163000–214000] |
| m1 | 0.00032 [0.00025 – 0.00039] |
| M3 | 0.00083 [0.0005 – 0.001] |
| me1 | 0.0000013 [0 – 0.000002] |
| Me3 | 0.00000097 [0 – 0.000003] |
| Tsplit | 3,115,000 [2,710,000 – 3,517, 000] |
| Tsc | 35,100 [31,200 – 39,000] |
| hrf | 0.213 [0.13 – 0.295] |
| P | 0.95 [0.80 – 1] |
| Q | 0.123 [0.06–0.18] |

**Table S7: Distribution of candidate outliers across chromosome inferred based on explicit demo-scan.** Shown is the distribution of outliers identified between the three lineages, the column “significant hit” displays the distribution of mutation with significant e-value (< 1e^-10^).

| **CHR** | **NWA vs GRE** | **NWA vs ARC** | **ARC vs GRE** | **Significant hits** | **nonsynonymous mutation** |
| --- | --- | --- | --- | --- | --- |
| Chr_1 | 12 | 32 | 115 | 36 | 1 |
| Chr_10 | 13 | 20 | 86 | 29 | 2 |
| Chr_11 | 7 | 11 | 53 | 18 | 1 |
| Chr_12 | 25 | 35 | 100 | 43 | 2 |
| Chr_13 | 15 | 39 | 99 | 36 | 4 |
| Chr_14 | 7 | 18 | 90 | 23 | 1 |
| Chr_15 | 16 | 26 | 85 | 29 | 1 |
| Chr_16 | 7 | 13 | 63 | 15 | 2 |
| Chr_17 | 10 | 28 | 69 | 21 | 1 |
| Chr_18 | 12 | 22 | 69 | 20 | 3 |
| Chr_19 | 16 | 28 | 111 | 29 | 0 |
| Chr_2 | 13 | 13 | 71 | 24 | 1 |
| Chr_20 | 12 | 31 | 77 | 19 | 2 |
| Chr_21 | 5 | 23 | 72 | 25 | 4 |
| Chr_22 | 6 | 14 | 51 | 6 | 0 |
| Chr_23 | 11 | 18 | 58 | 31 | 1 |
| Chr_24 | 14 | 19 | 77 | 20 | 1 |
| Chr_3 | 16 | 38 | 96 | 39 | 0 |
| Chr_4 | 17 | 29 | 88 | 44 | 4 |
| Chr_5 | 13 | 25 | 101 | 38 | 1 |
| Chr_6 | 17 | 43 | 80 | 27 | 0 |
| Chr_7 | 25 | 27 | 86 | 29 | 1 |
| Chr_8 | 13 | 28 | 118 | 39 | 4 |
| Chr_9 | 20 | 31 | 94 | 29 | 2 |
| Chr_unplaced | 8 | 11 | 76 | 18 | 1 |

**Table S8: Blast results for the best outliers associated with reproductive isolation between lineages.**

| **Contig** | **POS** | **SNP_ID** | **ProteinID** | **evalue** |
| --- | --- | --- | --- | --- |
| contig_1337 | 54196 | 5807_68 | sp\|Q9NR99\|MXRA5_HUMAN | 2.18e-103 |
| contig_1560 | 304560 | 10059_49 | sp\|Q8BH53\|CFA69_MOUSE | 1.58e-48 |
| contig_104 | 330576 | 1627_28 | sp\|Q8BVI5\|STX16_MOUSE | 9.43e-22 |
| contig_275 | 75424 | 21582_46 | sp\|Q9BXT8\|RNF17_HUMAN | 7.68e-20 |
| contig_223 | 72354 | 15829_11 | sp\|Q91684\|DPOG1_XENLA | 3.30e-58 |
| contig_232 | 1009303 | 16707_60 | sp\|Q9Y5L0\|TNPO3_HUMAN | 9.84e-69 |
| contig_546 | 176988 | 47863_65 | sp\|A3KP85\|ETKMT_DANRE | 5.81e-39 |
| contig_263 | 34011 | 19853_10 | sp\|Q9C0D6\|FHDC1_HUMAN | 4.03e-14 |
| contig_2971 | 75583 | 23848_62 | sp\|P10039\|TENA_CHICK | 3.45e-138 |
| contig_358 | 319676 | 31922_56 | sp\|P19835\|CEL_HUMAN | 1.42e-11 |
| contig_1378 | 129139 | 6654_29 | sp\|P79782\|TCF15_CHICK | 4.67e-40 |
| contig_489 | 158508 | 45385_51 | sp\|P06280\|AGAL_HUMAN | 5.33e-29 |
| scaffold_1851 | 78009 | 62126_42 | sp\|Q96NI6\|LRFN5_HUMAN | 3.62e-113 |
| contig_757 | 600753 | 55181_66 | sp\|O43847\|NRDC_HUMAN | 7.34e-19 |
| contig_989 | 862124 | 61334_59 | sp\|Q91W43\|GCSP_MOUSE | 7.28e-31 |
| scaffold_1118 | 456224 | 61728_7 | sp\|Q1LWH4\|FAN1_DANRE | 1.06e-17 |
| contig_15 | 106288 | 10281_32 | sp\|Q9TU53\|CUBN_CANLF | 1.38e-19 |
| contig_4380 | 15833 | 43058_37 | sp\|F1LMY4\|RYR1_RAT | 8.92e-81 |
| contig_2940 | 79330 | 23459_7 | sp\|A3KNA5\|FIL1L_DANRE | 0.0 |
| contig_115 | 406078 | 3748_42 | sp\|P34057\|RECO_MOUSE | 1.86e-52 |
| contig_1025 | 12303 | 771_58 | sp\|O18738\|DAG1_BOVIN | 1.41e-111 |
| contig_1083 | 97816 | 2560_40 | sp\|Q8IVF5\|TIAM2_HUMAN | 1.13e-20 |
| scaffold_5048 | 18400 | 63104_78 | sp\|Q9ESM2\|HPLN2_RAT | 6.96e-43 |
| contig_249 | 134450 | 18439_73 | sp\|Q7LHG5\|YI31B_YEAST | 2.39e-50 |
| scaffold_1118 | 476551 | 61732_49 | sp\|Q9NRD8\|DUOX2_HUMAN | 1.02e-92 |
| contig_370 | 328434 | 33652_74 | sp\|E7ERA6\|RN223_HUMAN | 6.67e-23 |
| contig_2171 | 80969 | 15032_64 | sp\|Q45R42\|LRRC4_RAT | 0.0 |
| scaffold_1118 | 82746 | 61790_60 | sp\|P14650\|PERT_RAT | 3.95e-64 |
| contig_29 | 144151 | 24270_40 | sp\|Q04637\|IF4G1_HUMAN | 8.56e-49 |
| contig_1665 | 686330 | 10696_53 | sp\|P30372\|ACM2_CHICK | 0.0 |
| contig_515 | 63874 | 46511_7 | sp\|Q92817\|EVPL_HUMAN | 0.0 |
| contig_3177 | 19748 | 25977_58 | sp\|P00535\|ERBB_AVIER | 3.60e-44 |
| contig_1120 | 446168 | 3056_37 | sp\|Q96DT5\|DYH11_HUMAN | 1.09e-48 |
| contig_4758 | 109739 | 44094_28 | sp\|O94979\|SC31A_HUMAN | 1.07e-39 |
| contig_644 | 123922 | 51468_25 | sp\|F4K487\|PHOX3_ARATH | 6.16e-14 |
| contig_2870 | 49708 | 22548_7 | sp\|Q9NTX9\|F217B_HUMAN | 2.24e-14 |
| contig_373 | 491926 | 34121_19 | sp\|P26010\|ITB7_HUMAN | 1.68e-34 |
| contig_5646 | 316463 | 48496_37 | sp\|P98088\|MUC5A_HUMAN | 1.23e-26 |
| contig_1325 | 120786 | 5567_73 | sp\|P35408\|PE2R4_HUMAN | 7.88e-122 |
| contig_1088 | 27511 | 2666_61 | sp\|Q5NVC7\|RNF34_PONAB | 1.15e-32 |
| contig_2949 | 16719 | 23515_30 | sp\|Q8BQN5\|FA78B_MOUSE | 9.11e-34 |
| contig_1811 | 7511 | 11803_64 | sp\|Q69Z23\|DYH17_MOUSE | 7.84e-34 |
| contig_1374 | 40864 | 6588_8 | sp\|E9PZQ0\|RYR1_MOUSE | 3.86e-115 |
| contig_3544 | 420130 | 31302_59 | sp\|Q6IZ48\|SOX8_TETNG | 2.59e-79 |
| contig_115 | 718117 | 3809_52 | sp\|Q8N884\|CGAS_HUMAN | 1.98e-27 |
| contig_853 | 40921 | 57999_42 | sp\|Q9UKA4\|AKA11_HUMAN | 2.56e-71 |
| contig_585 | 24387 | 49521_19 | sp\|Q9Y239\|NOD1_HUMAN | 1.23e-138 |
| contig_319 | 1486567 | 26206_36 | sp\|Q5PRC0\|CH25H_DANRE | 1.04e-108 |
| contig_3979 | 35194 | 37882_14 | sp\|F6NSX9\|Z687B_DANRE | 1.92e-72 |
| contig_479 | 970098 | 45119_42 | sp\|Q8NFP9\|NBEA_HUMAN | 1.08e-28 |
| contig_2306 | 70568 | 16452_72 | sp\|Q64332\|SYN2_MOUSE | 1.04e-22 |
| contig_7105 | 74901 | 53500_62 | sp\|P14629\|ERCC5_XENLA | 1.78e-44 |
| contig_278 | 217936 | 21799_31 | sp\|Q5XI42\|AL3B1_RAT | 5.34e-41 |
| contig_2948 | 89782 | 23506_65 | sp\|P07099\|HYEP_HUMAN | 4.94e-32 |
| contig_479 | 2942157 | 44927_65 | sp\|E7FEV0\|WDR81_DANRE | 0.0 |
| contig_599 | 57253 | 49963_15 | sp\|Q9NBX4\|RTXE_DROME | 2.26e-16 |
| contig_501 | 4319 | 46014_5 | sp\|Q9Y2T6\|GPR55_HUMAN | 6.04e-40 |
| contig_839 | 31441 | 57784_9 | sp\|Q5XIN7\|PMGT1_RAT | 3.62e-17 |
| contig_279 | 46251 | 21900_54 | sp\|Q9QZ81\|AGO2_RAT | 1.23e-28 |
| contig_4210 | 73046 | 41280_38 | sp\|Q8CJ96\|RASF8_MOUSE | 1.38e-32 |
| contig_856 | 339640 | 58042_16 | sp\|Q92604\|LGAT1_HUMAN | 4.46e-31 |
| contig_7780 | 28901 | 55866_36 | sp\|Q95SX7\|RTBS_DROME | 2.98e-22 |
| contig_1990 | 7292 | 13650_59 | sp\|Q54KA7\|SECG_DICDI | 8.79e-25 |
| contig_5238 | 21433 | 46776_50 | sp\|Q7LHG5\|YI31B_YEAST | 8.20e-80 |
| contig_588 | 590561 | 49728_5 | sp\|Q02566\|MYH6_MOUSE | 4.90e-157 |
| contig_988 | 361304 | 61136_12 | sp\|Q5ZHT1\|ACD11_CHICK | 4.00e-29 |
| contig_1384 | 27025 | 6723_55 | sp\|O35709\|ENC1_MOUSE | 0.0 |
| contig_263 | 1051062 | 19712_53 | sp\|Q6NXK2\|ZN532_MOUSE | 2.34e-110 |
| contig_2348 | 4323 | 17013_54 | sp\|Q13615\|MTMR3_HUMAN | 6.34e-40 |
| contig_466 | 556307 | 43676_52 | sp\|Q91790\|DUS1A_XENLA | 1.30e-49 |
| contig_901 | 286377 | 59396_58 | sp\|Q6IQ20\|NAPEP_HUMAN | 1.10e-136 |
| contig_2028 | 160817 | 13944_69 | sp\|P62944\|AP2B1_RAT | 2.25e-34 |
| contig_1838 | 64244 | 11941_72 | sp\|P42331\|RHG25_HUMAN | 5.55e-11 |
| contig_3937 | 103530 | 37054_30 | sp\|A0FGR8\|ESYT2_HUMAN | 1.60e-21 |
| contig_3552 | 16564 | 31375_44 | sp\|Q8BNA6\|FAT3_MOUSE | 6.47e-41 |
| contig_1337 | 222934 | 5780_49 | sp\|Q62722\|NAB1_RAT | 1.00e-115 |
| contig_424 | 410403 | 42017_58 | sp\|Q6NUP7\|PP4R4_HUMAN | 6.89e-14 |
| contig_2271 | 140312 | 16083_19 | sp\|Q96A49\|SYAP1_HUMAN | 4.47e-21 |
| contig_716 | 36636 | 53880_58 | sp\|P01267\|THYG_BOVIN | 1.55e-23 |
| contig_1223 | 206854 | 4835_53 | sp\|O96028\|NSD2_HUMAN | 1.51e-59 |
| contig_424 | 375263 | 42010_22 | sp\|Q96Q27\|ASB2_HUMAN | 5.16e-27 |
| contig_1321 | 38470 | 5446_69 | sp\|Q4LFA9\|SEM3G_MOUSE | 2.39e-26 |
| contig_319 | 1211550 | 26158_38 | sp\|E9Q7T7\|CHADL_MOUSE | 8.41e-35 |
| contig_758 | 188792 | 55237_29 | sp\|Q08BB2\|P2012_DANRE | 1.11e-64 |
| contig_4121 | 13322 | 39909_13 | sp\|Q6DIL6\|F124A_XENTR | 1.38e-84 |
| contig_3265 | 27338 | 27447_33 | sp\|P53804\|TTC3_HUMAN | 9.95e-22 |
| contig_877 | 563954 | 58910_54 | sp\|P35249\|RFC4_HUMAN | 5.10e-37 |
| contig_2971 | 75025 | 23847_61 | sp\|P10039\|TENA_CHICK | 3.45e-138 |
| contig_3480 | 82794 | 30449_41 | sp\|Q8N8N0\|RN152_HUMAN | 2.70e-95 |
| contig_4046 | 329951 | 38616_79 | sp\|A2A884\|ZEP3_MOUSE | 1.80e-28 |
| contig_859 | 136679 | 58167_35 | sp\|Q5DTT1\|NEXMI_MOUSE | 0.0 |
| scaffold_1118 | 71680 | 61789_58 | sp\|Q8NFU1\|BEST2_HUMAN | 3.62e-33 |
| contig_2210 | 13157 | 15529_53 | sp\|Q92832\|NELL1_HUMAN | 1.55e-20 |
| contig_586 | 219636 | 49586_72 | sp\|Q9NYY8\|FAKD2_HUMAN | 8.09e-20 |
| contig_1418 | 30708 | 7719_52 | sp\|Q5REV5\|ACSM3_PONAB | 2.30e-35 |
| contig_2686 | 69765 | 20382_44 | sp\|Q07008\|NOTC1_RAT | 1.43e-57 |
| contig_1027 | 748809 | 934_14 | sp\|A0JMQ0\|BOP1_DANRE | 1.78e-25 |
| contig_2932 | 95298 | 23310_51 | sp\|Q3TCH7\|CUL4A_MOUSE | 7.99e-54 |
| contig_158 | 113782 | 10181_11 | sp\|F1R345\|DDX11_DANRE | 2.81e-51 |
| contig_406 | 268452 | 39022_54 | sp\|Q9UNA0\|ATS5_HUMAN | 1.24e-62 |
| contig_1692 | 144122 | 10868_23 | sp\|P56564\|EAA1_MOUSE | 1.04e-61 |
| contig_930 | 12567 | 60058_41 | sp\|P35072\|TCB1_CAEBR | 1.24e-59 |
| scaffold_3795 | 757247 | 62780_59 | sp\|Q6ZVH7\|ESPNL_HUMAN | 1.62e-49 |
| contig_2736 | 40007 | 21221_12 | sp\|Q8TEU8\|WFKN2_HUMAN | 0.0 |
| contig_369 | 52818 | 33477_37 | sp\|Q9JKY5\|HIP1R_MOUSE | 2.94e-28 |
| contig_7152 | 224213 | 53747_25 | sp\|Q8BG87\|TET3_MOUSE | 4.55e-20 |
| contig_1413 | 136984 | 7543_17 | sp\|Q2KJC1\|CDC5L_BOVIN | 8.85e-55 |
| contig_186 | 701302 | 12411_71 | sp\|Q7Z614\|SNX20_HUMAN | 2.62e-54 |
| contig_3452 | 236234 | 29916_16 | sp\|P08938\|PURP_CHICK | 9.23e-19 |
| contig_489 | 124309 | 45375_39 | sp\|Q2QCI8\|MED12_DANRE | 1.28e-53 |
| contig_1011 | 1026775 | 272_34 | sp\|B0R0I6\|CHD8_DANRE | 2.80e-52 |
| contig_1337 | 239740 | 5784_68 | sp\|Q9BZR6\|RTN4R_HUMAN | 1.08e-19 |
| contig_3267 | 112551 | 27457_44 | sp\|Q8NEA6\|GLIS3_HUMAN | 7.60e-68 |
| contig_4889 | 9840 | 45333_61 | sp\|Q9H0H3\|KLH25_HUMAN | 0.0 |
| contig_943 | 171166 | 60282_56 | sp\|Q63744\|RHG07_RAT | 9.55e-35 |
| contig_553 | 242233 | 48132_21 | sp\|Q5TYW1\|ZN658_HUMAN | 6.94e-117 |
| contig_3621 | 223191 | 32428_46 | sp\|Q13023\|AKAP6_HUMAN | 3.67e-35 |
| contig_2028 | 26041 | 13972_55 | sp\|O18973\|RABX5_BOVIN | 7.81e-70 |
| contig_1917 | 234754 | 12798_60 | sp\|P35072\|TCB1_CAEBR | 1.14e-51 |
| contig_367 | 18799 | 33366_43 | sp\|P10394\|POL4_DROME | 2.71e-21 |
| contig_929 | 74927 | 60040_17 | sp\|Q8VE62\|PAIP1_MOUSE | 4.67e-26 |
| contig_904 | 73575 | 59448_73 | sp\|Q0II29\|CE024_BOVIN | 6.65e-43 |
| contig_3564 | 235507 | 31478_34 | sp\|P20237\|GBRA4_BOVIN | 1.40e-30 |
| contig_828 | 326823 | 57411_49 | sp\|Q5RGQ8\|T200A_DANRE | 1.91e-131 |
| contig_275 | 362417 | 21547_77 | sp\|Q2EMV9\|PAR14_MOUSE | 8.37e-19 |
| contig_3994 | 622409 | 38123_35 | sp\|Q80XG9\|LRRT4_MOUSE | 4.43e-166 |
| contig_617 | 1092671 | 50525_37 | sp\|O57415\|RREB1_CHICK | 2.44e-75 |
| contig_1876 | 46426 | 12527_33 | sp\|Q501V0\|KPSH1_DANRE | 8.87e-80 |
| contig_2474 | 73035 | 18278_11 | sp\|P22725\|WNT5A_MOUSE | 1.63e-80 |
| contig_1383 | 165450 | 6701_79 | sp\|Q9C0G6\|DYH6_HUMAN | 9.95e-18 |
| scaffold_1118 | 406652 | 61717_50 | sp\|Q2KHR2\|RFX7_HUMAN | 2.33e-133 |
| contig_1662 | 62749 | 10585_32 | sp\|A1XQX0\|NR1AA_DANRE | 4.05e-74 |
| contig_481 | 27806 | 45185_68 | sp\|Q9BX84\|TRPM6_HUMAN | 7.38e-40 |
| contig_296 | 222829 | 23785_73 | sp\|P21783\|NOTC1_XENLA | 1.86e-97 |
| contig_1011 | 213414 | 419_29 | sp\|Q9UQB3\|CTND2_HUMAN | 6.98e-34 |
| contig_2842 | 3759 | 22314_72 | sp\|P20469\|ICEA_PANAN | 3.48e-20 |
| scaffold_992 | 248978 | 63338_46 | sp\|Q5RHB5\|LRMP_DANRE | 9.05e-35 |
| contig_5140 | 5935 | 46485_65 | sp\|P53353\|ASPX_VULVU | 5.50e-12 |
| contig_1320 | 36605 | 5421_31 | sp\|E2RYF7\|PBMU2_HUMAN | 8.89e-11 |
| contig_4189 | 91700 | 40795_76 | sp\|Q6PAC3\|DCA13_MOUSE | 3.71e-38 |
| contig_4058 | 178017 | 38772_26 | sp\|Q9Z1S0\|BUB1B_MOUSE | 1.69e-30 |
| contig_6823 | 129567 | 52570_45 | sp\|Q9CUL5\|DRC11_MOUSE | 1.25e-34 |
| contig_710 | 33389 | 53555_11 | sp\|Q6XK22\|R9BP_CHICK | 7.95e-85 |
| contig_1200 | 891597 | 4593_16 | sp\|Q6PFW1\|VIP1_HUMAN | 1.23e-38 |
| scaffold_122 | 1200901 | 61885_58 | sp\|Q6DDJ3\|DFI8A_XENLA | 2.57e-35 |
| contig_1480 | 142168 | 9032_77 | sp\|Q02253\|MMSA_RAT | 6.14e-58 |
| contig_3595 | 105536 | 31994_44 | sp\|Q95SX7\|RTBS_DROME | 4.63e-11 |
| contig_813 | 684816 | 56919_7 | sp\|Q9WU22\|PTN4_MOUSE | 9.97e-32 |
| contig_479 | 1941594 | 44746_50 | sp\|Q58EG3\|NECT3_DANRE | 5.86e-45 |
| contig_1146 | 446532 | 3609_43 | sp\|A2ARV4\|LRP2_MOUSE | 1.30e-116 |
| contig_3445 | 44027 | 29816_58 | sp\|Q6XPR3\|RPTN_HUMAN | 8.99e-14 |
| contig_3172 | 50323 | 25872_21 | sp\|Q99NG0\|ARIP4_MOUSE | 3.21e-26 |
| contig_5557 | 90378 | 48232_68 | sp\|Q5D1E7\|ZC12A_MOUSE | 5.49e-51 |
| contig_321 | 176057 | 26995_33 | sp\|O95983\|MBD3_HUMAN | 2.57e-64 |
| contig_7404 | 105756 | 54522_14 | sp\|Q5TJ59\|TLR3_BOVIN | 9.78e-37 |
| contig_3124 | 346308 | 25496_15 | sp\|Q14789\|GOGB1_HUMAN | 9.40e-12 |
| contig_921 | 129882 | 59832_24 | sp\|Q08BN9\|NXPE3_DANRE | 2.44e-95 |
| contig_5646 | 130870 | 48459_9 | sp\|Q6IQ23\|PKHA7_HUMAN | 1.33e-19 |
| contig_1850 | 30550 | 12065_74 | sp\|P46063\|RECQ1_HUMAN | 4.93e-51 |
| contig_3818 | 40139 | 35378_8 | sp\|Q8BH70\|FBXL4_MOUSE | 9.27e-37 |
| contig_419 | 202308 | 40906_77 | sp\|P29294\|MYLK_RABIT | 6.25e-22 |
| contig_4181 | 17510 | 40672_14 | sp\|Q9UQF2\|JIP1_HUMAN | 2.22e-29 |
| contig_5646 | 367999 | 48505_48 | sp\|P17427\|AP2A2_MOUSE | 5.96e-82 |
| contig_1568 | 88296 | 10114_15 | sp\|Q8WUH2\|TGFA1_HUMAN | 1.82e-37 |
| contig_818 | 202581 | 57041_55 | sp\|Q92990\|GLMN_HUMAN | 1.90e-11 |
| contig_1577 | 26619 | 10154_16 | sp\|Q14703\|MBTP1_HUMAN | 2.14e-37 |
| contig_2363 | 73045 | 17164_34 | sp\|Q9BXA9\|SALL3_HUMAN | 5.26e-76 |
| contig_2806 | 119456 | 21966_32 | sp\|Q96DT5\|DYH11_HUMAN | 2.16e-84 |
| contig_720 | 300147 | 54014_55 | sp\|Q9BG79\|SOST_BOVIN | 6.40e-40 |
| contig_40 | 116229 | 39450_61 | sp\|Q96RT8\|GCP5_HUMAN | 7.63e-93 |
| contig_2696 | 167847 | 20641_67 | sp\|P33121\|ACSL1_HUMAN | 1.57e-42 |
| contig_417 | 110975 | 40636_64 | sp\|Q32KQ2\|WDR53_BOVIN | 2.34e-29 |
| contig_4190 | 38884 | 40809_71 | sp\|Q9UQ26\|RIMS2_HUMAN | 3.85e-49 |
| contig_2422 | 44636 | 17859_66 | sp\|Q8IUR5\|TMTC1_HUMAN | 9.17e-20 |
| contig_818 | 291258 | 57058_64 | sp\|Q91Y81\|SEPT2_RAT | 4.08e-57 |
| contig_364 | 477713 | 33008_76 | sp\|A2RUW7\|PCX4_DANRE | 4.36e-29 |
| contig_15 | 92346 | 10305_22 | sp\|O60494\|CUBN_HUMAN | 2.72e-19 |
| contig_2545 | 135364 | 18794_49 | sp\|E9Q8T7\|DYH1_MOUSE | 3.97e-128 |
| contig_3955 | 145335 | 37558_38 | sp\|E9Q7X7\|NRX2A_MOUSE | 3.16e-51 |
| contig_771 | 94438 | 55748_56 | sp\|Q7LHG5\|YI31B_YEAST | 7.56e-74 |
| contig_7550 | 74 | 55058_8 | sp\|Q92105\|AT2A1_PELES | 2.41e-53 |
| contig_59 | 66577 | 49995_71 | sp\|Q80TP3\|UBR5_MOUSE | 3.28e-55 |
| contig_373 | 836409 | 34167_63 | sp\|Q91ZX7\|LRP1_MOUSE | 8.15e-58 |
| contig_1054 | 57215 | 1713_43 | sp\|O95757\|HS74L_HUMAN | 2.68e-33 |
| contig_4210 | 398717 | 41263_64 | sp\|Q8CFX8\|SYAC_MESAU | 1.87e-82 |
| contig_544 | 67919 | 47795_32 | sp\|Q9BQS8\|FYCO1_HUMAN | 4.69e-21 |
| contig_629 | 78424 | 51004_17 | sp\|Q9P2E7\|PCD10_HUMAN | 0.0 |
| contig_1032 | 635513 | 1157_77 | sp\|Q1ECW2\|NPS4A_DANRE | 6.68e-66 |
| contig_2300 | 81171 | 16400_31 | sp\|P84039\|ENPP5_RAT | 1.16e-97 |
| contig_1073 | 289775 | 2205_49 | sp\|P15056\|BRAF_HUMAN | 3.52e-19 |
| contig_539 | 392009 | 47400_51 | sp\|P82458\|TRI18_RAT | 2.88e-56 |
| contig_3278 | 234973 | 27614_24 | sp\|Q9IAB2\|MTG8R_XENLA | 2.04e-30 |
| contig_1448 | 2755 | 8271_36 | sp\|Q8N9B5\|JMY_HUMAN | 1.88e-43 |
| contig_1418 | 146165 | 7701_32 | sp\|Q6DRD4\|CENPL_DANRE | 2.58e-76 |
| contig_6626 | 49405 | 52189_8 | sp\|Q7T076\|LSM11_XENLA | 1.06e-41 |
| contig_4149 | 167344 | 40247_47 | sp\|P13821\|SANT_PLAFW | 9.88e-18 |
| contig_2266 | 77167 | 16043_34 | sp\|Q8BLU0\|FLRT2_MOUSE | 1.84e-164 |
| contig_2028 | 100218 | 13916_10 | sp\|Q8R2J9\|RA51C_CRIGR | 6.24e-25 |
| contig_3648 | 25900 | 32889_10 | sp\|Q9DEN4\|FXD3A_XENLA | 1.31e-128 |
| contig_3705 | 53045 | 33585_23 | sp\|A1A5H8\|YES_DANRE | 1.44e-37 |
| contig_331 | 454073 | 28147_31 | sp\|Q54G14\|Y0266_DICDI | 5.92e-13 |
| contig_1235 | 142651 | 4920_19 | sp\|Q91398\|CSK2B_DANRE | 1.96e-48 |
| contig_373 | 288516 | 34103_20 | sp\|P36896\|ACV1B_HUMAN | 9.38e-42 |
| contig_40 | 114114 | 39448_38 | sp\|Q95K09\|GCP5_MACFA | 7.24e-53 |
| contig_5476 | 44720 | 47944_11 | sp\|Q6H236\|PEG3_BOVIN | 5.22e-18 |
| contig_2855 | 20848 | 22386_9 | sp\|P09815\|ICEN_PSEFL | 1.04e-17 |
| contig_345 | 115492 | 30008_7 | sp\|O00370\|LORF2_HUMAN | 5.34e-13 |
| scaffold_4014 | 584911 | 62920_57 | sp\|Q9Z1T6\|FYV1_MOUSE | 6.19e-48 |
| contig_80 | 572495 | 56682_16 | sp\|Q12967\|GNDS_HUMAN | 3.27e-39 |
| contig_1619 | 447465 | 10392_61 | sp\|O95405\|ZFYV9_HUMAN | 1.18e-35 |
| contig_1326 | 25680 | 5619_31 | sp\|P24399\|ZN239_MOUSE | 9.53e-35 |
| contig_7550 | 55387 | 55048_8 | sp\|Q5SSH7\|ZZEF1_MOUSE | 6.61e-58 |
| contig_2281 | 43881 | 16163_56 | sp\|F1QBY1\|NIPLB_DANRE | 6.44e-15 |
| contig_2396 | 817858 | 17672_26 | sp\|Q4V7R1\|TIM9_XENLA | 7.27e-32 |
| contig_479 | 1890665 | 44724_57 | sp\|Q96M86\|DNHD1_HUMAN | 2.53e-14 |
| contig_1351 | 169056 | 6195_5 | sp\|Q5SRE5\|NU188_HUMAN | 2.04e-24 |
| contig_116 | 89351 | 3900_44 | sp\|Q12788\|TBL3_HUMAN | 1.06e-19 |
| scaffold_1156 | 341925 | 61829_8 | sp\|Q80W93\|HYDIN_MOUSE | 1.99e-26 |
| contig_545 | 106589 | 47807_46 | sp\|P22331\|OPSG2_ASTFA | 1.95e-55 |
| contig_5712 | 360874 | 48871_69 | sp\|Q08C99\|VRTN_DANRE | 0.0 |
| contig_454 | 114181 | 43395_71 | sp\|Q15283\|RASA2_HUMAN | 5.00e-17 |
| contig_1585 | 30658 | 10177_63 | sp\|O95696\|BRD1_HUMAN | 2.12e-115 |
| contig_682 | 143542 | 52606_31 | sp\|Q921C5\|BICD2_MOUSE | 1.49e-83 |
| contig_407 | 536590 | 39194_9 | sp\|Q03164\|KMT2A_HUMAN | 9.50e-41 |
| contig_5557 | 91724 | 48234_28 | sp\|Q5D1E7\|ZC12A_MOUSE | 5.49e-51 |
| contig_507 | 21150 | 46120_75 | sp\|Q3U481\|MFS12_MOUSE | 5.56e-15 |
| contig_3712 | 128128 | 33753_78 | sp\|F1QAJ4\|GATA_DANRE | 8.58e-38 |
| contig_1811 | 10091 | 11800_67 | sp\|Q9UFH2\|DYH17_HUMAN | 2.03e-21 |
| contig_141 | 351861 | 7780_22 | sp\|Q96N64\|PWP2A_HUMAN | 1.58e-26 |
| contig_989 | 183247 | 61195_68 | sp\|P55937\|GOGA3_MOUSE | 3.04e-47 |
| contig_479 | 2301136 | 44795_69 | sp\|E9PZ19\|TUTLB_MOUSE | 1.30e-60 |
| contig_3015 | 29544 | 24455_7 | sp\|P35072\|TCB1_CAEBR | 7.24e-30 |
| contig_323 | 176022 | 27233_63 | sp\|P35072\|TCB1_CAEBR | 3.28e-34 |
| contig_1666 | 23315 | 10719_57 | sp\|P35072\|TCB1_CAEBR | 7.65e-44 |
| contig_896 | 278014 | 59305_13 | sp\|D2H0G5\|TBD2A_AILME | 8.52e-41 |
| contig_732 | 187592 | 54341_43 | sp\|F1RD40\|MYCB2_DANRE | 0.0 |
| contig_3055 | 181108 | 24884_37 | sp\|Q5R3F8\|PPR29_HUMAN | 2.24e-119 |
| contig_732 | 209160 | 54344_70 | sp\|Q9UKT7\|FBXL3_HUMAN | 1.01e-94 |
| contig_3793 | 159740 | 34950_73 | sp\|Q91V09\|WDR13_MOUSE | 2.49e-74 |
| contig_735 | 88054 | 54439_35 | sp\|P11029\|ACAC_CHICK | 6.94e-62 |
| contig_511 | 59951 | 46356_41 | sp\|Q5T5M9\|CCNJ_HUMAN | 2.86e-89 |
| contig_2330 | 103455 | 16908_63 | sp\|A6QLN9\|VWA1_BOVIN | 1.97e-34 |
| contig_29 | 135067 | 24268_28 | sp\|Q6NZJ6\|IF4G1_MOUSE | 4.86e-18 |
| contig_539 | 903201 | 47462_76 | sp\|P02542\|DESM_CHICK | 9.42e-48 |
| contig_1450 | 811072 | 8509_75 | sp\|Q29000\|IGF1R_PIG | 5.20e-29 |
| contig_384 | 139056 | 35942_7 | sp\|P70398\|USP9X_MOUSE | 1.11e-39 |
| contig_3124 | 335226 | 25494_51 | sp\|Q9WUV0\|ORC5_MOUSE | 1.34e-25 |
| contig_3750 | 85003 | 34297_39 | sp\|P08169\|MPRI_BOVIN | 5.10e-24 |
| contig_1392 | 192495 | 6978_62 | sp\|Q8N1I0\|DOCK4_HUMAN | 4.04e-30 |
| contig_6383 | 287976 | 51190_21 | sp\|Q505D1\|ANR28_MOUSE | 1.01e-47 |
| contig_2028 | 215644 | 13961_43 | sp\|Q98TZ8\|FLOT2_DANRE | 1.81e-71 |
| contig_588 | 16371 | 49657_45 | sp\|P97789\|XRN1_MOUSE | 1.68e-23 |
| contig_1560 | 280245 | 10054_17 | sp\|P57773\|CXA9_HUMAN | 1.33e-113 |
| contig_3422 | 152416 | 29358_36 | sp\|P16453\|DCHS_RAT | 6.16e-20 |
| contig_3489 | 48726 | 30612_13 | sp\|Q11130\|FUT7_HUMAN | 1.79e-73 |
| scaffold_122 | 4339 | 61972_41 | sp\|Q805F9\|DDB1_CHICK | 1.20e-38 |
| contig_7550 | 234044 | 55015_17 | sp\|Q9BQA5\|HINFP_HUMAN | 4.02e-97 |
| contig_3653 | 33149 | 33114_38 | sp\|Q9WTS4\|TEN1_MOUSE | 0.0 |
| contig_3124 | 346352 | 25495_32 | sp\|Q14789\|GOGB1_HUMAN | 9.09e-12 |
| contig_470 | 138891 | 43888_38 | sp\|O00481\|BT3A1_HUMAN | 1.03e-14 |
| contig_3793 | 91299 | 35040_55 | sp\|Q9H857\|NT5D2_HUMAN | 1.22e-74 |
| contig_445 | 36072 | 43170_64 | sp\|A4Z943\|ZBED5_BOVIN | 2.36e-28 |
| contig_293 | 324812 | 23370_77 | sp\|Q8VDJ3\|VIGLN_MOUSE | 3.72e-71 |
| contig_3551 | 6971 | 31371_62 | sp\|P51050\|MTR1B_CHICK | 1.77e-112 |
| contig_3675 | 48080 | 33310_56 | sp\|Q9NZM4\|BICRA_HUMAN | 3.15e-35 |
| contig_2193 | 424899 | 15180_14 | sp\|O15297\|PPM1D_HUMAN | 3.62e-40 |
| contig_115 | 613246 | 3799_77 | sp\|Q1L8L6\|RN19B_DANRE | 4.44e-46 |
| contig_5608 | 56378 | 48417_27 | sp\|Q99JY3\|GIMA4_MOUSE | 3.60e-17 |
| contig_720 | 148665 | 53986_49 | sp\|Q01702\|DLX3B_DANRE | 3.76e-48 |
| scaffold_4014 | 632369 | 62933_45 | sp\|P63039\|CH60_RAT | 3.75e-56 |
| contig_1103 | 188591 | 2852_69 | sp\|Q7ZVM1\|WASC5_DANRE | 1.63e-43 |
| contig_417 | 147299 | 40650_33 | sp\|A4J5X2\|IF2_DESRM | 1.42e-11 |
| contig_105 | 330356 | 1816_63 | sp\|B5DG67\|WDR12_SALSA | 4.82e-64 |
| contig_1212 | 17818 | 4674_68 | sp\|Q8N328\|PGBD3_HUMAN | 8.02e-11 |
| contig_105 | 458381 | 1851_13 | sp\|Q8K3U6\|FA7_RAT | 8.97e-42 |
| contig_479 | 975267 | 45128_68 | sp\|Q9DDD5\|NBEA_CHICK | 8.30e-59 |
| contig_1054 | 200221 | 1681_58 | sp\|Q38PU4\|GRIK1_MACFA | 2.02e-41 |
| contig_231 | 799004 | 16643_79 | sp\|Q5SZK8\|FREM2_HUMAN | 0.0 |
| contig_2686 | 41738 | 20369_75 | sp\|A5WUN7\|CAM1B_DANRE | 7.54e-128 |
| contig_1129 | 177296 | 3273_58 | sp\|Q9NUV9\|GIMA4_HUMAN | 4.21e-16 |
| contig_1086 | 52748 | 2599_69 | sp\|O35344\|IMA4_MOUSE | 3.84e-47 |
| contig_63 | 192914 | 51252_70 | sp\|Q09575\|YRD6_CAEEL | 1.94e-44 |
| contig_813 | 462528 | 56885_73 | sp\|O60229\|KALRN_HUMAN | 1.46e-120 |
| contig_3470 | 94234 | 30278_11 | sp\|P25104\|AGTR1_BOVIN | 1.07e-41 |
| contig_1170 | 129650 | 3910_32 | sp\|Q803R5\|CMTR1_DANRE | 1.72e-20 |
| contig_2274 | 16738 | 16137_77 | sp\|Q5RHQ8\|LIN9_DANRE | 6.39e-17 |
| contig_867 | 406321 | 58575_72 | sp\|Q7Z494\|NPHP3_HUMAN | 9.84e-66 |
| contig_4005 | 16512 | 38263_41 | sp\|Q7ZWF4\|RN145_DANRE | 3.56e-71 |
| contig_4994 | 107429 | 45892_44 | sp\|Q92072\|DNMT1_CHICK | 0.0 |
| contig_583 | 106718 | 49383_24 | sp\|Q90X49\|CCD80_DANRE | 2.06e-42 |
| contig_1630 | 191890 | 10442_64 | sp\|Q92817\|EVPL_HUMAN | 0.0 |
| contig_29 | 42137 | 24271_73 | sp\|O00400\|ACATN_HUMAN | 1.60e-24 |
| scaffold_6838 | 21564 | 63156_35 | sp\|Q80T02\|GPBAR_RAT | 9.65e-35 |
| contig_3618 | 77818 | 32377_61 | sp\|Q96RD7\|PANX1_HUMAN | 5.85e-38 |
| contig_1337 | 240115 | 5787_62 | sp\|Q9BZR6\|RTN4R_HUMAN | 1.08e-19 |
| contig_3967 | 81237 | 37781_47 | sp\|P10686\|PLCG1_RAT | 5.94e-43 |
| contig_495 | 495829 | 45720_71 | sp\|Q63755\|PRDM2_RAT | 1.08e-130 |
| contig_1054 | 42138 | 1706_13 | sp\|Q7YS78\|ARRC_PIG | 1.22e-16 |
| contig_1665 | 486215 | 10665_42 | sp\|Q8BSS9\|LIPA2_MOUSE | 6.22e-21 |
| scaffold_6136 | 41830 | 63139_17 | sp\|Q9JIY2\|HAKAI_MOUSE | 1.95e-12 |
| contig_1135 | 237368 | 3361_45 | sp\|Q9D620\|RFIP1_MOUSE | 6.44e-22 |
| contig_877 | 50271 | 58899_54 | sp\|P34978\|TA2R_RAT | 5.11e-50 |
| contig_3422 | 135741 | 29355_20 | sp\|P40818\|UBP8_HUMAN | 1.13e-56 |
| contig_271 | 369572 | 20979_34 | sp\|P70031\|CCKAR_XENLA | 1.14e-47 |
| contig_853 | 39157 | 57997_48 | sp\|Q9UKA4\|AKA11_HUMAN | 2.17e-112 |
| contig_480 | 125768 | 45158_71 | sp\|Q09575\|YRD6_CAEEL | 1.75e-16 |
| contig_511 | 1384823 | 46280_76 | sp\|Q3UGY8\|BIG3_MOUSE | 9.02e-46 |
| contig_1385 | 54584 | 6765_23 | sp\|Q96HQ0\|ZN419_HUMAN | 2.13e-30 |
| contig_296 | 70516 | 23792_8 | sp\|O02751\|CFDP2_BOVIN | 1.77e-37 |
| contig_232 | 732673 | 16853_25 | sp\|Q7SXK5\|HUTI_DANRE | 7.59e-69 |
| contig_828 | 54794 | 57423_41 | sp\|Q5TZ24\|MOXD1_DANRE | 6.72e-30 |
| contig_238 | 160556 | 17423_75 | sp\|F1QJX5\|UBR3_DANRE | 6.74e-46 |
| contig_479 | 505906 | 45061_27 | sp\|Q8IZL2\|MAML2_HUMAN | 7.90e-27 |
| contig_1581 | 29711 | 10165_71 | sp\|Q80XG9\|LRRT4_MOUSE | 2.84e-137 |
| contig_1842 | 35831 | 11987_32 | sp\|A4IG55\|PKHO1_DANRE | 4.91e-38 |
| contig_1012 | 81383 | 584_52 | sp\|Q91012\|T22D1_CHICK | 3.71e-33 |
| contig_6067 | 33711 | 50280_32 | sp\|Q5E9Z2\|HABP2_BOVIN | 4.00e-18 |
| scaffold_3356 | 479497 | 62613_26 | sp\|Q9Y663\|HS3SA_HUMAN | 1.64e-125 |
| contig_989 | 181620 | 61191_69 | sp\|P55937\|GOGA3_MOUSE | 1.71e-56 |
| contig_697 | 80580 | 53219_34 | sp\|Q6ZQA6\|IGSF3_MOUSE | 2.51e-35 |
| contig_3426 | 82382 | 29409_60 | sp\|Q6XZF7\|DNMBP_HUMAN | 1.94e-37 |
| contig_2130 | 24600 | 14794_67 | sp\|C0HBT3\|RN146_SALSA | 3.67e-109 |
| contig_40 | 58630 | 39583_75 | sp\|Q6DCC6\|GTPB6_XENLA | 1.98e-34 |
| contig_1215 | 79261 | 4743_74 | sp\|Q7ZW24\|ALG11_DANRE | 1.26e-87 |
| contig_2920 | 16645 | 22949_64 | sp\|Q1LY77\|SE1BA_DANRE | 8.87e-33 |
| contig_539 | 908204 | 47465_53 | sp\|Q5RBT2\|DNPEP_PONAB | 3.92e-19 |
| contig_1001 | 19793 | 3_67 | sp\|P09012\|SNRPA_HUMAN | 1.14e-31 |
| contig_479 | 73401 | 45091_40 | sp\|Q67FY3\|BCL9L_DANRE | 2.13e-110 |
| contig_2585 | 23226 | 19244_19 | sp\|Q02290\|XYNB_NEOPA | 4.12e-19 |
| contig_229 | 64145 | 16345_61 | sp\|Q9Y4B6\|DCAF1_HUMAN | 7.80e-60 |
| contig_3292 | 8314 | 27799_56 | sp\|Q8ILR9\|YPF17_PLAF7 | 5.49e-11 |
| contig_3115 | 27078 | 25371_36 | sp\|P35331\|NRCAM_CHICK | 1.84e-33 |
| contig_3440 | 130051 | 29717_8 | sp\|Q5RBG4\|KAT5_PONAB | 2.12e-33 |
| contig_1351 | 140025 | 6187_11 | sp\|Q80WG5\|LRC8A_MOUSE | 0.0 |
| contig_1377 | 164358 | 6620_13 | sp\|Q8TCN5\|ZN507_HUMAN | 1.83e-70 |
| contig_1619 | 33211 | 10377_76 | sp\|A2AAJ9\|OBSCN_MOUSE | 5.31e-29 |
| contig_2989 | 136317 | 24098_37 | sp\|G5E8K5\|ANK3_MOUSE | 1.90e-64 |
| contig_275 | 74599 | 21581_10 | sp\|Q9BXT8\|RNF17_HUMAN | 7.68e-20 |
| contig_6626 | 219058 | 52154_62 | sp\|Q9Z1J2\|NEK4_MOUSE | 1.09e-65 |
| contig_857 | 131976 | 58067_10 | sp\|Q921G8\|GCP2_MOUSE | 2.22e-38 |
| contig_983 | 14890 | 60995_22 | sp\|Q92090\|CP2K1_ONCMY | 2.08e-47 |
| contig_350 | 36215 | 30947_21 | sp\|P35072\|TCB1_CAEBR | 1.32e-50 |
| contig_2070 | 223476 | 14358_43 | sp\|Q29RH3\|NUD12_BOVIN | 5.98e-38 |
| contig_2924 | 386900 | 23050_74 | sp\|Q7SY48\|HEAT1_DANRE | 4.42e-96 |
| contig_3793 | 104367 | 34943_20 | sp\|Q8TD16\|BICD2_HUMAN | 6.26e-49 |
| contig_115 | 611484 | 3798_31 | sp\|Q1L8L6\|RN19B_DANRE | 4.44e-46 |
| contig_722 | 26630 | 54102_67 | sp\|Q8NEG4\|FA83F_HUMAN | 3.27e-37 |
| contig_1407 | 662893 | 7400_54 | sp\|Q9NX36\|DJC28_HUMAN | 9.11e-92 |
| contig_4441 | 21344 | 43121_55 | sp\|B5X165\|SMG9_SALSA | 4.73e-48 |
| contig_919 | 230556 | 59779_56 | sp\|B5X3Z6\|LIS1A_SALSA | 1.66e-40 |
| contig_1475 | 59599 | 9004_64 | sp\|P35789\|ZNF93_HUMAN | 9.21e-87 |
| contig_3207 | 34893 | 26740_71 | sp\|Q8JH47\|CDK8_DANRE | 2.38e-30 |
| contig_510 | 81386 | 46191_27 | sp\|Q5R5X9\|SMYD4_PONAB | 1.50e-23 |
| contig_4241 | 316276 | 41862_73 | sp\|Q8R066\|C1QT4_MOUSE | 3.86e-112 |
| contig_4098 | 155711 | 39386_67 | sp\|Q69ZH9\|RHG23_MOUSE | 4.63e-30 |
| contig_3455 | 34019 | 29990_13 | sp\|Q08BD8\|MTRF2_DANRE | 2.26e-17 |
| contig_1684 | 32278 | 10846_56 | sp\|Q9Y5Y4\|PD2R2_HUMAN | 3.40e-49 |
| contig_2384 | 12951 | 17333_47 | sp\|Q5R9K1\|CLD12_PONAB | 9.73e-34 |
| contig_437 | 38179 | 43044_52 | sp\|B9EJ80\|PDZD8_MOUSE | 0.0 |
| contig_588 | 20176 | 49663_15 | sp\|Q8IZH2\|XRN1_HUMAN | 2.71e-69 |
| contig_40 | 181030 | 39477_63 | sp\|O95714\|HERC2_HUMAN | 1.18e-27 |
| contig_530 | 364441 | 46976_8 | sp\|Q9QYV0\|ADA15_RAT | 2.33e-16 |
| contig_1135 | 43990 | 3416_37 | sp\|B9U3F2\|DISP3_CHICK | 4.23e-19 |
| contig_989 | 182021 | 61193_75 | sp\|P55937\|GOGA3_MOUSE | 1.66e-55 |
| contig_617 | 1159028 | 50544_70 | sp\|Q7TNE1\|SUCHY_MOUSE | 1.16e-24 |
| contig_692 | 322447 | 52958_26 | sp\|Q4ADV7\|RIC1_HUMAN | 4.66e-112 |
| contig_699 | 361838 | 53296_52 | sp\|Q9P0T7\|TMEM9_HUMAN | 1.62e-30 |
| scaffold_1939 | 205872 | 62178_28 | sp\|Q5TF21\|SOGA3_HUMAN | 1.41e-52 |
| contig_2753 | 193787 | 21441_62 | sp\|E7FAW3\|NBEL2_DANRE | 4.15e-32 |
| contig_1517 | 4017 | 9569_43 | sp\|P35072\|TCB1_CAEBR | 9.60e-43 |
| contig_3706 | 11589 | 33594_12 | sp\|C9ZN16\|FAZ1_TRYB9 | 8.31e-18 |
| contig_134 | 210863 | 6135_25 | sp\|P14315\|CAPZB_CHICK | 2.20e-11 |
| contig_1209 | 826 | 4634_32 | sp\|Q9NUD5\|ZCHC3_HUMAN | 5.04e-24 |
| contig_3402 | 40753 | 29082_8 | sp\|Q5RJ20\|AP2D_DANRE | 1.05e-84 |
| contig_1388 | 44454 | 6797_46 | sp\|Q8IZJ3\|CPMD8_HUMAN | 1.72e-36 |
| contig_479 | 3186337 | 44982_52 | sp\|Q8WU58\|F222B_HUMAN | 7.94e-48 |
| contig_1389 | 278004 | 6852_55 | sp\|Q9NXZ2\|DDX43_HUMAN | 2.61e-38 |
| contig_3561 | 28197 | 31437_71 | sp\|Q8C1B2\|PARPT_MOUSE | 2.45e-41 |
| contig_5380 | 5978 | 47160_16 | sp\|P35072\|TCB1_CAEBR | 8.09e-24 |
| contig_1012 | 46391 | 576_53 | sp\|Q49538\|VLPF_MYCHR | 2.80e-14 |
| contig_479 | 2182338 | 44780_19 | sp\|Q90473\|HSP7C_DANRE | 4.38e-46 |
| contig_5829 | 34120 | 49377_14 | sp\|P53448\|ALDOC_CARAU | 2.08e-41 |
| contig_236 | 165470 | 17206_74 | sp\|Q9Y4G6\|TLN2_HUMAN | 2.21e-21 |
| contig_3958 | 21821 | 37615_5 | sp\|Q08AD1\|CAMP2_HUMAN | 5.83e-29 |
| contig_602 | 39798 | 50155_31 | sp\|Q7ZUW6\|WIPI3_DANRE | 3.53e-22 |
| contig_4257 | 11145 | 42084_56 | sp\|P35072\|TCB1_CAEBR | 3.14e-52 |
| contig_893 | 171344 | 59262_40 | sp\|Q14667\|K0100_HUMAN | 4.32e-98 |
| contig_1054 | 45222 | 1710_61 | sp\|P36575\|ARRC_HUMAN | 8.83e-20 |
| contig_732 | 170917 | 54333_39 | sp\|F1RD40\|MYCB2_DANRE | 6.09e-32 |
| contig_2021 | 83867 | 13888_37 | sp\|P10401\|POLY_DROME | 1.77e-33 |
| contig_620 | 137018 | 50785_34 | sp\|Q3UY90\|GAK1A_MOUSE | 5.93e-22 |
| contig_2385 | 28598 | 17342_17 | sp\|Q6UX07\|DHR13_HUMAN | 5.82e-18 |
| contig_5002 | 283857 | 45966_46 | sp\|Q02241\|KIF23_HUMAN | 2.34e-84 |
| contig_3173 | 155261 | 25892_35 | sp\|Q1LV15\|DAW1_DANRE | 4.01e-37 |
| contig_4065 | 299198 | 38931_26 | sp\|Q25434\|FP1_MYTCO | 7.37e-14 |
| scaffold_3795 | 304838 | 62721_47 | sp\|Q5I598\|MTHR_BOVIN | 8.72e-33 |
| contig_1577 | 11828 | 10151_28 | sp\|Q501V0\|KPSH1_DANRE | 1.72e-180 |
| scaffold_1075 | 71357 | 61649_71 | sp\|P48442\|CASR_RAT | 1.15e-49 |
| contig_1237 | 42131 | 4949_11 | sp\|A4Z944\|ZBED5_CANLF | 7.31e-14 |
| contig_3586 | 24090 | 31815_27 | sp\|P35574\|GDE_RABIT | 2.07e-36 |
| contig_4827 | 16572 | 45208_54 | sp\|Q68EH8\|FAD1_DANRE | 3.58e-20 |
| contig_4145 | 134995 | 40090_24 | sp\|P97279\|ITIH2_MESAU | 7.06e-17 |
| contig_4163 | 87068 | 40451_6 | sp\|Q9BYZ6\|RHBT2_HUMAN | 1.98e-179 |
| contig_4155 | 43094 | 40333_68 | sp\|Q9I8K7\|CITE3_CHICK | 2.06e-25 |
| contig_1392 | 161356 | 6971_23 | sp\|P35072\|TCB1_CAEBR | 3.48e-32 |
| contig_2622 | 193302 | 19612_68 | sp\|O97859\|NEUR3_BOVIN | 4.08e-103 |
| contig_1129 | 97678 | 3301_59 | sp\|Q684M4\|KEAP1_PIG | 2.44e-117 |
| contig_105 | 726437 | 1907_45 | sp\|P29074\|PTN4_HUMAN | 9.66e-32 |
| contig_158 | 112642 | 10180_51 | sp\|F1R345\|DDX11_DANRE | 1.92e-51 |
| contig_1662 | 12339 | 10573_54 | sp\|P16423\|POLR_DROME | 1.57e-16 |
| contig_2940 | 176231 | 23447_62 | sp\|Q3UHK3\|GREB1_MOUSE | 8.00e-30 |
| contig_515 | 92657 | 46518_7 | sp\|A5A6J2\|DDX5_PANTR | 4.40e-18 |
| contig_2253 | 169947 | 15947_18 | sp\|O15040\|TCPR2_HUMAN | 1.58e-28 |
| contig_3328 | 116321 | 28247_11 | sp\|Q8BHN0\|PPM1L_MOUSE | 5.90e-73 |
| contig_2656 | 4970 | 20140_73 | sp\|Q10567\|AP1B1_HUMAN | 5.70e-37 |
| contig_319 | 557971 | 26276_57 | sp\|Q8TD57\|DYH3_HUMAN | 1.22e-24 |
| contig_4561 | 3057 | 43407_22 | sp\|O42400\|AXIN1_CHICK | 4.47e-46 |
| contig_320 | 329579 | 26800_10 | sp\|Q9YGM4\|CAMKV_TAKRU | 1.68e-19 |
| contig_1146 | 432014 | 3601_60 | sp\|C0HL13\|LRP2_PIG | 2.68e-62 |
| contig_3422 | 91262 | 29388_63 | sp\|A3KQQ9\|SCG3_DANRE | 3.65e-30 |
| contig_3672 | 127829 | 33259_14 | sp\|Q9C0C7\|AMRA1_HUMAN | 2.37e-21 |
| contig_4053 | 81805 | 38727_21 | sp\|Q8VDU5\|SNRK_MOUSE | 4.84e-125 |
| contig_3612 | 35003 | 32254_67 | sp\|Q696W0\|SPEG_DANRE | 6.62e-22 |
| contig_2257 | 17879 | 16001_75 | sp\|Q03400\|SANT_PLAF7 | 5.11e-43 |
| contig_696 | 239455 | 53176_29 | sp\|A2AWA9\|RBGP1_MOUSE | 1.98e-56 |
| contig_955 | 4339 | 60629_16 | sp\|Q9QX15\|CA3A1_MOUSE | 3.24e-27 |
| contig_3452 | 241922 | 29917_42 | sp\|P23440\|PDE6B_MOUSE | 6.05e-52 |
| contig_6163 | 18412 | 50483_79 | sp\|O15553\|MEFV_HUMAN | 2.08e-27 |
| contig_1323 | 121378 | 5459_53 | sp\|O35149\|ZNT4_MOUSE | 9.91e-17 |
| contig_1485 | 181180 | 9138_32 | sp\|O93574\|RELN_CHICK | 1.15e-43 |
| contig_3361 | 12342 | 28639_50 | sp\|Q9Y4D8\|HECD4_HUMAN | 3.72e-51 |
| scaffold_1939 | 129706 | 62170_75 | sp\|Q5RGU1\|COQ8A_DANRE | 1.79e-15 |
| contig_1032 | 624292 | 1154_13 | sp\|O55058\|FBLN4_CRIGR | 3.96e-45 |
| contig_3797 | 43215 | 35087_66 | sp\|Q92621\|NU205_HUMAN | 9.53e-25 |
| contig_319 | 1582662 | 26222_39 | sp\|Q8CDU6\|HECD2_MOUSE | 7.13e-28 |
| contig_3865 | 79963 | 36202_73 | sp\|B3DIY3\|MMS22_DANRE | 7.10e-14 |
| contig_472 | 478933 | 44032_13 | sp\|O60443\|GSDME_HUMAN | 1.16e-29 |
| contig_34 | 648014 | 30811_75 | sp\|Q61830\|MRC1_MOUSE | 4.95e-14 |
| contig_3994 | 497160 | 38082_70 | sp\|Q6P4T2\|U520_MOUSE | 4.21e-66 |
| contig_2472 | 15686 | 18229_17 | sp\|Q5SWW4\|MED13_MOUSE | 6.14e-35 |
| contig_6957 | 18334 | 53133_14 | sp\|Q7M732\|RTL1_MOUSE | 2.18e-11 |
| contig_272 | 50408 | 21197_74 | sp\|O35264\|PA1B2_RAT | 1.64e-32 |
| contig_3748 | 19851 | 34250_39 | sp\|Q9P2E2\|KIF17_HUMAN | 2.27e-51 |
| contig_375 | 392523 | 34565_47 | sp\|Q00725\|SGS4_DROME | 6.66e-15 |
| contig_2881 | 7608 | 22601_8 | sp\|P03934\|TC1A_CAEEL | 1.62e-43 |
| contig_4371 | 25668 | 43000_45 | sp\|P51513\|NOVA1_HUMAN | 2.65e-125 |
| contig_4720 | 143943 | 43975_63 | sp\|A5WWB6\|ZWILC_DANRE | 2.87e-32 |
| contig_1876 | 93886 | 12547_14 | sp\|P33176\|KINH_HUMAN | 4.13e-33 |
| contig_3745 | 24426 | 34239_55 | sp\|C9JH25\|PRRT4_HUMAN | 1.20e-13 |
| contig_3328 | 76193 | 28266_57 | sp\|Q9NTJ3\|SMC4_HUMAN | 2.12e-49 |
| contig_4171 | 36190 | 40566_16 | sp\|Q90923\|NET3_CHICK | 8.33e-44 |
| contig_452 | 88553 | 43385_5 | sp\|Q6NV31\|WDR82_DANRE | 4.24e-73 |
| contig_2924 | 294849 | 23019_21 | sp\|B0LPN4\|RYR2_RAT | 4.43e-59 |
| contig_4016 | 49856 | 38371_43 | sp\|Q8CI94\|PYGB_MOUSE | 4.41e-31 |
| contig_411 | 327865 | 39857_22 | sp\|A2AX52\|CO6A4_MOUSE | 1.01e-74 |
| contig_1417 | 80519 | 7685_13 | sp\|Q5SZK8\|FREM2_HUMAN | 0.0 |
| contig_6690 | 6013 | 52323_24 | sp\|P51513\|NOVA1_HUMAN | 2.36e-48 |
| contig_19 | 22865 | 13758_56 | sp\|P0CT40\|TF29_SCHPO | 1.50e-38 |
| contig_364 | 494124 | 33015_55 | sp\|A0JM56\|LRRC9_XENTR | 9.39e-37 |
| contig_431 | 112457 | 42666_78 | sp\|Q9Y2E4\|DIP2C_HUMAN | 5.75e-57 |
| contig_1455 | 227550 | 8589_65 | sp\|Q9NRM2\|ZN277_HUMAN | 9.15e-26 |
| contig_831 | 125674 | 57597_22 | sp\|Q95SX7\|RTBS_DROME | 3.15e-23 |
| contig_1449 | 184328 | 8300_78 | sp\|Q5U395\|CNEPA_DANRE | 4.27e-51 |
| contig_479 | 1890916 | 44725_35 | sp\|Q96M86\|DNHD1_HUMAN | 2.53e-14 |
| contig_2569 | 24875 | 19045_16 | sp\|Q9JIP3\|I17RB_MOUSE | 3.06e-12 |
| contig_2753 | 158675 | 21429_58 | sp\|Q501X2\|CEP72_DANRE | 8.72e-30 |
| contig_1184 | 322899 | 4155_36 | sp\|Q80WQ9\|ZBED4_MOUSE | 4.86e-13 |
| contig_697 | 28260 | 53212_29 | sp\|P35072\|TCB1_CAEBR | 1.58e-60 |
| contig_2753 | 190157 | 21439_73 | sp\|E7FAW3\|NBEL2_DANRE | 1.86e-30 |
| contig_80 | 112771 | 56541_31 | sp\|Q8WX92\|NELFB_HUMAN | 1.36e-31 |
| contig_3832 | 450067 | 35533_64 | sp\|Q5RBC8\|CMC1_PONAB | 4.91e-56 |
| contig_617 | 239479 | 50579_49 | sp\|Q9QZ04\|MAGL2_MOUSE | 2.60e-13 |
| contig_338 | 39944 | 28981_47 | sp\|Q02391\|GSLG1_CHICK | 7.77e-31 |
| contig_2042 | 14454 | 14154_46 | sp\|Q1XH10\|SKDA1_HUMAN | 1.47e-62 |
| contig_718 | 147902 | 53909_35 | sp\|Q9Y263\|PLAP_HUMAN | 6.78e-56 |
| contig_799 | 365380 | 56461_11 | sp\|Q9CU65\|ZMYM2_MOUSE | 1.12e-12 |
| contig_1533 | 63618 | 9724_14 | sp\|E7FCN8\|INTU_DANRE | 3.18e-47 |
| contig_818 | 175479 | 57035_62 | sp\|P19336\|CYR61_CHICK | 2.63e-34 |
| contig_3837 | 608932 | 35801_24 | sp\|Q8K4G1\|LTBP4_MOUSE | 2.09e-65 |
| contig_3946 | 149887 | 37145_15 | sp\|Q96JK4\|HIPL1_HUMAN | 2.24e-69 |
| contig_1507 | 183307 | 9445_47 | sp\|O14792\|HS3S1_HUMAN | 1.19e-83 |
| contig_931 | 94193 | 60097_13 | sp\|P16295\|FA9_CAVPO | 1.50e-41 |
| contig_3428 | 231642 | 29446_45 | sp\|Q8NDN9\|RCBT1_HUMAN | 2.94e-44 |
| contig_2046 | 52252 | 14186_79 | sp\|O35678\|MGLL_MOUSE | 8.88e-21 |
| contig_373 | 740535 | 34159_58 | sp\|A0N0X6\|LRRN1_BOVIN | 0.0 |
| contig_1450 | 162240 | 8423_49 | sp\|Q9R049\|AMFR_MOUSE | 8.81e-26 |
| contig_830 | 145959 | 57488_67 | sp\|O57409\|DLLB_DANRE | 6.91e-61 |
| contig_580 | 1276821 | 49215_47 | sp\|P84239\|H3_URECA | 1.34e-57 |
| contig_130 | 560840 | 5325_59 | sp\|Q6KAS7\|ZN521_MOUSE | 0.0 |
| contig_199 | 69314 | 13745_73 | sp\|Q5RKJ0\|F110C_RAT | 2.36e-22 |
| contig_118 | 426615 | 4336_28 | sp\|Q9CWH1\|ZBT8A_MOUSE | 3.05e-102 |
| contig_1485 | 160285 | 9132_53 | sp\|O75747\|P3C2G_HUMAN | 4.69e-24 |
| contig_1027 | 456416 | 879_33 | sp\|P35072\|TCB1_CAEBR | 9.00e-23 |
| contig_1859 | 87624 | 12125_39 | sp\|O35732\|CFLAR_MOUSE | 1.81e-23 |
| contig_497 | 17889 | 45780_29 | sp\|Q53R41\|FAKD1_HUMAN | 5.33e-66 |
| contig_3449 | 58648 | 29849_5 | sp\|Q6WRI0\|IGS10_HUMAN | 1.15e-94 |
| scaffold_992 | 353757 | 63374_21 | sp\|Q96HN2\|SAHH3_HUMAN | 2.15e-26 |
| contig_3154 | 241020 | 25703_33 | sp\|Q60809\|CNOT7_MOUSE | 2.37e-55 |
| contig_416 | 892288 | 40548_75 | sp\|P21328\|RTJK_DROME | 2.64e-18 |
| contig_762 | 82853 | 55445_54 | sp\|Q923L3\|CSMD1_MOUSE | 1.72e-37 |
| contig_2206 | 144546 | 15476_42 | sp\|Q8TE73\|DYH5_HUMAN | 5.27e-75 |
| contig_1456 | 119136 | 8626_49 | sp\|Q6DRC5\|USPL1_DANRE | 2.21e-11 |
| contig_5270 | 80718 | 46871_27 | sp\|P51030\|WNT8C_CHICK | 7.59e-93 |
| contig_3463 | 62545 | 30076_46 | sp\|Q1XI86\|IMPG2_CHICK | 6.75e-47 |
| contig_4825 | 2428 | 45203_55 | sp\|Q5R7Y0\|AGRB2_PONAB | 5.95e-39 |
| contig_989 | 856303 | 61333_30 | sp\|P23378\|GCSP_HUMAN | 6.46e-32 |
| contig_7621 | 15150 | 55413_22 | sp\|Q5ZMQ9\|PEX5_CHICK | 1.64e-16 |
| contig_979 | 483726 | 60857_68 | sp\|Q62018\|CTR9_MOUSE | 2.05e-131 |
| contig_278 | 47602 | 21811_34 | sp\|O60437\|PEPL_HUMAN | 8.25e-13 |
| contig_989 | 233184 | 61205_12 | sp\|A0A087WPF7\|AUTS2_MOUSE | 1.77e-11 |
| contig_1480 | 42362 | 9070_66 | sp\|A2AWL7\|MGAP_MOUSE | 4.67e-35 |
| contig_3672 | 13280 | 33267_24 | sp\|O60290\|ZN862_HUMAN | 1.99e-57 |
| contig_989 | 614226 | 61290_69 | sp\|A6P7L8\|PIWL2_ONCMY | 6.09e-39 |
| contig_2707 | 70952 | 20836_38 | sp\|Q96LT4\|SAMD8_HUMAN | 1.59e-56 |
| contig_3455 | 183313 | 29975_7 | sp\|Q6NWF1\|GTR12_DANRE | 4.23e-75 |
| contig_546 | 82111 | 47879_44 | sp\|Q8K3W2\|LRC10_MOUSE | 8.25e-91 |
| contig_411 | 259587 | 39840_23 | sp\|Q6EDY6\|CARL1_MOUSE | 3.51e-24 |
| contig_3008 | 89672 | 24385_8 | sp\|Q96I24\|FUBP3_HUMAN | 2.58e-22 |
| contig_1215 | 155834 | 4713_64 | sp\|Q8NF99\|ZN397_HUMAN | 3.43e-31 |
| contig_1213 | 54324 | 4684_35 | sp\|A8E0Y8\|IGSF2_MOUSE | 2.02e-29 |
| contig_545 | 243236 | 47815_46 | sp\|O88632\|SEM3F_MOUSE | 7.33e-25 |
| contig_3755 | 46893 | 34357_76 | sp\|P30994\|5HT2B_RAT | 1.40e-41 |
| contig_2901 | 85962 | 22807_54 | sp\|Q9NVR0\|KLH11_HUMAN | 2.18e-47 |
| scaffold_71 | 82146 | 63202_11 | sp\|Q5CZR5\|AEP1_DANRE | 5.61e-110 |
| contig_489 | 119079 | 45373_63 | sp\|Q2QCI8\|MED12_DANRE | 7.74e-47 |
| contig_1557 | 2472 | 9961_42 | sp\|Q15428\|SF3A2_HUMAN | 4.64e-19 |
| contig_1215 | 77714 | 4741_54 | sp\|Q7ZW24\|ALG11_DANRE | 1.26e-87 |
| contig_1046 | 620124 | 1537_20 | sp\|Q6GN29\|ATLA2_XENLA | 2.75e-26 |
| contig_575 | 86716 | 49069_5 | sp\|Q9GLP1\|FA5_PIG | 1.84e-22 |
| contig_3612 | 58689 | 32261_8 | sp\|P05548\|CAVPT_BRALA | 1.30e-16 |
| contig_2058 | 175549 | 14268_70 | sp\|Q9Y2L6\|FRM4B_HUMAN | 6.15e-16 |
| contig_4994 | 172228 | 45900_62 | sp\|Q568Y7\|NOE2_RAT | 2.09e-104 |
| contig_479 | 2209211 | 44784_64 | sp\|Q9HCK4\|ROBO2_HUMAN | 5.19e-56 |
| contig_4994 | 36713 | 45905_55 | sp\|P29597\|TYK2_HUMAN | 2.04e-26 |
| contig_1103 | 188358 | 2851_48 | sp\|Q7ZVM1\|WASC5_DANRE | 1.63e-43 |
| contig_130 | 960608 | 5392_6 | sp\|P46101\|DPP6_RAT | 1.02e-34 |
| contig_1406 | 163181 | 7291_47 | sp\|Q9H799\|CPLN1_HUMAN | 1.29e-22 |
| contig_436 | 247331 | 42973_67 | sp\|Q5T011\|SZT2_HUMAN | 5.46e-31 |
| contig_1490 | 106797 | 9189_35 | sp\|Q9ERS6\|IRPL2_MOUSE | 4.67e-81 |
| contig_3723 | 257911 | 33920_59 | sp\|Q9NPA5\|ZF64A_HUMAN | 5.11e-80 |
| contig_351 | 137781 | 31033_26 | sp\|Q9QYC5\|G37L1_RAT | 9.81e-51 |
| contig_813 | 540497 | 56895_47 | sp\|P22727\|WNT6_MOUSE | 3.29e-32 |
| contig_105 | 341588 | 1818_55 | sp\|P28331\|NDUS1_HUMAN | 4.08e-71 |
| contig_1842 | 105431 | 11968_13 | sp\|Q49GP3\|PI4KB_DANRE | 1.85e-79 |
| contig_792 | 271697 | 56260_42 | sp\|Q63ZU3\|MYADM_XENLA | 4.22e-57 |
| contig_818 | 212233 | 57044_15 | sp\|A2PYH4\|HFM1_HUMAN | 9.80e-60 |
| contig_155 | 64203 | 10021_69 | sp\|Q95YJ5\|TXND3_CIOIN | 7.56e-17 |
| contig_384 | 95530 | 35968_5 | sp\|P53794\|SC5A3_HUMAN | 0.0 |
| contig_3934 | 206891 | 36981_63 | sp\|Q8K224\|NAT10_MOUSE | 1.09e-77 |
| contig_1519 | 23056 | 9600_61 | sp\|Q96WV6\|YHU2_SCHPO | 1.06e-21 |
| contig_556 | 92389 | 48256_68 | sp\|Q5SUR0\|PUR4_MOUSE | 5.20e-37 |
| contig_1521 | 29145 | 9621_25 | sp\|Q28107\|FA5_BOVIN | 1.49e-27 |
| contig_996 | 18241 | 61409_67 | sp\|Q13972\|RGRF1_HUMAN | 3.55e-48 |
| contig_2077 | 2899 | 14472_39 | sp\|Q9JJ79\|DYHC2_RAT | 1.38e-19 |
| contig_861 | 26377 | 58360_57 | sp\|Q96WV6\|YHU2_SCHPO | 1.55e-19 |
| contig_372 | 215998 | 34027_55 | sp\|Q91WJ8\|FUBP1_MOUSE | 1.14e-44 |
| contig_3201 | 229924 | 26530_14 | sp\|P80316\|TCPE_MOUSE | 2.03e-60 |
| contig_364 | 203851 | 32956_65 | sp\|Q8NCM2\|KCNH5_HUMAN | 6.62e-25 |
| contig_80 | 634735 | 56697_78 | sp\|P29274\|AA2AR_HUMAN | 1.01e-71 |
| contig_1540 | 16555 | 9791_67 | sp\|Q9UJX0\|OSGI1_HUMAN | 1.11e-70 |
| contig_511 | 1106147 | 46220_7 | sp\|Q8VI59\|PCX3_MOUSE | 3.30e-28 |
| contig_3612 | 54127 | 32259_40 | sp\|Q696W0\|SPEG_DANRE | 7.69e-59 |
| contig_239 | 193113 | 17736_39 | sp\|Q6Q7X9\|KLH31_DANRE | 0.0 |
| contig_3615 | 176524 | 32287_40 | sp\|O70240\|AXIN2_RAT | 2.91e-19 |
| contig_4115 | 29411 | 39788_23 | sp\|Q8NDT2\|RB15B_HUMAN | 1.20e-136 |
| contig_549 | 112665 | 47960_60 | sp\|Q8BVD7\|C1QT7_MOUSE | 1.21e-44 |
| contig_1088 | 35405 | 2683_26 | sp\|Q8BTZ4\|APC5_MOUSE | 3.68e-25 |
| contig_729 | 263063 | 54194_20 | sp\|Q309B1\|TR16L_HUMAN | 3.93e-61 |
| contig_465 | 16107 | 43577_68 | sp\|Q86MA7\|PRQFV_APLCA | 2.77e-39 |
| contig_403 | 130621 | 38509_28 | sp\|P10394\|POL4_DROME | 1.22e-41 |
| scaffold_4014 | 819961 | 62956_40 | sp\|Q5ZHW4\|RAB5B_CHICK | 2.15e-68 |
| contig_2753 | 195157 | 21442_66 | sp\|E7FAW3\|NBEL2_DANRE | 4.15e-32 |
| contig_185 | 367429 | 12202_53 | sp\|Q2TL32\|UBR4_RAT | 2.77e-104 |
| contig_1025 | 192406 | 784_45 | sp\|Q68ED2\|GRM7_MOUSE | 1.88e-55 |
| contig_2686 | 68386 | 20381_43 | sp\|Q07008\|NOTC1_RAT | 1.43e-57 |
| contig_3211 | 135977 | 26870_77 | sp\|Q6P4R8\|NFRKB_HUMAN | 3.15e-26 |
| contig_2932 | 95210 | 23311_40 | sp\|Q3TCH7\|CUL4A_MOUSE | 7.99e-54 |
| scaffold_2818 | 48657 | 62284_6 | sp\|Q9PU36\|PCLO_CHICK | 2.30e-65 |
| contig_3110 | 147962 | 25280_70 | sp\|O76050\|NEUL1_HUMAN | 2.80e-37 |
| contig_927 | 89803 | 59917_35 | sp\|Q96S38\|KS6C1_HUMAN | 1.16e-32 |
| contig_2924 | 899551 | 23169_66 | sp\|Q5R7U6\|COX11_PONAB | 4.43e-35 |
| contig_4380 | 11212 | 43057_61 | sp\|P11716\|RYR1_RABIT | 5.65e-75 |
| contig_4236 | 59414 | 41710_79 | sp\|O02751\|CFDP2_BOVIN | 8.39e-22 |
| contig_1029 | 97670 | 989_79 | sp\|Q8WXD0\|RXFP2_HUMAN | 1.70e-73 |
| contig_3621 | 50019 | 32433_68 | sp\|Q8BFQ3\|OGR1_MOUSE | 6.37e-75 |
| contig_1011 | 648912 | 495_38 | sp\|Q9Y6N8\|CAD10_HUMAN | 2.58e-36 |
| contig_2080 | 10432 | 14481_41 | sp\|P10394\|POL4_DROME | 1.40e-19 |
| contig_3615 | 113164 | 32276_26 | sp\|Q8IWB9\|TEX2_HUMAN | 1.26e-23 |
| contig_1106 | 99382 | 2942_74 | sp\|P42337\|PK3CA_MOUSE | 4.71e-109 |
| contig_929 | 316096 | 59967_42 | sp\|Q9XWZ2\|ACD11_CAEEL | 6.99e-31 |
| contig_822 | 234069 | 57185_47 | sp\|O15072\|ATS3_HUMAN | 9.74e-41 |
| contig_3231 | 27018 | 27193_53 | sp\|Q6DHE8\|RHOAD_DANRE | 5.16e-54 |
| contig_539 | 896071 | 47457_45 | sp\|Q498W5\|T198B_DANRE | 9.10e-46 |
| contig_1510 | 50461 | 9511_6 | sp\|Q1LVK9\|N4BP1_DANRE | 8.76e-41 |
| contig_495 | 208574 | 45630_31 | sp\|Q8AY73\|XPO2_ORENI | 2.87e-45 |
| contig_3858 | 18269 | 36082_17 | sp\|Q9BYR4\|KRA43_HUMAN | 2.33e-12 |
| contig_186 | 703608 | 12414_14 | sp\|Q99KD5\|UN45A_MOUSE | 4.14e-46 |
| contig_826 | 350495 | 57302_38 | sp\|F1QUP1\|RPZ_DANRE | 5.36e-46 |
| contig_2686 | 61927 | 20376_47 | sp\|A2RUV0\|NOTC1_XENTR | 2.80e-91 |
| contig_2943 | 15438 | 23477_7 | sp\|Q3SY56\|SP6_HUMAN | 5.47e-73 |
| contig_5125 | 89418 | 46445_56 | sp\|Q9DC23\|DJC10_MOUSE | 1.41e-40 |
| contig_3452 | 265696 | 29924_47 | sp\|Q8R2G6\|CCD80_MOUSE | 4.01e-24 |
| contig_1200 | 455405 | 4514_22 | sp\|Q5RCK5\|ASB7_PONAB | 1.62e-123 |
| contig_877 | 529585 | 58904_77 | sp\|Q9BYG0\|B3GN5_HUMAN | 7.36e-160 |
| contig_55 | 225365 | 48328_35 | sp\|Q91W92\|BORG5_MOUSE | 8.39e-21 |
| contig_1692 | 144304 | 10869_66 | sp\|O57321\|EAA1_AMBTI | 3.48e-41 |
| contig_489 | 932327 | 45530_34 | sp\|Q13620\|CUL4B_HUMAN | 3.89e-64 |
| contig_3678 | 28356 | 33355_32 | sp\|Q9NZU0\|FLRT3_HUMAN | 0.0 |
| contig_2330 | 25798 | 16933_74 | sp\|Q62504\|MINT_MOUSE | 2.98e-65 |
| contig_4273 | 97411 | 42304_53 | sp\|Q7Z5L7\|PODN_HUMAN | 5.97e-59 |
| contig_2956 | 117247 | 23642_36 | sp\|Q90854\|KCNJ3_CHICK | 9.51e-128 |
| contig_979 | 683350 | 60915_28 | sp\|G3V7Q0\|DEN5A_RAT | 7.97e-49 |
| contig_1339 | 110702 | 5885_25 | sp\|A1XQY1\|NR3BA_DANRE | 1.41e-105 |
| contig_505 | 43983 | 46081_50 | sp\|Q076A5\|MYH4_CANLF | 5.71e-83 |

**Table S9**. GO enrichment for transcripts located within the 1Kbp window around the outlier loci associated to the type of spawning habitat (beach-spawning sites, shallow-water demersal sites, and deep-water demersal sites)

| id | level | namespace | name | RS | RP | p_fdr |
| --- | --- | --- | --- | --- | --- | --- |
| GO:0098805 | 2 | cellular_component | whole membrane | 16/104 | 1577/29353 | 0.0705 |
| GO:0097472 | 2 | molecular_function | cyclin-dependent protein kinase activity | 4/104 | 40/29353 | 0.0168 |
| GO:0140096 | 2 | molecular_function | catalytic activity, acting on a protein | 28/104 | 3511/29353 | 0.0252 |
| GO:0043167 | 2 | molecular_function | ion binding | 55/104 | 10219/29353 | 0.086 |
| GO:0019538 | 3 | biological_process | protein metabolic process | 36/104 | 4689/29353 | 0.00864 |
| GO:0009268 | 3 | biological_process | response to pH | 4/104 | 48/29353 | 0.0252 |
| GO:0044260 | 3 | biological_process | cellular macromolecule metabolic process | 38/104 | 5767/29353 | 0.0364 |
| GO:0098588 | 3 | cellular_component | bounding membrane of organelle | 25/104 | 2517/29353 | 0.00719 |
| GO:0097708 | 3 | cellular_component | intracellular vesicle | 19/104 | 1755/29353 | 0.0168 |
| GO:0031982 | 3 | cellular_component | vesicle | 25/104 | 2843/29353 | 0.0171 |
| GO:0000151 | 3 | cellular_component | ubiquitin ligase complex | 8/104 | 402/29353 | 0.0554 |
| GO:0019904 | 3 | molecular_function | protein domain specific binding | 17/104 | 1036/29353 | 0.00141 |
| GO:0019787 | 3 | molecular_function | ubiquitin-like protein transferase activity | 13/104 | 636/29353 | 0.00212 |
| GO:0004693 | 3 | molecular_function | cyclin-dependent protein serine/threonine kinase activity | 4/104 | 40/29353 | 0.0168 |

**Table S10**. GO enrichment for transcripts located within the 1Kbp window around the outlier loci associated with temperature.

| id | level | namespace | name | RS | RP | p_fdr |
| --- | --- | --- | --- | --- | --- | --- |
| GO:1902494 | 2 | cellular_component | catalytic complex | 24/76 | 1852/29353 | 2.16e-07 |
| GO:0120114 | 2 | cellular_component | Sm-like protein family complex | 5/76 | 90/29353 | 0.00386 |
| GO:0040008 | 3 | biological_process | regulation of growth | 10/76 | 804/29353 | 0.0205 |
| GO:0009314 | 3 | biological_process | response to radiation | 8/76 | 532/29353 | 0.0273 |
| GO:0031647 | 3 | biological_process | regulation of protein stability | 6/76 | 347/29353 | 0.0748 |
| GO:1990234 | 3 | cellular_component | transferase complex | 15/76 | 1048/29353 | 0.000223 |
| GO:0000151 | 3 | cellular_component | ubiquitin ligase complex | 10/76 | 402/29353 | 0.000259 |
| GO:0030532 | 3 | cellular_component | small nuclear ribonucleoprotein complex | 5/76 | 76/29353 | 0.00257 |
| GO:0072357 | 3 | cellular_component | PTW/PP1 phosphatase complex | 3/76 | 15/29353 | 0.00608 |
| GO:0071013 | 3 | cellular_component | catalytic step 2 spliceosome | 5/76 | 112/29353 | 0.00835 |
| GO:1902710 | 3 | cellular_component | GABA receptor complex | 3/76 | 35/29353 | 0.0354 |
| GO:0031371 | 3 | cellular_component | ubiquitin conjugating enzyme complex | 2/76 | 11/29353 | 0.0853 |
| GO:0019787 | 3 | molecular_function | ubiquitin-like protein transferase activity | 9/76 | 636/29353 | 0.0198 |
| GO:0101005 | 3 | molecular_function | ubiquitinyl hydrolase activity | 5/76 | 160/29353 | 0.0245 |

**Table S11**. GO enrichment for transcripts located within the 1Kbp window around the outlier loci associated with chlorophyll concentration.

| id | level | namespace | name | RS | RP | p_fdr |
| --- | --- | --- | --- | --- | --- | --- |
| GO:0050896 | 1 | biological_process | response to stimulus | 45/126 | 6396/29353 | 0.0746 |
| GO:0043226 | 1 | cellular_component | organelle | 87/126 | 14901/29353 | 0.0155 |
| GO:0043227 | 2 | cellular_component | membrane-bounded organelle | 81/126 | 13083/29353 | 0.00555 |
| GO:0043229 | 2 | cellular_component | intracellular organelle | 83/126 | 13945/29353 | 0.0155 |
| GO:0030055 | 2 | cellular_component | cell-substrate junction | 9/126 | 453/29353 | 0.05 |
| GO:0070161 | 2 | cellular_component | anchoring junction | 11/126 | 701/29353 | 0.0599 |
| GO:0005515 | 2 | molecular_function | protein binding | 63/126 | 9900/29353 | 0.0549 |
| GO:0048798 | 3 | biological_process | swim bladder inflation | 2/126 | 2/29353 | 0.00985 |
| GO:0044770 | 3 | biological_process | cell cycle phase transition | 7/126 | 194/29353 | 0.0107 |
| GO:0050794 | 3 | biological_process | regulation of cellular process | 76/126 | 13089/29353 | 0.0885 |
| GO:0000307 | 3 | cellular_component | cyclin-dependent protein kinase holoenzyme complex | 7/126 | 43/29353 | 3.29e-06 |
| GO:0043231 | 3 | cellular_component | intracellular membrane-bounded organelle | 72/126 | 11647/29353 | 0.0284 |
| GO:0005924 | 3 | cellular_component | cell-substrate adherens junction | 9/126 | 446/29353 | 0.0459 |
| GO:0005912 | 3 | cellular_component | adherens junction | 11/126 | 680/29353 | 0.053 |

**Table S12**. $\chi^{2}$ tests for deviation to Hardy-Weinberg equilibrium for the putative homokaryotes (haplogroup1 and haplogroup2) and the heterokaryote (haplogroup3). Number of individuals assigned to each haplogroup is given (num_haplo1, num_haplo2, num_haplo3).

| pop | num_haplo1 | num_haplo2 | num_haplo3 | $\chi^{2}$ | p-value |
| --- | --- | --- | --- | --- | --- |
| N2 | 25 | 19 | 1 | 0.76025231 | 0.3832496 |
| N3 | 29 | 17 | 2 | 0.01990748 | 0.8877958 |
| L13 | 35 | 13 | 2 | 0.02855261 | 0.8658162 |
| S4 | 22 | 24 | 1 | 2.57027651 | 0.1088887 |
| S6 | 28 | 20 | 1 | 0.76103741 | 0.3830041 |
| S3 | 30 | 14 | 3 | 0.16831629 | 0.6816125 |
| L10 | 30 | 18 | 0 | 1.44532106 | 0.2292802 |
| L11 | 30 | 16 | 1 | 0.09218221 | 0.761421 |
| S1 | 34 | 11 | 0 | 0.11883095 | 0.7303059 |
| L2 | 23 | 15 | 7 | 1.75519811 | 0.1852246 |
| L1 | 7 | 8 | 4 | 0.07073218 | 0.7902737 |
| N4 | 28 | 17 | 3 | 0.02599217 | 0.8719194 |
| L12 | 20 | 22 | 4 | 0.12000347 | 0.7290307 |
| L14 | 17 | 12 | 0 | 0.88597459 | 0.3465705 |
| L6 | 29 | 20 | 1 | 0.69666678 | 0.403906 |
| L7 | 22 | 17 | 6 | 0.41619631 | 0.5188411 |
| L8 | 27 | 10 | 0 | 0.1305957 | 0.7178152 |
| L9 | 26 | 18 | 4 | 0.01122087 | 0.915639 |
| L4 | 7 | 11 | 2 | 0.18222222 | 0.6694704 |
| S2 | 28 | 18 | 1 | 0.39437911 | 0.5300065 |
| L5 | 25 | 20 | 0 | 2.44387755 | 0.1179843 |
| L15 | 26 | 20 | 2 | 0.21759259 | 0.6408804 |
| S8 | 31 | 14 | 4 | 0.85066448 | 0.3563644 |
| S5 | 34 | 12 | 1 | 0.22131059 | 0.6380431 |
| K1 | 23 | 23 | 3 | 0.38371731 | 0.5356205 |

**Table S13**. Genetic differentiation with NWA lineage. We present pairwise $F_{ST}$ values; those for which the p-value was higher than 0.01 are shown in grey.

|  | N2 | N3 | L13 | S4 | S6 | S3 | L10 | L11 | S1 | L2 | N4 | L1 | L12 | L14 | L6 | L7 | L8 | L9 | L4 | S2 | L5 | L15 | S5 | K1 | S8 |
| --- | --- | --- | --- | --- | --- | --- | --- | --- | --- | --- | --- | --- | --- | --- | --- | --- | --- | --- | --- | --- | --- | --- | --- | --- | --- |
| N2 | NA | NA | NA | NA | NA | NA | NA | NA | NA | NA | NA | NA | NA | NA | NA | NA | NA | NA | NA | NA | NA | NA | NA | NA | NA |
| N3 | 0.000 | NA | NA | NA | NA | NA | NA | NA | NA | NA | NA | NA | NA | NA | NA | NA | NA | NA | NA | NA | NA | NA | NA | NA | NA |
| L13 | 0.002 | 0.002 | NA | NA | NA | NA | NA | NA | NA | NA | NA | NA | NA | NA | NA | NA | NA | NA | NA | NA | NA | NA | NA | NA | NA |
| S4 | 0.000 | 0.000 | 0.002 | NA | NA | NA | NA | NA | NA | NA | NA | NA | NA | NA | NA | NA | NA | NA | NA | NA | NA | NA | NA | NA | NA |
| S6 | 0.000 | 0.000 | 0.002 | 0.000 | NA | NA | NA | NA | NA | NA | NA | NA | NA | NA | NA | NA | NA | NA | NA | NA | NA | NA | NA | NA | NA |
| S3 | 0.001 | 0.000 | 0.002 | 0.001 | 0.001 | NA | NA | NA | NA | NA | NA | NA | NA | NA | NA | NA | NA | NA | NA | NA | NA | NA | NA | NA | NA |
| L10 | 0.004 | 0.003 | 0.005 | 0.003 | 0.003 | 0.004 | NA | NA | NA | NA | NA | NA | NA | NA | NA | NA | NA | NA | NA | NA | NA | NA | NA | NA | NA |
| L11 | 0.005 | 0.005 | 0.007 | 0.004 | 0.005 | 0.006 | 0.001 | NA | NA | NA | NA | NA | NA | NA | NA | NA | NA | NA | NA | NA | NA | NA | NA | NA | NA |
| S1 | 0.003 | 0.003 | 0.001 | 0.003 | 0.003 | 0.002 | 0.006 | 0.007 | NA | NA | NA | NA | NA | NA | NA | NA | NA | NA | NA | NA | NA | NA | NA | NA | NA |
| L2 | 0.004 | 0.003 | 0.005 | 0.003 | 0.003 | 0.004 | 0.002 | 0.004 | 0.006 | NA | NA | NA | NA | NA | NA | NA | NA | NA | NA | NA | NA | NA | NA | NA | NA |
| N4 | 0.002 | 0.002 | 0.004 | 0.002 | 0.002 | 0.003 | 0.001 | 0.001 | 0.005 | 0.002 | NA | NA | NA | NA | NA | NA | NA | NA | NA | NA | NA | NA | NA | NA | NA |
| L1 | 0.002 | 0.002 | 0.003 | 0.001 | 0.002 | 0.003 | 0.002 | 0.003 | 0.007 | 0.001 | 0.001 | NA | NA | NA | NA | NA | NA | NA | NA | NA | NA | NA | NA | NA | NA |
| L12 | 0.003 | 0.003 | 0.000 | 0.003 | 0.003 | 0.003 | 0.007 | 0.008 | 0.001 | 0.007 | 0.006 | 0.005 | NA | NA | NA | NA | NA | NA | NA | NA | NA | NA | NA | NA | NA |
| L14 | 0.002 | 0.002 | 0.000 | 0.002 | 0.003 | 0.002 | 0.006 | 0.008 | 0.001 | 0.006 | 0.005 | 0.003 | 0.000 | NA | NA | NA | NA | NA | NA | NA | NA | NA | NA | NA | NA |
| L6 | 0.003 | 0.003 | 0.006 | 0.003 | 0.004 | 0.004 | 0.003 | 0.004 | 0.006 | 0.004 | 0.003 | 0.002 | 0.006 | 0.006 | NA | NA | NA | NA | NA | NA | NA | NA | NA | NA | NA |
| L7 | 0.003 | 0.002 | 0.005 | 0.002 | 0.003 | 0.004 | 0.002 | 0.002 | 0.006 | 0.002 | 0.002 | 0.000 | 0.006 | 0.005 | 0.002 | NA | NA | NA | NA | NA | NA | NA | NA | NA | NA |
| L8 | 0.002 | 0.002 | 0.004 | 0.001 | 0.002 | 0.002 | 0.001 | 0.002 | 0.004 | 0.003 | 0.001 | 0.002 | 0.005 | 0.005 | 0.002 | 0.001 | NA | NA | NA | NA | NA | NA | NA | NA | NA |
| L9 | 0.002 | 0.001 | 0.003 | 0.001 | 0.001 | 0.002 | 0.001 | 0.002 | 0.004 | 0.002 | 0.001 | 0.000 | 0.005 | 0.004 | 0.002 | 0.001 | 0.000 | NA | NA | NA | NA | NA | NA | NA | NA |
| L4 | 0.004 | 0.003 | 0.005 | 0.002 | 0.003 | 0.004 | 0.003 | 0.004 | 0.007 | 0.000 | 0.002 | 0.001 | 0.007 | 0.005 | 0.004 | 0.002 | 0.003 | 0.002 | NA | NA | NA | NA | NA | NA | NA |
| S2 | 0.002 | 0.001 | 0.000 | 0.002 | 0.002 | 0.001 | 0.005 | 0.007 | 0.000 | 0.005 | 0.004 | 0.003 | 0.000 | 0.000 | 0.005 | 0.005 | 0.004 | 0.003 | 0.005 | NA | NA | NA | NA | NA | NA |
| L5 | 0.002 | 0.002 | 0.003 | 0.001 | 0.002 | 0.002 | 0.001 | 0.002 | 0.004 | 0.002 | 0.001 | 0.001 | 0.005 | 0.004 | 0.002 | 0.001 | 0.000 | 0.000 | 0.002 | 0.003 | NA | NA | NA | NA | NA |
| L15 | 0.004 | 0.004 | 0.006 | 0.003 | 0.005 | 0.005 | 0.001 | 0.000 | 0.007 | 0.003 | 0.001 | 0.002 | 0.008 | 0.007 | 0.003 | 0.002 | 0.002 | 0.002 | 0.003 | 0.006 | 0.002 | NA | NA | NA | NA |
| S5 | 0.001 | 0.000 | 0.002 | 0.001 | 0.000 | 0.001 | 0.003 | 0.004 | 0.003 | 0.003 | 0.002 | 0.002 | 0.004 | 0.003 | 0.004 | 0.003 | 0.002 | 0.002 | 0.004 | 0.002 | 0.002 | 0.004 | NA | NA | NA |
| K1 | 0.007 | 0.006 | 0.008 | 0.005 | 0.006 | 0.007 | 0.004 | 0.003 | 0.009 | 0.002 | 0.002 | 0.003 | 0.010 | 0.009 | 0.006 | 0.004 | 0.005 | 0.004 | 0.002 | 0.008 | 0.004 | 0.003 | 0.006 | NA | NA |
| S8 | 0.004 | 0.003 | 0.005 | 0.002 | 0.003 | 0.004 | 0.001 | 0.001 | 0.006 | 0.002 | 0.000 | 0.002 | 0.006 | 0.003 | 0.002 | 0.002 | 0.003 | 0.005 | 0.001 | 0.001 | 0.004 | 0.001 | 0.000 | 0.007 | NA |

**Table S14**. Temperature and chlorophyll concentration in the 19 beach spawning sites in the NWA lineage.

| Sampling  site | Temperature  (T°C.) | Chlorophyll concentration (mg.m-3) |
| --- | --- | --- |
| N2 | 5.52039623260498 | 0.688368320465088 |
| L13 | 1.12419378757477 | 0.478062391281128 |
| S4 | 4.74665212631226 | 1.37381112575531 |
| S6 | 3.89567613601685 | 2.68240904808044 |
| S3 | 2.09210324287415 | 0.0834950134158134 |
| L10 | 3.82797312736511 | 0.634994804859161 |
| S1 | 2.42955923080444 | 0.299261420965195 |
| L2 | 1.4544506072998 | 0.617295384407043 |
| N4 | 0.137531459331512 | 0.0966810435056686 |
| L12 | 2.86810970306396 | 0.820952594280243 |
| L14 | 5.97181272506714 | 0.493824124336243 |
| L6 | 2.15750002861023 | 0.676358282566071 |
| L8 | 2.54782342910767 | 0.729797780513763 |
| S2 | 5.75103664398193 | 1.09577214717865 |
| L5 | 1.74898135662079 | 0.719707727432251 |
| L15 | 1.12419378757477 | 0.478062391281128 |
| S8 | 4.07262229919434 | 1.63975191116333 |
| S5 | 4.80997228622437 | 1.27641952037811 |
| K1 | 0.477539420127869 | 0.537216484546661 |
